# Towards an understanding of the taxonomy of the Eurasian montane riparian genus *Myricaria* (Tamaricaceae) through Angiosperms353 herbariomics

**DOI:** 10.64898/2026.10.03.751413

**Authors:** Kailin Weitkämper, Elena Lyakh, Diego F. Morales-Briones, Felix Weitkämper, Silke Werth

**Affiliations:** Fakultät für Biologie, Ludwig-Maximilians-Universität München, Menzinger Str. 67, 80638 München, Germany; Central Siberian Botanical Garden of the Siberian Branch of the RAS, 101, Zolotodolinskaya Str., 630090 Novosibirsk, Russia; Research Center for Artificial Intelligence, German University of Digital Science, Marlene-Dietrich-Allee 14, 14482 Potsdam, Germany; Institut für Informatik, Ludwig-Maximilians-Universität München, Oettingenstraße 67, 80538 München, Germany

**Keywords:** *Myricaria*, Angiosperms353, taxonomy, phylogenomics, herbariomics

## Abstract

*Myricaria* Desv. (Tamaricaceae) is a genus of shrubs that play a keystone role in montane riparian ecosystems across Eurasia, shaping riverine habitats and supporting ecological function. The genus has a centre of origin in the Qinghai–Tibetan Plateau and exhibits a broad distribution, but many species boundaries are unresolved. Here, we present the first phylogenomic analysis of *Myricaria* using the Angiosperms353 target enrichment bait set, with sampling across its full geographic range. Off-target plastid loci were extracted from next-generation sequencing data using the CAPTUS bioinformatics pipeline. We infer species relationships using ASTRAL-IV and network analysis with SplitsTree, in conjunction with morphology. We also provide a taxonomic treatment of the genus, integrating our morphological observations and phylogenomic analyses for species delimitations. Based on our analyses, recognise two infrageneric sections: sect. *Parallelantherae* (containing only *M. elegans*) and sect. *Myricaria* (the remainder of the genus, with type species *M. germanica*). We also propose synonymising *M. davurica*, *M. bracteata* and *M. squamosa* with *M. germanica*. Although species boundaries in the Asian highlands remain unresolved, we conclude that the species *M. elegans*, *M. germanica*, *M. laxiflora*, *M. longifolia*, *M. platyphylla*, and *M. pulcherrima* are well-supported both phylogenomically and morphologically.

## INTRODUCTION

*Myricaria* Desv. (Tamaricaceae) is a genus of woody shrubs occurring in riparian montane habitats across Eurasia (Bobrov, 1967; Webb, 1968; Wu & al., 2007; GBIF 2026). The genus has its origin in the Qinghai-Tibetan Plateau (QTP) (Liu & al., 2006; Wang & al., 2009; Zhang & al., 2014; Sun & Werth, 2024), from whence it spread west to Europe via mountain ranges in Central Asia, as well as spreading south to central China and western and eastern Siberia (Sun & Werth, 2024). The current Eurasian distribution according to GBIF occurrences (GBIF 2026) is illustrated as cyan dots in Fig. 1.

**Figure 1:**
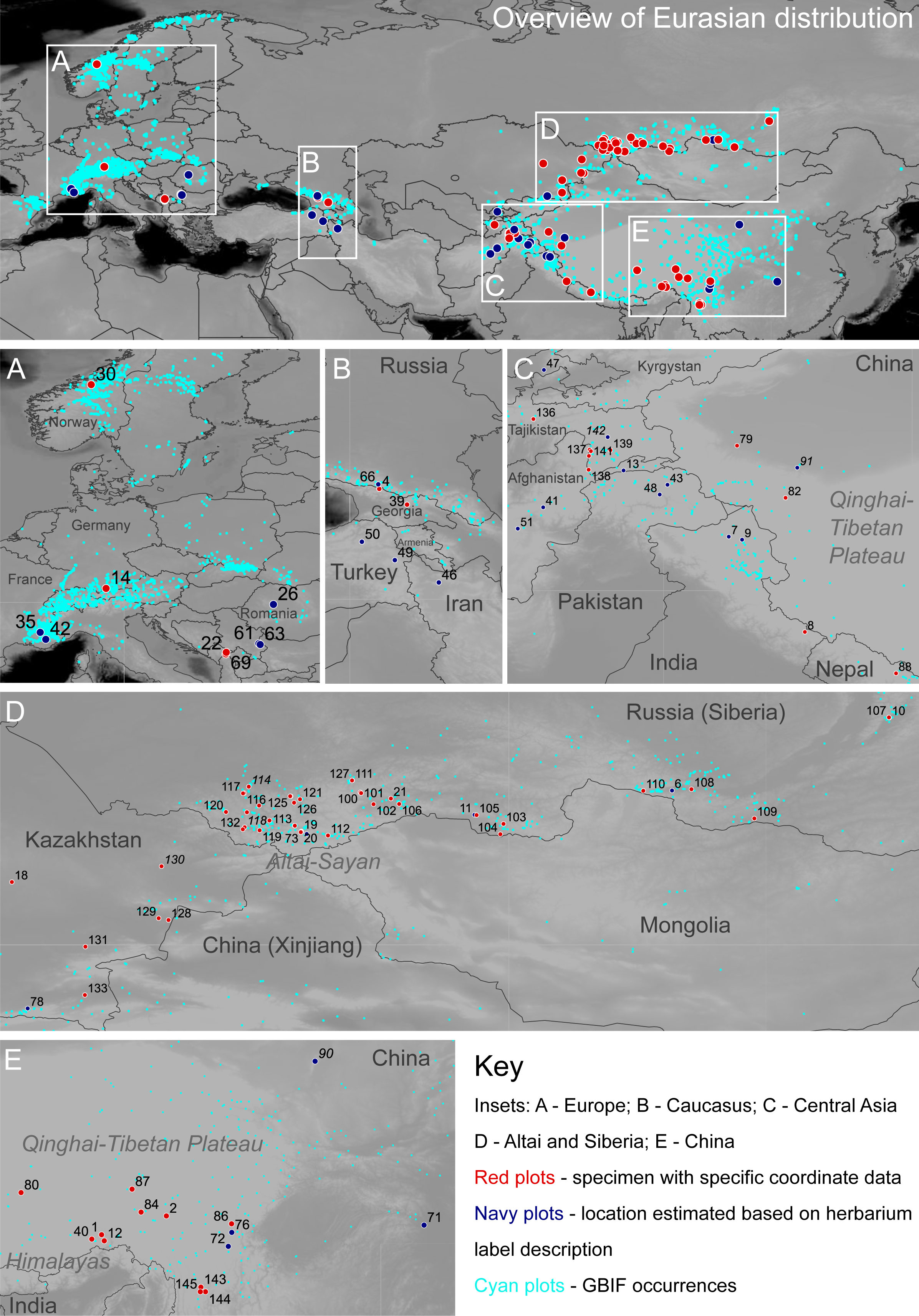
Maps illustrating the distribution of the genus *Myricaria* taken from GBIF (2026), as well as provenance locations of all herbarium specimens with location data. Specimen numbers correspond to nuclear orthologue species tree (Fig. 3), SplitsTree analyses (Fig. 4; suppl. Fig. S4), plastid trees (Fig. 5; suppl. Fig. S5), photographs (Fig. 6-10), and lists of specimens sampled (suppl. Table S1; Appendix S4). Specimen numbers in italics denote that genetic data was not included in nuclear orthologue species tree (Fig. 3) due to insufficient quality.

Species like *Myricaria germanica* (L.) Desv. serve as indicator taxa of undisturbed rivers and function as keystone species in gravel-bed river ecosystems (Harzer & al., 2018). Unfortunately, due to anthropogenic activity such as dam building, concrete river banks, and river straightening (Sitzia & al., 2021), *Myricaria* habitats have been damaged throughout the range of the genus. For example, the controversial construction of the Three Gorges Dam in Hubei Province nearly rendered *Myricaria laxiflora* (Franch.) P.Y.Zhang & Y.J.Zhang extinct in the wild (Liu & al., 2009), as well as affecting the hydrochory of all riparian plants in this section of the Yangtze River system (Liu & al., 2006). In Europe, dams have also had a large disruptive effect on the gene flow of *M. germanica* populations (Werth & Scheidegger, 2014; Werth & al., 2014).

Despite its ecological and biogeographic significance, the genus remains taxonomically unresolved. First described by Desvaux (1825), *Myricaria* was split from *Tamarix* L. based on floral structure. Ehrenberg (1827) noted the problem of unclear species delineations in this genus, and this issue appears to persist. The difficulties with taxonomy reflect those of its sister genus *Tamarix*, which is “one of the taxonomically most complex genera among the angiosperms” (Villar & al., 2019). To give an idea of the size of the genus, and to give a brief overview of differing opinions on species delimitations, Table 1 provides a list of validly published names with citations from the accepting authority. This table provides a summary from the most recent floristic treatments or taxonomic studies covering each name; the most recent study of *Myricaria,* which covers its entire distribution, is Bobrov (1967), but more recent floristic treatments are available covering local geographies, such as the Flora of China (Wu & al., 2007). A more detailed history of the names can be found in the section Taxonomic Treatment.

**Table 1:** List of validly published species names, authorities which accept these names, and authorities which synonymise names under a different name.

| Validly published names | Authorities that accept name | Authorities that synonymise name under a different name | Notes |
| --- | --- | --- | --- |
| **Species accepted within *Myricaria*** |  |  |  |
| *Myricaria albiflora* Grierson & D.G.Long | Grierson (1982); Mao & Dash (2020) |  |  |
| *Myricaria alopecuroides* Schrenk ex Fisch. & C.A.Mey. |  | Bobrov (1967) – *M. bracteata*; Lyakh (2008) in Siberia – *M. bracteata*; Gangurde & al. (2026) – *M. bracteata*; Qaiser (1982) – *M. germanica*; Zhao & al. (2019); – *M. bracteata*; Dar & Govaerts (2025) – *M. bracteata* |  |
| *Myricaria armena* Boiss. |  | Qaiser (1982) – *M. squamosa*; Mao & Dash (2020) – *M. squamosa* |  |
| *Myricaria bracteata* Royle | Bobrov (1967); Wu & al. (2007); Lyakh (2008); Tsarenko & Peregrym (2013) in Ukraine; Zhao & al. (2019); Mao & Dash (2020); Dar & Govaerts (2025); Gangurde & al. (2026) | Qaiser (1982) – *M. germanica* |  |
| *Myricaria brevifolia* Turcz. |  | Bobrov (1967) – *M. davurica*; Qaiser (1982) – *M. davurica* |  |
| *Myricaria davurica* (Willd.) Ehrenb. (orthographical variation “*Myricaria dahurica*”) | Bobrov (1967); Qaiser (1982); Gangurde & al. (2026) | Lyakh (2008) – *M. longifolia* | Wu & al. (2007): “further study is needed to ascertain ... if this species actually occurs in China” |
| *Myricaria elegans* Royle | Bobrov (1967); Wu & al. (2007) | Baum (1978) – *Tamarix ladachensis* B.R.Baum; Ovchinnikov & Kinzikaeva (1977) – *Myrtama elegans* (Royle) Ovcz. & Kinzik.; Qaiser (1982) – *Tamaricaria elegans* (Royle) Qaiser | Accepted as well-supported species, but disagreement on whether it is part of *Myricaria* |
| *Myricaria germanica* (L.) Desv. | Bobrov (1967); Qaiser (1982); Webb (1968); Tsarenko & Peregrym (2013) in Ukraine; Sitzia & al. (2021) literature review |  |  |
| *Myricaria hedinii* Paulsen |  | Bobrov (1967) – *M. prostrata*; Hara (1972) – *M. prostrata*; Qaiser (1982) – *M. prostrata*; Wu & al. (2007) – *M. prostrata* |  |
| *Myricaria herbacea* Desv. |  | Bobrov (1967) – does not explicitly mention which species, but possibly *M. germanica* |  |
| *Myricaria hoffmeisteri* Klotzsch |  | Bobrov (1967) – *M. squamosa*; Qaiser (1982) – *M. squamosa* |  |
| *Myricaria laxiflora* (Franch.) P.Y.Zhang & Y.J.Zhang | Wu & al. (2007) |  |  |
| *Myricaria linearifolia* Desv. |  | Bobrov (1967) – *M. longifolia* |  |
| *Myricaria longifolia* (Willd.) Ehrenb. | Bobrov (1967); Lyakh (2008) |  |  |
| *Myricaria macrostachya* Kar. & Kir. |  | Bobrov (1967) – *M. bracteata*; Qaiser (1982) – *M. germanica* |  |
| *Myricaria paniculata* P.Y.Zhang & Y.J.Zhang | Wu & al. (2007) |  |  |
| *Myricaria platyphylla* Maxim. | Bobrov (1967); Wu & al. (2007); Zhao & al. (2019) |  |  |
| *Myricaria prostrata* Hook.f. & Thomson | Bobrov (1967); Qaiser (1982); Wu & al. (2007); Mao & Dash (2020) |  |  |
| *Myricaria pulcherrima* Batalin | Bobrov (1967); Wu & al. (2007) |  |  |
| *Myricaria rosea* W.W.Sm. | Wu & al. (2007); Mao & Dash (2020) | Bobrov (1967) – *M. squamosa* |  |
| *Myricaria schartii* Vassilcz. |  | Bobrov (1967) – *M. bracteata*; Qaiser (1982) – *M. germanica* |  |
| *Myricaria squamosa* Desv. | Bobrov (1967); Qaiser (1982); Wu & al. (2007); Mao & Dash (2020); Rajbhandari & al. (2021) | Webb (1968) – *M. germanica*; Lyakh (2008) in Siberia – *M. bracteata* |  |
| *Myricaria wardii* C.Marquand | Bobrov (1967); Wu & al. (2007); Rajbhandari & al. (2021) |  |  |
| **Species not currently accepted within *Myricaria*** |  |  |  |
| *Myricaria vaginata* Desv. |  | Mao & Dash (2020) – *Tamarix ericoides* Rottler & Willd. | Currently assigned to genus *Tamarix* rather than *Myricaria*; not included in study |

Species of the genus *Myricaria* are often found in the border regions of countries because many mountain regions and rivers mark territorial boundaries. Due to its presence in many countries and its ecological importance, this genus has been widely studied around the world. We observe, however, that modern authors tend to focus on studying *Myricaria* in their local regions rather than include the whole distribution (for example, in China, Liu & al. (2009), Wang & al. (2009), Zhang & al. (2014), and Hu & al. (2023); in Russia, Lyakh (2013), in the European Alps, Werth & Scheidegger (2014) and Werth & al. (2014); in the Balkans, Lakušić & Pavlović (1971) and Lakušić & al. (1974)).

### Areas of taxonomic confusion

#### Myricaria, Myrtama, or Tamaricaria?

Royle (1835) first published the species *Myricaria elegans* Royle, which has large leaves and unfused stamens, and was found in Kashmir. Subsequently, Baum (1978) transferred this species to *Tamarix*, with the new name *Tamarix ladachensis* B.R.Baum (the name *Tamarix elegans* was already occupied by a different species), on the basis that this is the only species within *Myricaria* that has free stamens. Bobrov (1967) reinstated the original name *Myricaria elegans* Royle. Ovchinnikov & Kinzikaeva (1977) designated this species *Myrtama elegans* (Royle) Ovcz. & Kinzik, placing it into the monospecific genus *Myrtama* Ovcz. & Kinzik. On the basis of the presence of unfused stamens, and also from the morphology of the pollen grains, as well as the absence of anthocyanin, Qaiser & Ali (1978) also support placing this species in a monospecific genus, proposing the name *Tamaricaria elegans* (Royle) Qaiser & Ali. However, since the name *Myrtama elegans* was published earlier, this name has priority, so *Tamaricaria* would be a homotypic synonym of *Myrtama*. Gaskin & al. (2004) also find support for *Myrtama* as a separate genus, as it is placed as sister to both *Tamarix* and *Myricaria* in their molecular phylogeny.

Currently, there is still no consensus among modern authors on the generic classification. For example, *Myrtama elegans* is listed as accepted on the Plants of the World Online database (POWO, 2026), while *Myricaria elegans* is accepted on World Flora Online (WFO, 2026). Many recent authors still use the name *Myricaria elegans* in their publications, implicitly considering this species to be part of the genus *Myricaria* (Liu & al., 2009; Wang & al., 2009; Zhang & al., 2014; Hu & al., 2023). In the context of this contribution, we refer to this species as *Myricaria elegans*, thereby treating it as part of the genus.

#### *Myricaria* in the Asian Highlands

Several accepted species’ type specimens have been collected in the Asian Highlands regions (including the Himalayas, the Pamirs, the Hindukush, and the Karakorum), such as *M. elegans* (discussed in section *Myricaria, Myrtama*, or *Tamaricaria*?), *Myricaria albiflora* Grierson & D.G.Long, *Myricaria prostrata* Hook.f. & Thomson, *Myricaria rosea* W.W.Sm., and *Myricaria wardii* C.Marquand. Although *M. albiflora* is seen by some sources as an accepted name (Mao & Dash, 2020; POWO, 2026), there are few sources citing this name.

#### The “*Myricaria squamosa* species complex”

The species *Myricaria squamosa* Desv. is particularly mysterious. This name was first published by Desvaux (1825), with distinguishing features being listed, including dense flowers, imbricate scales, and subulate-oblong leaves. Neither Desvaux (1825) nor the holotype, held at P (barcode P01902095), provide a clear provenance location (the herbarium sheet reads “ex Oriente”). This specimen has imbricate scales, small leaves of around 3–5 mm, and racemes that are 2–3 cm long, and shares its herbarium sheet with an unlabelled specimen.

Perhaps due to the ambiguity of the type specimen and its collection locality, authors have contradictory ideas about its species delimitation. For example, Bobrov (1967) describes *M. squamosa* as being similar in morphology to *Myricaria longifolia* (Willd.) Ehrenb. (leaves 0.8–1.5 cm long, 1–1.5 mm wide), with *M. squamosa* differing only by its leaf shape (broadly elliptic or subovate, 0.6–1 cm long). According to the Flora Europaea (Webb, 1968), *M. squamosa* is a synonym of *M. germanica*, erroneously applied to specimens that show imbricate scales. The Flora of China (Wu & al., 2007) describes *M. squamosa* as having lateral or axillary inflorescences, as well as many persistent imbricate scales.

Phylogenetic analyses with Sanger Sequencing methods, such as studies by Liu & al. (2009), Wang & al. (2009), and Zhang & al. (2014), also group the putative species *Myricaria bracteata* Royle, *Myricaria paniculata* P.Y.Zhang & Y.J.Zhang, *Myricaria alopecuroides* Schrenk ex Fisch. & C.A.Mey. and *M. squamosa* in a single unresolved clade. Recently, a molecular phylogeny by Hu & al. (2023) using whole chloroplast genomes showed that even with significantly more base pairs, there are still polytomies in the phylogenies produced; the authors call this the “*M. squamosa* species complex”, which includes other specimens diagnosed as *M. bracteata* and *M. paniculata*. Taken together, these studies suggest that morphological characters traditionally used for species delimitation in this group are insufficient and potentially misleading.

#### *Myricaria* in the Altai Mountains and Siberia

Authors disagree as to how many species and which species are found in Siberia and its surrounding regions. Two type specimens, *M. longifolia* and *Myricaria davurica* (Willd.) Ehrenb., were collected from this geographical area; these were described by Willdenow (1816) in the same publication, as *Tamarix longifolia* Willd. and *Tamarix davurica* Willd., respectively. Ehrenberg (1827) transferred these species to *Myricaria*. The Flora of the U.S.S.R. (Gorshkova, 1974) describes four species in this area, namely, *Myricaria alopecuroides*, *M. davurica*, *M. longifolia*, and *M. squamosa*. Bobrov (1967) also treats *M. davurica* (leaves 0.6–1 cm long; petals 1.5 mm wide) and *M. longifolia* as separate species with a coinciding range in southern Siberia; the Altai; southern Transbaikalia; and northern Mongolia, noting that they are similar but clearly distinguishable. However, Bobrov (1967) notes that *M. squamosa* is not found in this region, but only in Central Asia, including Gorno-Badakhshan, the Pamirs, the central Tien Shan, Gilgit, Ladakh, Lahul, Punjab, and the Qaidam Basin. In addition, Bobrov (1967) considers *M. alopecuroides* to be a synonym of *M. bracteata*.

In-situ morphological studies of Siberian *Myricaria* populations (Lyakh, 2013) suggest that there are only two species in the region – which Lyakh (2013) refers to as *M. bracteata* and *M. longifolia* - and that Siberian specimens previously diagnosed as *M. alopecuroides*, *M. squamosa* or *M. bracteata* are in fact the same species, while *M. davurica* (orthographical variation *M. dahurica*) is a synonym of *M. longifolia*. Sun & Werth (2024) also demonstrated that specimens from central Siberia, Transbaikalia and the Altai-Sayan Mountains, diagnosed as *Myricaria longifolia* according to Lyakh (2013), form a well-supported clade. However, the clade’s position in the phylogenetic tree was not clear, with the plastid tree showing a different topology to that of the nuclear ITS tree.

#### *Myricaria* in Europe to Iran and surrounding areas

In western Europe, most authors accept that *M. germanica* is the only species (Webb, 1968; Sitzia & al. 2021; Villar & Alonso, 2017). However, in the Balkans, Lakušić & Pavlović (1971) put forward a new species, *Myricaria ernesti-mayeri*, as separate from *M. germanica*, on the basis of the populations in the Balkans and in Austria being associated with differing species of *Salix* L. and *Alnus* Mill., as well as variations in the morphometric measurements and xerophytic characteristics between these two populations (Lakušić & al., 1974). However, the name *M. ernesti-mayeri* has not been validly published, as its publication does not provide sufficient details to designate a type specimen (no collector, date or specific location is cited).

Some authors believe that *M. bracteata* and *M. squamosa* are also present in eastern Europe. For example, according to Tsarenko & Peregrym (2013), there are three species of *Myricaria* in Ukraine, namely *M. germanica*, *M. bracteata*, and *M. squamosa,* and these can be differentiated by their bract shapes. According to Schiman-Czeika (1964), *M. germanica* has a range from Scandinavia and the Pyrenees to the Himalayas, while *M. squamosa* has a range from the Crimea and the Caucasus to the Himalayas and China. The Euro+Med PlantBase database (Villar & Alonso, 2017, accessed 17 April 2026) also acknowledges the presence of *M. bracteata* in the Crimea and the Caucasus, and *M. squamosa* in the Crimea, Turkey, and the Caucasus region. It is not clear how the “*M. squamosa* species complex” mentioned by Hu & al. (2023) relates to European specimens, because no European specimens were included in that study. The biogeographical study by Sun & Werth (2024) is so far the only study to have included a large sample of European specimens as well as Asian specimens. Their phylogenetic analysis found that European (west of the Caucasus) *Myricaria* specimens were monophyletic, suggesting that there was only one migration of *Myricaria* to Europe.

### Infrageneric sections and series

Authors have proposed grouping *Myricaria* into various sections. Niedenzu (1895) suggested two sections, *Parallelantherae* containing *M. elegans* only, and *Renantherae*, encompassing all remaining species in the genus. The sections described by Niedenzu (1895) were also used by Gorshkova (1974). Bobrov (1967), who accepted 10 species, suggested dividing the genus into four series:

1. Germanicae: *M. germanica*, *M. bracteata*, *M. prostrata*, *M. wardii*;
2. Dahuricae: *M. squamosa*, *M. davurica*;
3. Elegantes: *M. elegans*, *M. longifolia*; and
4. Platyphyllae: *M. platyphylla*, *M. pulcherrima*.

More recently, based on their molecular phylogeny, Zhang & al. (2014) suggest two new sections in addition to those of Niedenzu (1895), adding the section Alpinae, consisting of *M. prostrata*, *M. rosea*, and *M. wardii*, and the section *Laxiflorae*, containing only *M. laxiflora*. These new section divisions reflect the aforementioned geographic distributions, with the Alpinae representing species from the Himalayas, while *Laxiflorae* consists of the only species found in subtropical central China. While the infrageneric classifications of Zhang & al. (2014) are based on a molecular phylogeny from Sanger sequencing methods, these classifications have not been evaluated in a comprehensive phylogenomic framework including specimens from across the full range of *Myricaria*. Since Zhang & al. (2014) did not designate a type, the section name Alpinae has not been validly published.

### GENOMIC TOOLS FOR PHYLOGENETIC INFERENCE

The development of Next Generation Sequencing (NGS) methods has much improved the efficiency and output of phylogenetic studies, in particular, the development of the Angiosperms353 probe set (Johnson & al., 2018), a “universal” kit which is designed to capture 353 nuclear genes across angiosperms (Baker & al., 2021b). Since then, the Angiosperms353 bait set has been used to explore phylogenetic relationships from the long macroevolutionary scale, like the PAFTOL project (Baker & al., 2021a), to the population level (Slimp & al., 2021).

To our knowledge, the present study is the first genus-wide phylogenomic study of *Myricaria* using targeted nuclear as well as chloroplast loci. However, phylogenomic analyses with the Angiosperms353 bait set, along with morphological studies and other analyses, have provided important insights into the evolutionary history of other genera. Recent examples include an integrative study on *Kirganelia* Juss. (Phyllanthaceae), using phylogenomic analyses with target capture and population genetics methods to aid species delimitation (Putri & al., 2026). Another recent phylogenomic study with Angiosperms353, focusing on *Orostachys* Fisch. (Crassulaceae), investigated the infrageneric relationships within this complex genus (Lee & al., 2025). Rasaminirina & al. (2026) also used Angiosperms353 in conjunction with morphology to study the genus *Bulbostylis* (Cyperaceae), concluding that characteristics such as leaf sheath morphology, inflorescence structure, and glume morphology are reliable for species identification.

A great benefit of the Angiosperms353 bait set is that it works well with herbarium specimens. The 353 target nuclear genes have been successfully extracted from very old or degraded herbarium specimens (Hart & al., 2016; Brewer & al., 2019). The wide Eurasian distribution of *Myricaria* means that herbarium specimens are a particularly useful source of genetic material for sampling. This is very useful for studying *Myricaria*, because of its extensive range, much of which is geographically inaccessible due to the terrain. The genus’ habitat in riparian montane regions is also frequently on territorial borders, which increases the difficulty of collecting fresh specimens, as these areas are more likely to be politically unstable. In this study, we take advantage of the benefits of the Angiosperms353 bait set to construct a phylogeny of *Myricaria* covering the full Eurasian range.

### AIMS

We aim to build a robust phylogeny of *Myricaria* and to assess gene discordance, building on recent work clarifying the biogeographic history of *Myricaria* (Sun & Werth, 2024), which elucidated the migration patterns of the genus from the QTP to its current distribution, including western Europe. In addition, we test currently accepted species delineations, as well as the validity of infrageneric sections proposed by previous authors. Overall, this study aims to:

1. reconstruct a phylogeny of *Myricaria* nuclear genes, using the Angiosperms353 bait set;
2. assess the congruence between molecular lineages and current species delimitations;
3. interpret taxonomic implications based on phylogenomic and morphological evidence; and
4. provide an up-to-date summary of the genus, including details of type specimens and synonyms.

## MATERIALS AND METHODS

### TAXON SAMPLING

Overall, 98 *Myricaria* specimens from across the native Eurasian range of this genus have been selected for this study. We aimed to include specimens representing as many of the putative species within the genus and covering as much of the geographic range as possible. These specimens have been sourced from a range of herbaria and botanical gardens from around the world, including A, BEOU, BRNU, GH, GOET, HIB, M, MB, MSB, NS, NSK, P, TAIF, UPS. An overview of all the herbarium specimens sequenced can be found in suppl. Appendix S1. Details including specimen number, species identification, geographical provenance, date, collector, and herbarium of all specimens sampled can be found in suppl. Table S1. Some of the herbarium specimens sampled in this study have also been sequenced by Sun & Werth (2024) using Sanger sequencing, where they are assigned the same specimen numbers.

### NEXT GENERATION SEQUENCING WITH THE ANGIOSPERMS353 BAIT SET

#### DNA extraction

A modified CTAB extraction protocol (Doyle and Doyle, 1987) was used to extract DNA from the selected herbarium specimens. The following modifications to the protocol were made to maximise yield. For each herbarium specimen, one to two branchlets were ground using a TissueLyser II (Qiagen, Venlo, Netherlands) bead mill, then 8 µL of β-mercaptoethanol was added to each sample in addition to 500 µL CTAB before incubation (Doyle and Doyle (1987), steps 1 and 2). Extraction with 500 µL chloroform-isoamyl alcohol (24:1) was performed twice (steps 3 and 4). Upon adding chilled 300 µL 70% isopropanol (step 5), samples were left overnight at -20 °C to maximise DNA precipitation. The supernatant was then removed, and the resulting DNA pellet redissolved in DNA-free purified water. After extraction, DNA concentration was measured with the dsDNA HS DNA Assay Kit on a Qubit 4 Fluorometer (Thermo Fisher Scientific, Waltham, Massachusetts, USA). Agarose gel electrophoresis with 1% gels stained with ethidium bromide was run to assess the DNA fragment lengths. In most cases, samples had short fragment lengths (gel streaks were mostly under 500 bp), so sonication was not necessary, but those that showed longer fragment lengths (just six samples) were sonicated for 60 s or 80 s in a 50 µL Covaris microTUBE using a Covaris M220 sonicator (Covaris, Woburn, MA, USA). The settings used were peak power at 75.0 W, duty cycle at 10.0% and cycles/burst at 200 (Covaris, 2024). DNA purity was assessed using a NanoDrop spectrophotometer (Thermo Fisher Scientific). Where quality was low, DNA extracts were cleaned with SPRISelect or AMPure XP beads (both by Beckman Coulter, Brea, CA, USA) with a DNA extract to bead volume ratio of 1:1.8.

#### Library preparation and pooling

DNA libraries were made with the NEBNext Ultra II DNA Library Preparation Kit (New England Biolabs, Ipswich, MA, USA), following the manufacturer’s protocol (New England Biolabs, 2022) at half volume to reduce costs. The adaptor was diluted one part in ten in nuclease-free water. To size-select adaptor-ligated DNA, 12.5 µL SPRISelect beads were added at step 3A.2 of the protocol, and 22.5 µL beads were added at step 3A.6. Library concentration was measured with a Qubit 4 Fluorometer and average fragment size checked on a TapeStation with HS D1000 Screen Tape (Agilent, Santa Clara, CA, USA). Libraries were normalised by molarity and then pooled by average fragment size into four pools containing 22 to 24 libraries. The pools were dehydrated and eluted in 7.5 µL nuclease-free water and contained roughly 1 µg DNA each.

#### Target capture and Illumina sequencing

Target capture was carried out following the Standard Plants myBaits protocol v.5.2 (Daicel Arbor Biosciences, 2022) for the Angiosperms353 bait set, with the hybridisation and wash temperature set to 60 °C. The hybridisation reaction (Part 1) was incubated for 20 hours at 60 °C (same as wash temperature). The final PCR amplification (Part 3) was performed using NEBNext Ultra II Q5 Polymerase (New England Biolabs) with myBaits beads (Daicel Arbor Biosciences, Ann Arbor, MI, USA) in solution, for 14 cycles. The resulting target-enriched library pools were cleaned using the NucleoSpin Gel and PCR Clean-up kit (Macherey-Nagel, Düren, Germany) following protocol 5.1 PCR Cleanup (Macherey-Nagel, 2017). The four pools were again quantified using Qubit and TapeStation, then pooled together for two final pools with a molarity of roughly 7 nM at 30 µL each. The final pooled libraries were sequenced by the Core Genomics Facility of the Helmholtz Zentrum Munich, Germany, on Illumina NextSeq 1000 (Illumina, San Diego, CA, USA) with 150 bp paired-end reads on two P1.300 flow cells, with an average of 1,983,379 reads per sample or library.

### BIOINFORMATICS

The bioinformatics (Fig. 2) workflow used is based on the pipeline described by Yang & Smith (2014) and Morales-Briones & al. (2022). A more detailed step-by-step process of the entire bioinformatics pipeline (steps A to I) can be found in Appendix S2.

**Figure 2:**
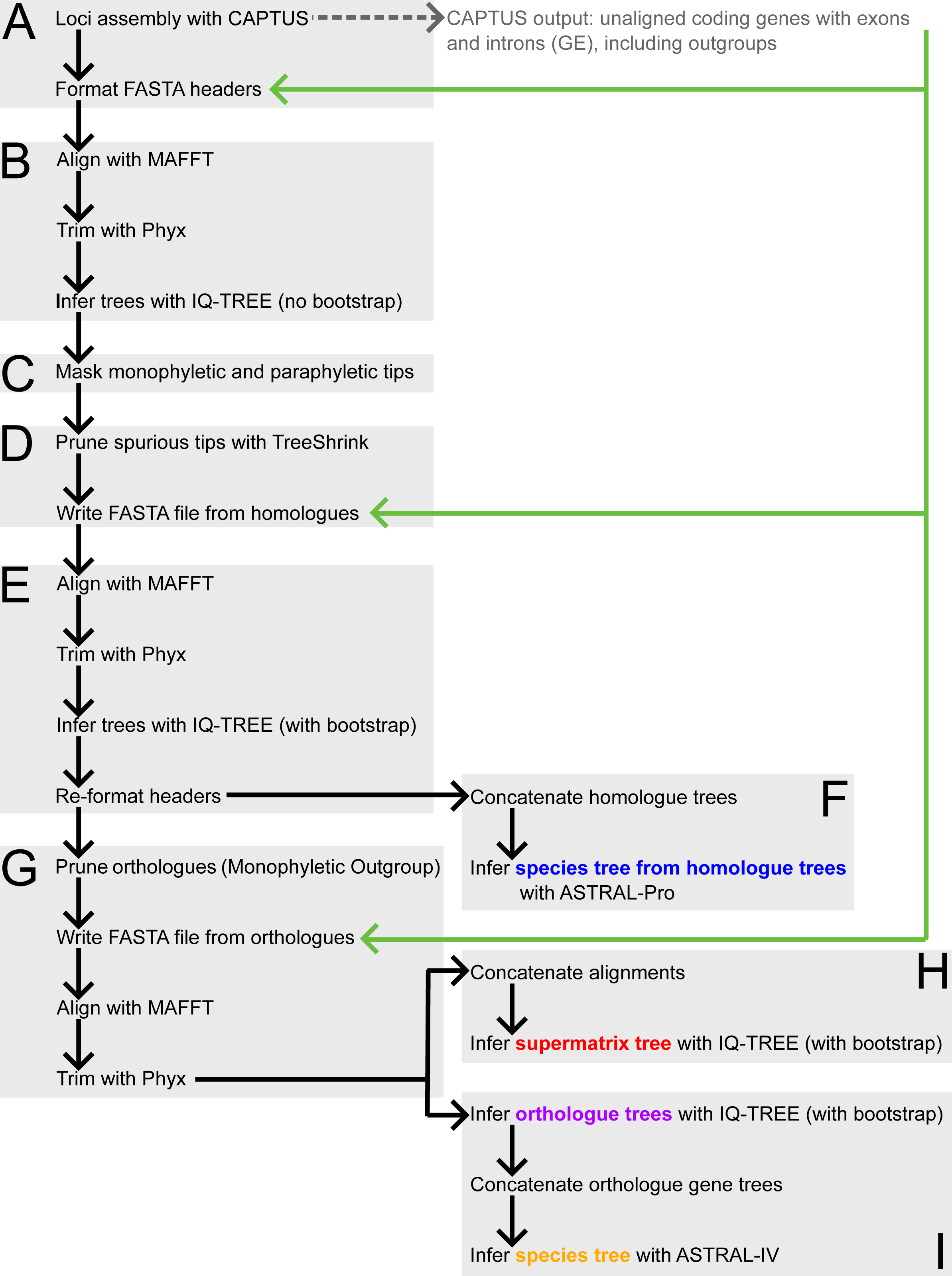
Summary of bioinformatics pipeline.

Scripts (including wrappers and bash scripts used throughout the pipeline) were adapted from https://bitbucket.org/dfmoralesb/target_enrichment_orthology/src/master/ and can be seen in Appendix S3. Sequences and alignments were viewed using SeaView v.5.0.5 (Gouy & al., 2010), and trees were viewed and formatted using FigTree v.1.4.4 (Rambaut, 2018) throughout.

#### Assembly of loci

Raw reads were cleaned, assembled, and the target genes extracted using CAPTUS v.1.0.1 (Ortiz & al., 2026), with the Tamaricaceae sequences available from PAFTOL as a reference (Baker & al., 2021a). The sequences of the species *Tamarix gallica* (Darwin Tree of Life Project, 2024a) and the outgroup *Reaumuria songarica* (Song & al., 2024) were downloaded from GenBank (Sayers & al., 2021) and extracted using the CAPTUS pipeline.

#### Sequence alignment, trimming, and inferring initial trees

The assembled unaligned coding gene sequences with exons and introns (GE), including the outgroup (CAPTUS output), were aligned in MAFFT (Katoh & Standley, 2013). Sequences were then trimmed using Phyx v.1.3.1 (Brown & al., 2017); sites with more than 10% missing data were removed. We then used IQ-TREE v2.3.6 (Minh & al., 2020a) without bootstraps to infer initial gene trees. The best substitution model was found using the ModelFinder function on IQ-TREE (Kalyaanamoorthy & al., 2017).

#### Pruning paralogs and spurious tips

Monophyletic and paraphyletic tips were masked (suppl. Appendices S1, S2) and spurious tips were pruned using TreeShrink v.1.3.9 (Mai & Mirarab, 2018) with quantile 0.10. After pruning, a new FASTA file was written with the remaining sequences from the original unaligned CAPTUS assembly output.

#### Inferring the homologue tree

The resulting FASTA file was again aligned with MAFFT, and sites with more than 30% missing data were trimmed with Phyx. From the resulting alignment, the final homologue trees were inferred using IQ-TREE with 1000 bootstrap replicates. These homologue trees were reformatted and concatenated. A final species tree from homologue trees was then inferred with ASTRAL-Pro3 v.1.19.3.5 (Zhang & al., 2020; Zhang & Mirarab, 2022a; Tabatabaee & al., 2023).

#### Align and trim orthologue sequences

Orthologues were inferred using the tree-based approach Monophyletic Outgroups (MO) following Yang & Smith (2014). The homologue trees were pruned, and a new FASTA file was again written with the remaining sequences from the original unaligned CAPTUS output. These orthologue sequences were again aligned with MAFFT, and sites with more than 30% missing data were trimmed with Phyx.

#### Infer supermatrix tree

The aligned orthologue sequences were concatenated, and from the concatenated orthologue sequences, a supermatrix tree with 1000 bootstrap replicates was inferred with IQ-TREE. To find the best substitution model for the supermatrix tree, we used a partition model on IQ-TREE (Chernomor & al., 2016).

#### SplitsTree analysis

The supermatrix alignment from the Angiosperms353 nuclear genes was subjected to a neighbour-net analysis. This was performed with the SplitsTree App (SplitsTree v.6.4.11; Huson and Bryant, 2024). This analysis generates a split network which represents incompatible or ambiguous signals in the dataset (Huson and Bryant, 2006), where a non-tree-like topology indicates possible discrepancies between the phylogenetic modelling and the dataset. We performed this analysis both including *Reaumuria songarica* (outgroup) and *Tamarix gallica*, and also without.

#### Infer orthologue trees and species tree

From the aligned orthologue sequences, we inferred orthologue gene trees with bootstraps using IQ-TREE. The resulting gene trees were concatenated, and a species tree was inferred from this with ASTRAL-IV v1.19.4.5 (Zhang & al., 2018; Zhang and Mirarab, 2022b; Tabatabaee & al., 2023; Zhang & al., 2025).

#### Gene concordance and site concordance

To assess the gene concordance factor (gCF), the gCF function on IQ-TREE (Minh & al., 2020a) was used. The sCF and gCF values were calculated using the topology of the supermatrix tree and the concatenated orthologue gene trees generated in ASTRAL-IV. The site concordance (sCF) values were also calculated on IQ-TREE (Mo & al., 2022), using the supermatrix tree and the supermatrix alignment as inputs.

While these concordance factors are not strictly support values (Lanfear & Hahn, 2024), they provide some information on the topological variation of the orthologue gene trees that the species tree was inferred from. The gCF values represent what proportion of the gene trees at each node agree with the one shown by the phylogenetic tree (Lanfear & Hahn, 2024). Meanwhile, the sCF values show what proportion of sites (or base pairs in the alignment) agree with the shown tree topology at each node (Minh & al., 2020b; Lanfear & Hahn, 2024). These values are useful proxies for understanding the amount of genetic variation at each node that is characteristic of the subsequent clade.

#### Plastid sequence assembly and tree inference

An advantage of CAPTUS is that it is able to extract plastid loci from off-target reads. Therefore, we were able to run the pipeline shown in Fig. 2 on both the Angiosperms353 sequence dataset as well as on the extra plastid dataset. Off-target cpDNA coding sequences from the same raw Illumina sequencing reads were assembled using CAPTUS. *Tamarix gallica* (Darwin Tree of Life Project, 2024b) and the designated outgroup *Reaumuria songarica* (Yao & al., 2019) were downloaded from GenBank, and coding sequences from these plastomes were also assembled using CAPTUS. We used the plastid sequence data to infer a species tree from orthologue gene trees with ASTRAL-IV. Individuals with below 50% loci retrieval were removed from the analysis, as well as individuals which failed the chi-squared composition test with a *P-value* of 0.01 in IQ-TREE. The code used for plastid sequence assembly and tree inference can be seen in Appendix S2 and Appendix S3.

### MORPHOLOGICAL OBSERVATIONS

Due to well-documented taxonomic inconsistencies within *Myricaria* (see the Areas of taxonomic confusion section), species identification based on existing authorities is difficult. Therefore, we adopted an integrative approach where we used our nuclear orthologue phylogeny in addition to an assessment of morphological variation. Key morphological features were selected based on protologues and subsequent publications, namely habit (erect or prostrate), bracts, leaves, floral parts (sepals, petals, stamens), racemes, and terminal flowers (Bobrov, 1967; Webb, 1968; Gorshkova, 1974; Wu & al., 2007; Tsarenko & Peregrym, 2013), and where available, sampled specimens were assessed for these features. Type specimens were examined to ensure correct assignment of names. Specimen morphology and collection location were compared to the type specimen and the protologue description, and morphological variation evaluated across the clades on our nuclear orthologue tree. This approach enabled us to assess whether putatively diagnostic morphological characters correspond to molecular results.

## RESULTS

### TAXON SAMPLING, DNA EXTRACTION, AND LIBRARY PREPARATION

The maps in Fig. 1 present the provenance locations of 97 (all except one) specimens chosen for this study, where some provenance data were available. Of these 97 specimens, 68 came with specific provenance coordinates, which have been plotted in QGIS Desktop v.3.34.13 (QGIS Development Team, 2023) with red dots. The remaining 29 specimens did not have coordinate data but provided descriptions of the collection location, for example, in relation to towns or geographical features, such that a point could be estimated (shown as navy blue dots). Of the 98 specimens sampled, DNA was successfully extracted from 97 specimens using the CTAB extraction method, and of these, library preparation was successful for 93 individuals. Occurrence data from GBIF were also added (cyan dots) to show the distribution of the genus (GBIF 2026).

### CLEANING OF RAW READS, *DE NOVO* ASSEMBLY AND GENE EXTRACTION

The bioinformatics pipeline CAPTUS (Ortiz & al., 2026) was used to clean raw sequencing reads, as well as to perform *de novo* assembly of reads and extract the Angiosperms353 and plastid genes. All DNA data in this contribution were newly sequenced for this study. The highest number of Angiosperms353 loci extracted for one sample (with the CAPTUS “genes” setting for assembling both introns and exons) was 344 for Samples 14 and 39, while Sample 73 had the lowest recovery among those that were included in the nuclear orthologue gene tree (Fig. 3) at 100 loci. More than 200 loci per sample were recovered for the vast majority of samples, although there were some samples with a poor recovery rate.

**Figure 3:**
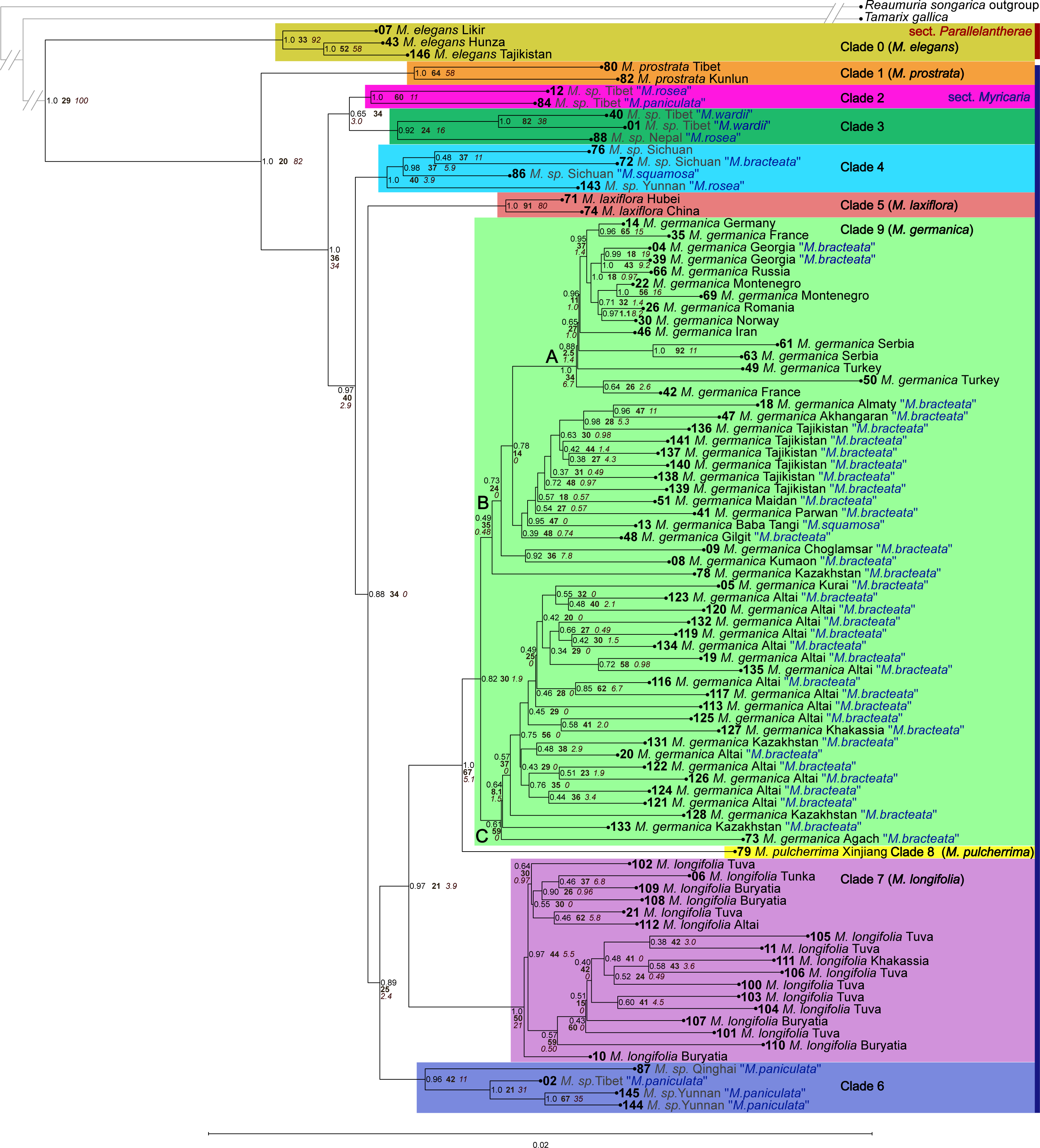
Orthologue-based coalescent species tree calculated with ASTRAL-IV. Tip labels show specimen number, new species identifications, and locality. Node labels are LPP, followed by sCF shown in bold, then gCF shown in italics. Specimen numbers correspond to maps (Fig. 1), SplitsTree analysis (Fig. 4; suppl. Fig. S4), other nuclear trees (Fig. S2; Fig. S3), plastid trees (Fig. 5; suppl. Fig. S5), photographs (Fig. 6-10), and lists of specimens sampled (suppl. Table S1; Appendix S4). Clade letter and colour codes correspond to groups shown in SplitsTree analysis (Fig. 4) and plastid species tree inferred with ASTRAL-IV (Fig. 5).

Plastid gene retrieval was more patchy; samples were either “good” (at least 61 loci recovered), or they were “bad” (33 or fewer loci recovered; see Appendix S4d). We removed samples with a recovery rate below 50 loci. This disparity was already noted during the DNA extraction and library preparation steps in line with the provenance of the specimens. Since the library preparation and target capture were optimised for the Angiosperms353 loci, this disparity is not reflected in the nuclear loci recovery rates. A summary of the numbers of loci assembled by CAPTUS for both the Angiosperms353 loci and the plastid loci can be seen in suppl. Fig. S1. Generally, the number of paralogs found was low. A total of 27,188 specimen-loci combinations (the vast majority) were single-copy. Meanwhile, 1,404 specimen-loci combinations were double-copy, four specimen-loci combinations were quadruple-copy, and four specimen-loci combinations were quintuple-copy. One locus in *Tamarix gallica* was 18-copy. Among *Myricaria* specimens for which at least 90% of the reference length was recovered, the number of paralogs found varied between 14 and 28; 59 paralogs were recovered for *Tamarix gallica*, and 83 paralogs were recovered for the *Reaumuria songarica* outgroup. Further details on the CAPTUS clean, extract, assemble and align processes can be seen in the CAPTUS reports, Appendices S4A-F.

### ANALYSES WITH THE ANGIOSPERMS353 DATASET PHYLOGENOMIC ANALYSES

The nuclear orthologue species tree (Fig. 3) was inferred using ASTRAL-IV with branch support given by local posterior probabilities (LPP). The site concordance factor (sCF) and gene concordance factor (gCF) values were also calculated, which give an idea of the proportion of gene trees and alignment sites supporting each bipartition, respectively. Generally, LPPs were higher (≥ 0.9) closer to the root of the tree, and lower towards the tips.

The Angiosperms353 nuclear dataset yielded a well-resolved species tree (Fig. 3) with consistently high support for major clades (clades labelled 0-9; LPP ≥ 0.82). Clade 0 (*M. elegans*) was strongly supported and recovered as sister to the rest of *Myricaria* (LPP = 1.0; sCF = 33; gCF = 92). Clade 5 (*M. laxiflora*) was also strongly supported (LPP = 1.0; sCF = 91; gCF = 80), as was clade 7 (*M. longifolia*) (LPP = 1.0; sCF = 50; gCF = 21). All specimens collected in or west of the Caucasus also formed a clade (clade 9, subgroup A: LPP = 1.0). In contrast, specimens from the Himalayan region (assigned to clades 1–4 and 6) did not form monophyletic groups and, taken together, formed grades near the root of the tree. These clades showed low gene concordance (gCF = 3.9 - 58) despite high LPP values (mostly ≥ 0.9), indicating limited agreement among gene trees.

In addition to this orthologue-based coalescent tree (Fig. 3), a species tree was inferred with ASTRAL-Pro (suppl. Fig. S2); this species tree was inferred after concatenating the homologue trees, which were generated after paralogues were removed (Fig. 2; inset F). A maximum likelihood tree from the supermatrix of the nuclear Angiosperms353 loci (suppl. Fig. S3) was also generated with IQ-TREE; this tree was generated from the concatenated orthologue alignments of the nuclear genes (Fig. 2; inset H). These trees were very similar to the orthologue-based coalescent tree seen in Fig. 3, concatenated from the orthologue trees inferred with IQ-TREE (Fig. 2; inset I).

### SplitsTree analysis

A SplitsTree analysis (Fig. 4) with the neighbour-net algorithm was run using the supermatrix alignment of the Angiosperms353 nuclear loci, without an outgroup and including only members of *Myricaria*, as the algorithm is prone to long-branch attraction. A second run including the outgroup can be found in suppl. Fig. S4, so that the position of *Reaumuria songarica* and *Tamarix gallica* can be seen. This analysis complements the sCF values in the nuclear orthologue species tree (Fig. 3) as it visualises site discordance in the supermatrix. To reduce crowding in the diagram, specimen numbers have only been shown on the tips, but letter and colour coding of groups correspond to that of the nuclear orthologue species tree (Fig. 3). The diagram was also split up into zones A to D for easier viewing. Clade 0 (gold), shown in zone A, represents the *M. elegans* clade, whose long branch showed that its sequences were more different than the rest of the specimens. The groups shown in zone B correspond to clades 1 to 6 in the nuclear orthologue species tree. This zone showed non-tree-like topology near its roots, reflecting the low gCF and sCF values for these groups as shown in Fig. 3. The group represented by zone C corresponds to clade 6 and the single *M. pulcherrima* specimen. There was also non-tree-like topology near its roots, albeit less so than in zone A. Zone D (clade 7, *M. longifolia*) showed the least non-tree-like topology, with branching showing a largely tree-like topology.

**Figure 4:**
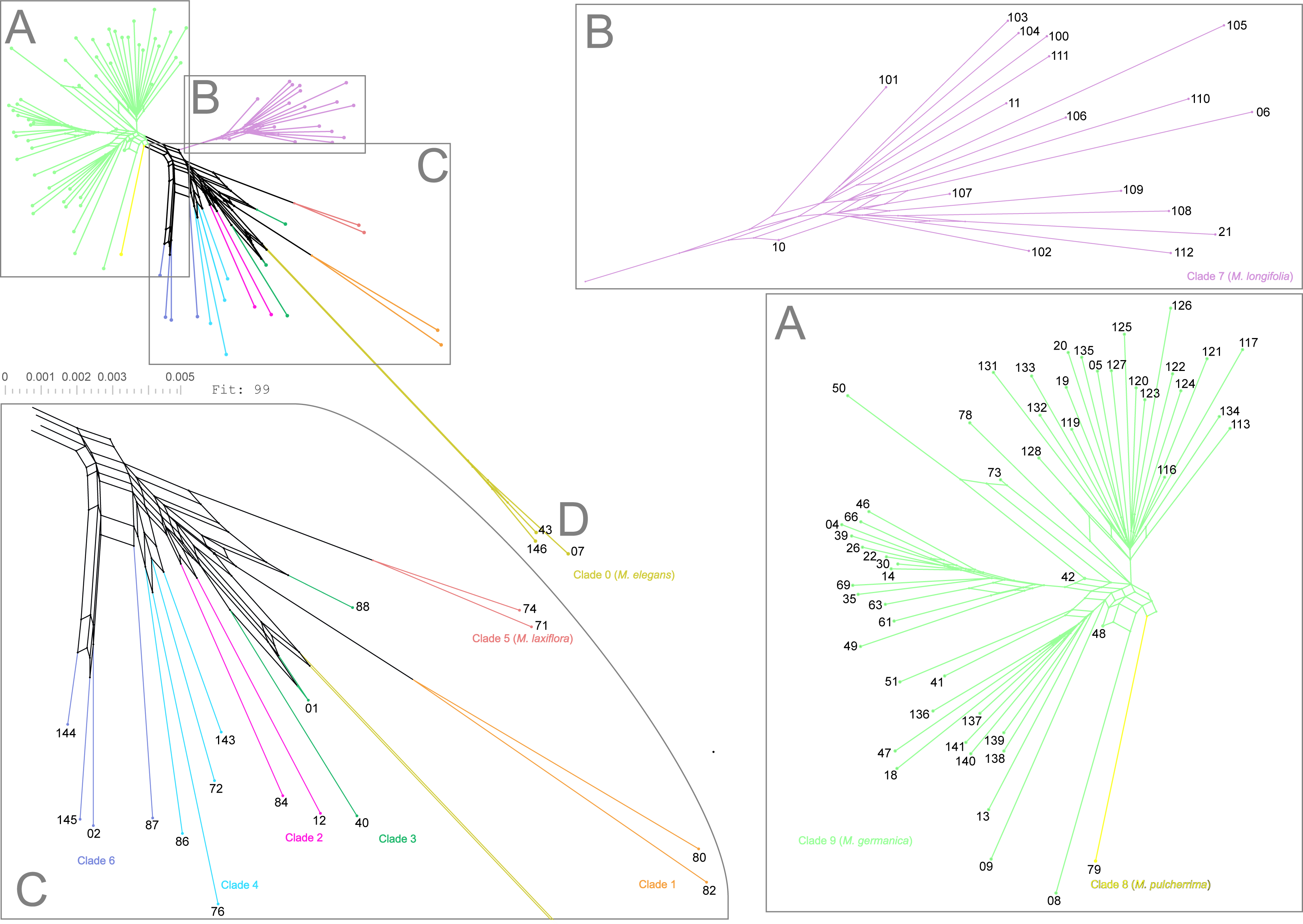
SplitsTree analysis using nuclear Angiosperms353 supermatrix alignment. Analysis run with neighbour-net algorithm including only *Myricaria* specimens and no outgroup. Specimen numbers correspond to maps (Fig. 1), nuclear orthologue species tree (Fig. 3), other nuclear trees (Fig. S2; Fig. S3), plastid trees (Fig. 5; suppl. Fig. S5), photographs (Fig. 6-10), and lists of specimens sampled (suppl. Table S1; Appendix S4). Clade letter and colour codes correspond to groups shown in orthologue-based species tree (Fig. 3) and plastid species tree inferred with ASTRAL-IV (Fig. 5).

### Comparisons with the plastid dataset

For completeness, and as a comparison to previous studies using chloroplast data (Hu & al., 2023; Sun & Werth, 2024), a species tree was inferred from plastid genes (introns and exons) using ASTRAL-IV, with LPP as support values (Fig. 5). Specimen numbers, clade numbers, and colouring correspond to the ASTRAL-IV nuclear species tree (Fig. 3) as well as the SplitsTree analysis (Fig. 4). The plastid orthologue species tree (Fig. 5) contained fewer samples than the nuclear Angiosperms353 trees due to the lower gene retrieval rates for plastid loci (suppl. Fig. S1B). Support values in this plastid tree (Fig. 5) were generally lower (LPP < 0.7 in most nodes) than in the nuclear orthologue species tree (Fig. 3), reflecting both the smaller number of plastid loci and the reduced number of base pairs. A concatenated supermatrix tree was also calculated (suppl. Fig. S4) for comparison of analytical approaches. The plastid results are therefore presented as complementary evidence to the nuclear phylogeny.

**Figure 5:**
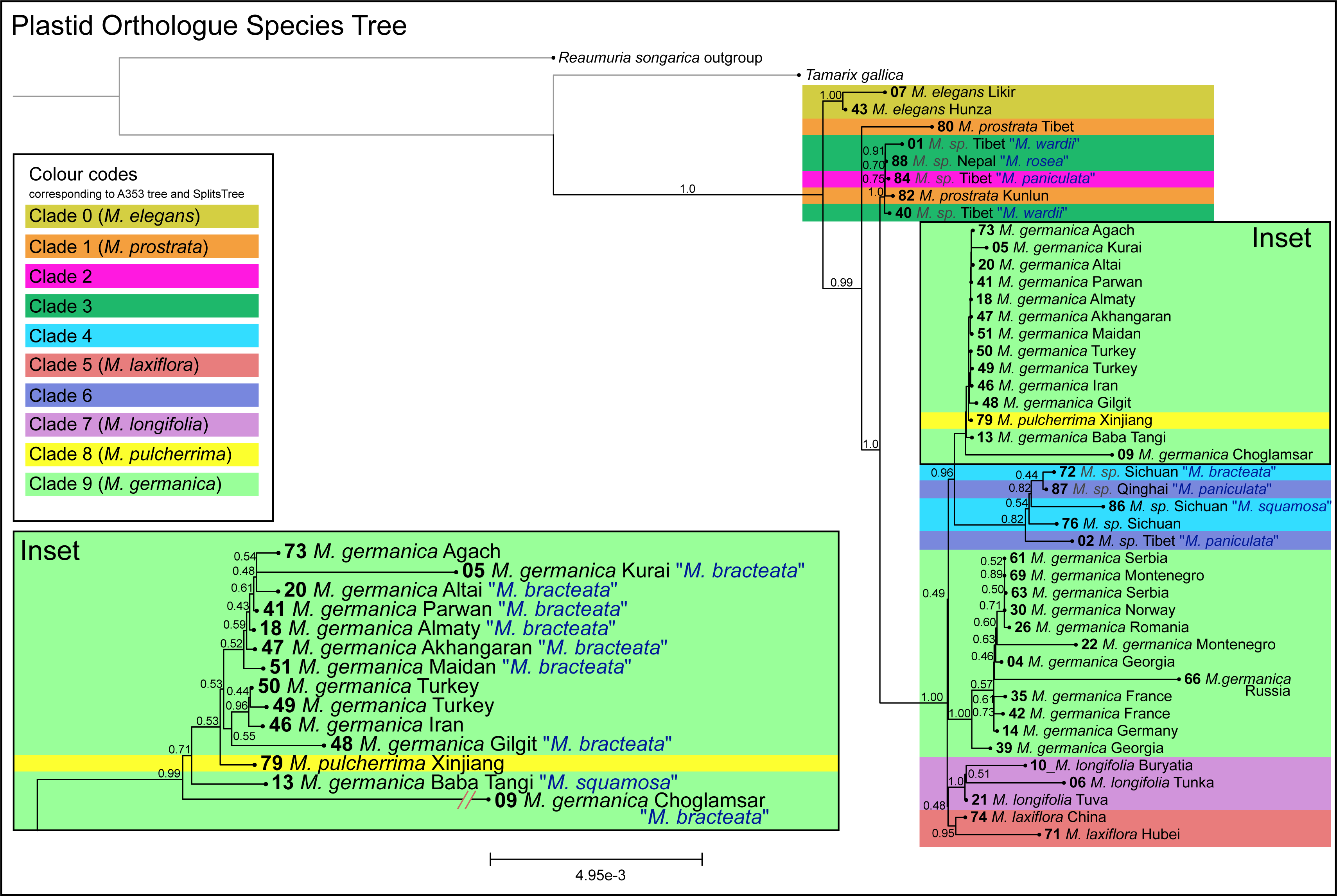
Plastid species tree calculated with ASTRAL-IV. Tip labels show specimen number, new species identifications, and locality. Node labels are LPP, followed by sCF shown in bold, then gCF shown in italics. Specimen numbers correspond to maps (Fig. 1), nuclear orthologue species tree (Fig. 3), other nuclear trees (Fig. S2; Fig. S3), SplitsTree analyses (Fig. 4; suppl. Fig. S4), supermatrix plastid tree (suppl. Fig. S5), photographs (Fig. 6-10), and lists of specimens sampled (suppl. Table S1; Appendix S4). Clade letter and colour codes correspond to groups shown in nuclear orthologue species tree (Fig. 3) and SplitsTree analysis (Fig. 4).

Similarities between the nuclear orthologue species tree (Fig. 3) and the plastid orthologue species tree (Fig. 5) include the strong support for individuals in clade 0 (*M. elegans*) as sister to the rest of *Myricaria*. The plastid species tree (Fig. 5) showed that specimens 07 and 43 (clade 0, *M. elegans*) were the sister to the rest of the genus (LPP = 1.0). The next lineage to branch in the plastid tree (Fig. 5) was one of the two *M. prostrata* specimens (specimen 80, clade 1) included in this analysis (LPP = 0.99). The rest of the genus formed a well-supported clade (LPP = 1.0). Another similarity between these two trees (Fig. 3; Fig. 5) was the strong support for clade 7 (*M. longifolia*), both with LPP = 1.0.

Beyond these points, the nuclear orthologue species tree (Fig. 3) and the plastid orthologue species tree (Fig. 5) showed substantial topological differences. The main difference is that specimens diagnosed as *M. germanica* in the plastid coalescent tree (Fig. 5) did not form a clade as they do in the nuclear orthologue species tree (Fig. 3; clade 9). In the nuclear orthologue species tree (Fig. 3), *M. pulcherrima* (specimen 79) was a sister to the *M. germanica* clade (clade 9), whereas in the plastid orthologue species tree (Fig. 5), this individual was nested among individuals from the *M. germanica* clade in Fig. 3. This may be since only one specimen of *M. pulcherrima* could be sourced for this study. Another discrepancy between the topologies of the nuclear (Fig. 3) and plastid (Fig. 5) orthologue species trees is the placement of *M. laxiflora* (clade 5, Fig. 3) and *M. longifolia* (clade 7, Fig. 3) individuals. The low support value (LPP = 0.48) of the clade containing the *M. longifolia* individuals 10, 06, and 21, and the *M. laxiflora* individuals 74 and 71 indicates uncertainty of placement in the plastid orthologue species tree (Fig. 5). In addition, individuals from clades 1, 2, 3, 4, and 6 in the nuclear orthologue species tree (Fig. 3) did not form the same clades in the plastid orthologue species tree (Fig. 5), as indicated by the colour codes.

### MORPHOLOGICAL OBSERVATIONS

Morphological observations were conducted on all specimens included in the molecular analyses to evaluate the congruence between phylogenetic structure and current species delimitations. Diagnostic characters previously used in the literature were tested across phylogenetic clades.

For clade 9, species delimitations proposed by Bobrov (1967) were treated as testable hypotheses. Bobrov distinguishes *M. germanica* and *M. bracteata* based on leaf size, raceme length, and bract morphology. These characters were assessed across all sampled individuals within this clade. Leaf length (typically 3–5 mm in *M. germanica*), raceme length (5–10 cm in *M. germanica* vs. 10–20 cm in *M. bracteata*), and bract shape (lanceolate vs. trapezoidal and membranous) were evaluated for consistency within and between phylogenetic groupings. Additional diagnostic characters described in other taxonomic treatments (e.g. Wu & al., 2007; Webb, 1968; Tsarenko & Peregrym, 2013), including the presence of imbricate scales, terminal flowers, and variation in inflorescence position, were also examined.

To summarise these observations, imbricate scales can be found on specimens from France, Afghanistan, and the Altai (Fig. 6). Raceme diversity also varied across clade 9, as individuals have both terminal (specimens 14, 09 and 117) and lateral racemes (specimens 35, 05), across the three subgroups of clade 9 (denoted by the letters A, B, and C in Fig. 3 and Fig. 7). Racemes also vary in length, with specimens 14, 51 and 116 showing long racemes 10 cm or greater, whereas specimens 35, 18, and 117 show raceme length shorter than 7 cm (Fig. 7). Terminal flowers can also be found on representatives from each clade 9 subgroup (Fig. 3: A, B and C), for example specimens 04, 140 and 121 (Fig. 8) respectively. Leaf length is also a variable trait – larger leaves of 3 – 5 mm can be found on specimens 30, 41, and 19, compared to smaller leaves found on specimens 14, 51, and 116 (Fig. 9). Finally, differing bract shapes can be seen across clade 9, with individuals such as 14, 123, and 05 featuring a more pointed, lanceolate morphology, while individuals 69, 09, and 19 feature bracts wider near the centre and with wider membranous margins (Fig. 10).

**Figure 6:**
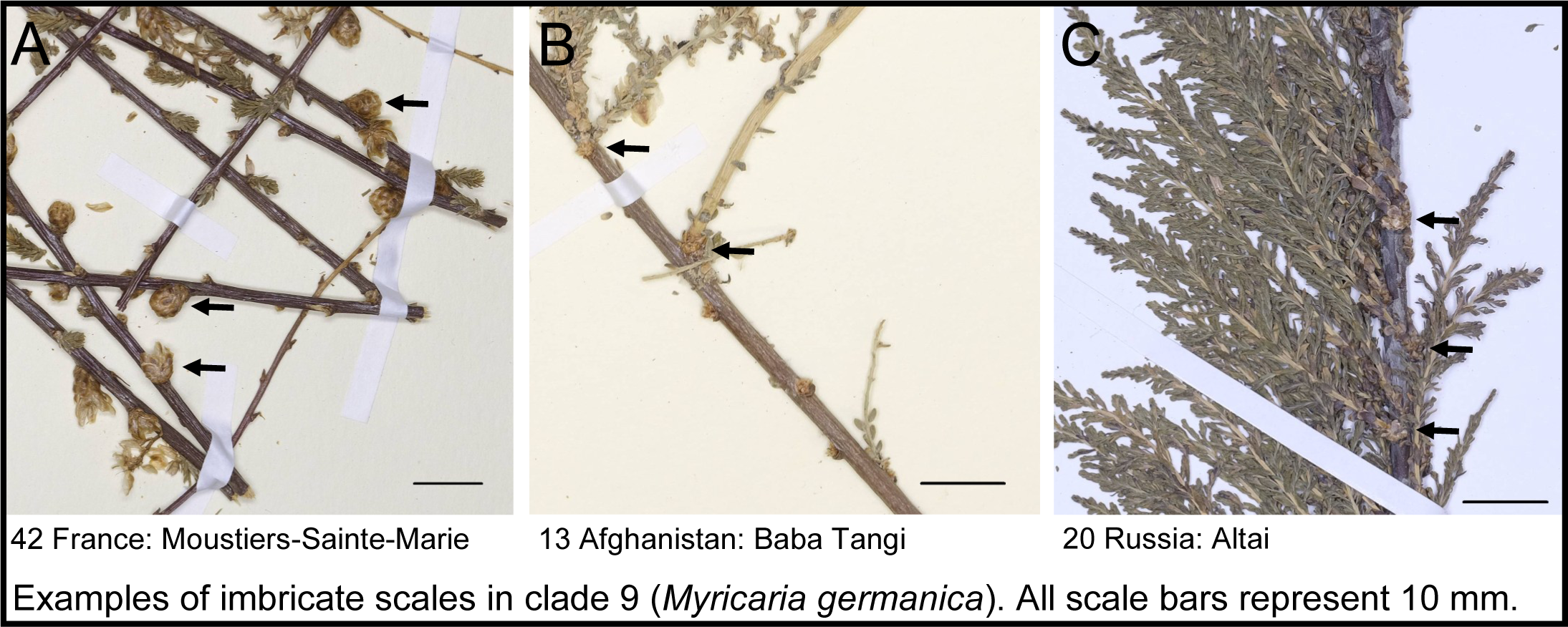
Photographs showing presence of imbricate scales on *M. germanica* specimens. Specimen numbers correspond to maps (Fig. 1), nuclear species trees (Fig. 3; Fig. S2; Fig. S3), SplitsTree analyses (Fig. 4; suppl. Fig. S4), plastid trees (Fig. 5; suppl. Fig. S5), and lists of specimens sampled (suppl. Table S1; Appendix S4). Herbarium specimens digitised by K. Weitkämper.

**Figure 7:**
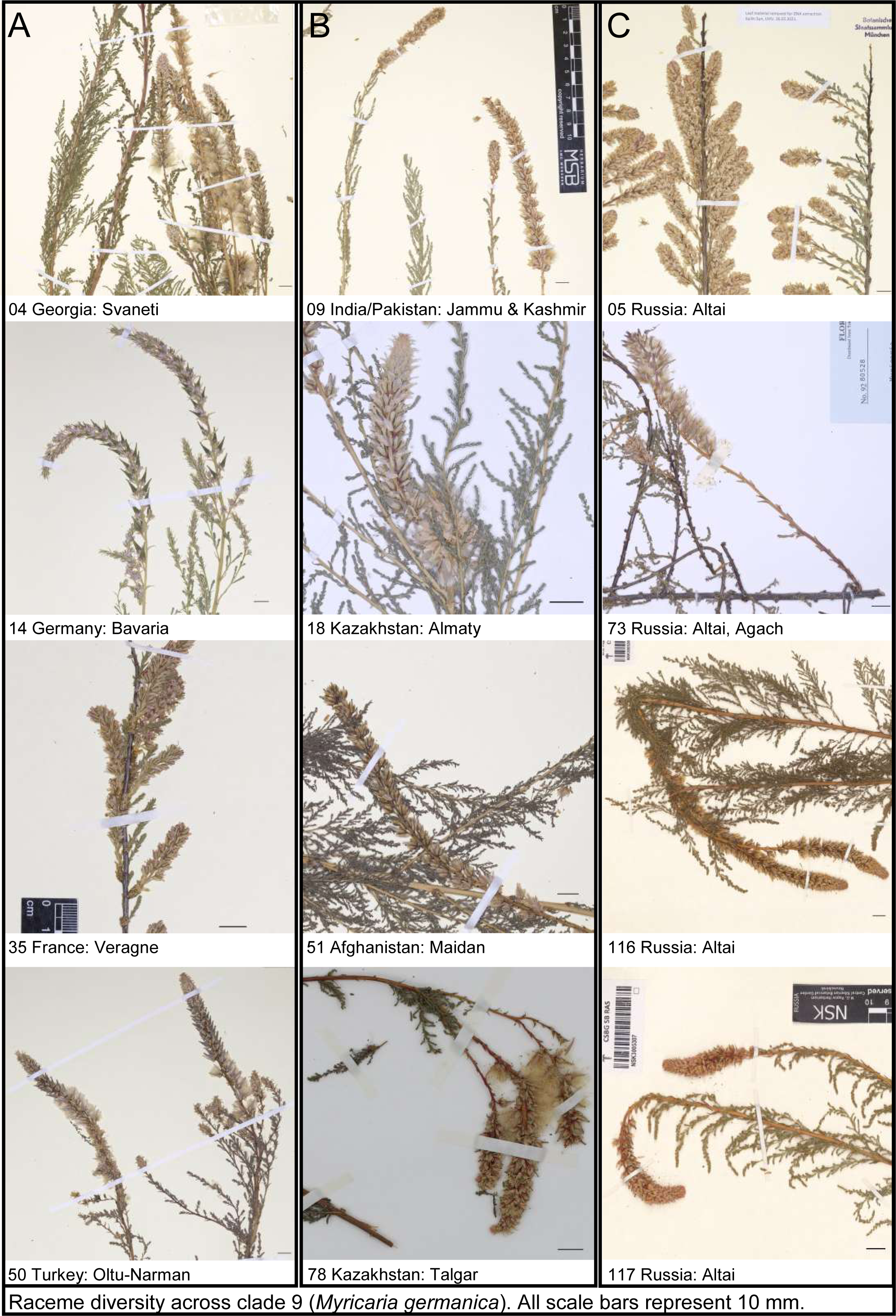
Photographs showing diversity of racemes on *M. germanica* specimens. Specimen numbers correspond to maps (Fig. 1), nuclear species trees (Fig. 3; Fig. S2; Fig. S3), SplitsTree analysis (Fig. 4), plastid trees (Fig. 5; suppl. Fig. S5), and lists of specimens sampled (suppl. Table S1; Appendix S4). Herbarium specimens 116 and 117 digitised by E. Lyakh; specimen 78 digitised by GOET; remaining specimens digitised by K. Weitkämper.

**Figure 8:**
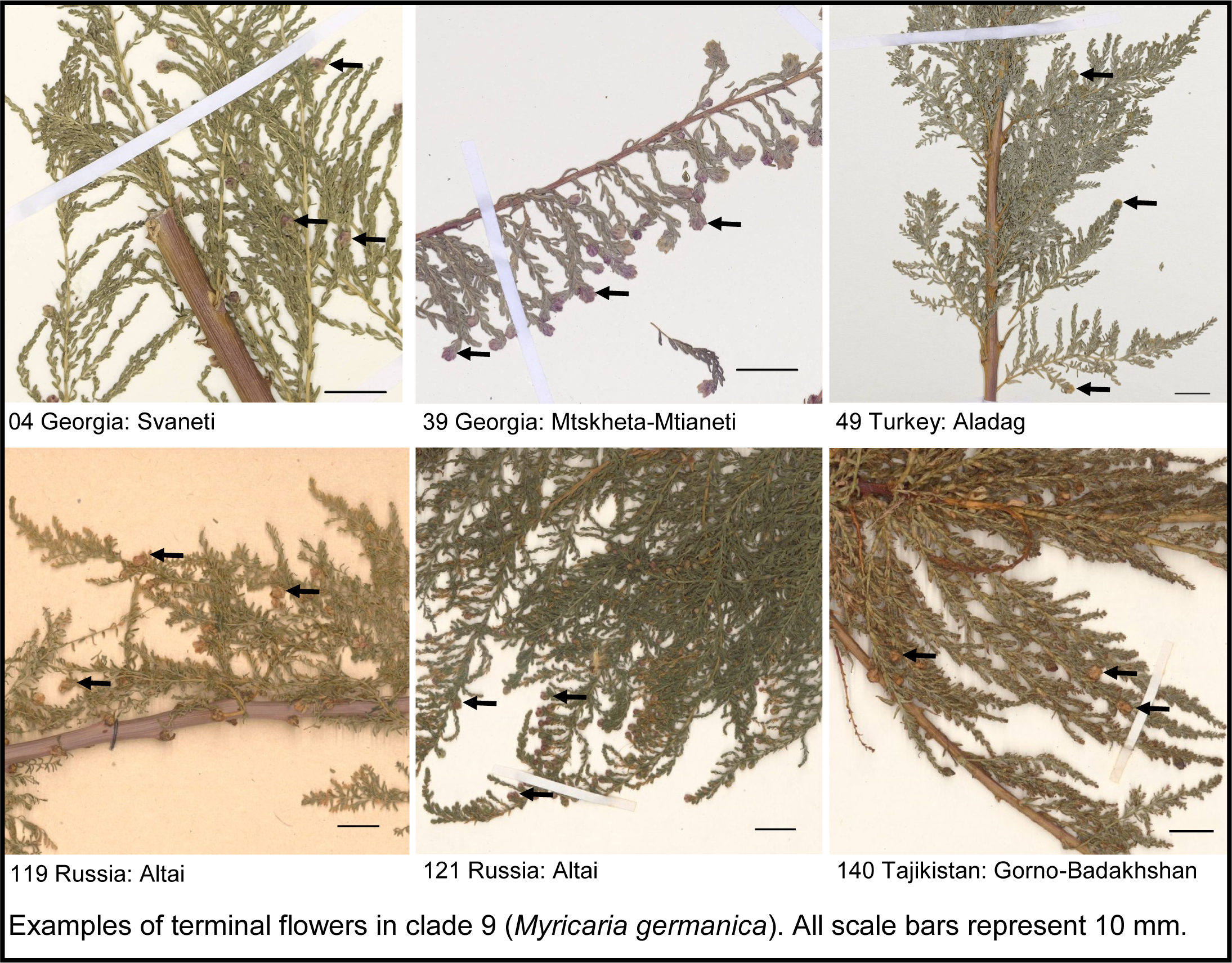
Photographs showing presence of terminal flowers on *M. germanica* specimens. Specimen numbers correspond to maps (Fig. 1), nuclear species trees (Fig. 3; Fig. S2; Fig. S3), SplitsTree analyses (Fig. 4; suppl. Fig. S4), plastid trees (Fig. 5; suppl. Fig. S5), and lists of specimens sampled (suppl. Table S1; Appendix S4). Herbarium specimens 04, 39, and 49 digitised by K. Weitkämper, specimens 119, 121, and 140 digitised by E. Lyakh.

**Figure 9:**
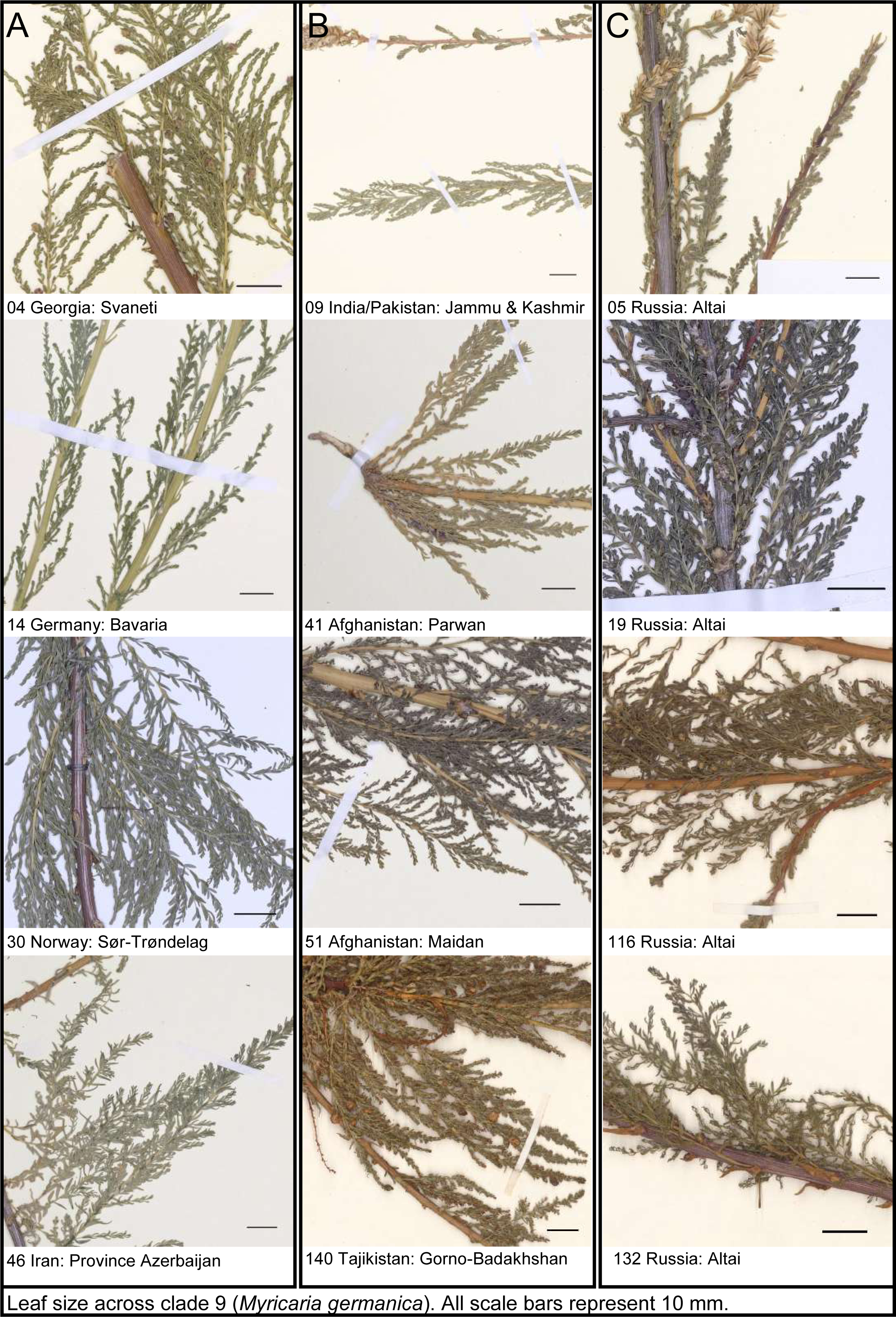
Photographs showing diversity of leaf size *M. germanica* specimens. Specimen numbers correspond to maps (Fig. 1), nuclear species trees (Fig. 3; Fig. S2; Fig. S3), SplitsTree analyses (Fig. 4; suppl. Fig. S4), plastid trees (Fig. 5; suppl. Fig. S5), and lists of specimens sampled (suppl. Table S1; Appendix S4). Herbarium specimens 116, 132, and 140 digitised by E. Lyakh, remaining specimens digitised by K. Weitkämper.

**Figure 10:**
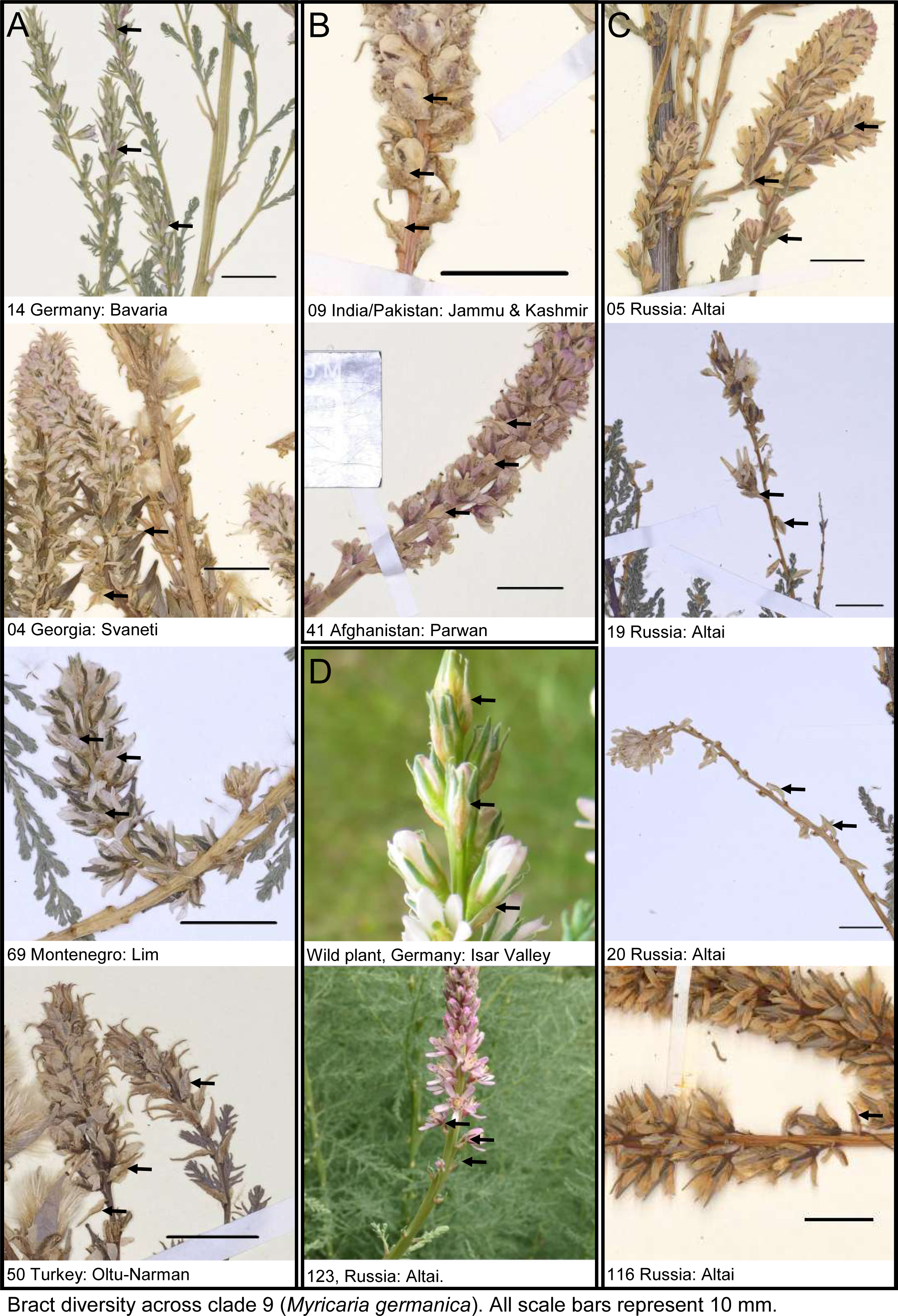
Photographs showing diversity of bract morphology on *M. germanica* specimens. Specimen numbers correspond to maps (Fig. 1), nuclear species trees (Fig. 3; Fig. S2; Fig. S3), SplitsTree analyses (Fig. 4; suppl. Fig. S4), plastid trees (Fig. 5; suppl. Fig. S5), and lists of specimens sampled (suppl. Table S1; Appendix S4). Herbarium specimen 116 digitised by E. Lyakh; remaining herbarium specimens digitised by K. Weitkämper; photograph of *M. germanica* in Isar valley by K. Weitkämper; photograph of specimen 123 by E. Lyakh. TABLES

For specimens belonging to the Himalayan and Central Asian clades (clades 1–4 and 6), morphological identification followed the keys provided in the Flora of China (Wu & al., 2007), and the resulting species assignments were compared to phylogenetic placement. A summary of these observations is provided in suppl. Table S2. Further, high-resolution images of type specimens of each putative species were located and were examined for key morphological features – details on the location of type specimens can be seen in the Taxonomic Treatment section.

## DISCUSSION

### PHYLOGENOMIC RESOLUTION AND CONCORDANCE

This study is the first to present a phylogeny of *Myricaria* based on nuclear genes, sequenced with NGS methods using the Angiosperms353 bait set via target enrichment. The nuclear orthologue species tree (Fig. 3), inferred from approximately 200 loci per individual using ASTRAL-IV, provides a well-resolved phylogeny, with consistently high support across major clades (clades 0–9; LPP ≥ 0.82). Using the multispecies coalescent approach, this analysis integrates multiple loci and accounts for incomplete lineage sorting.

While LPP reflects support for a given topology based on quartet frequencies, gCF and sCF values quantify the proportion of gene trees and alignment sites supporting each branch, respectively. High LPP values, especially nearer the base of the tree, indicate robustness of this species tree. Meanwhile, the gCF values across the tree can be interpreted as a proxy for the percentage of genes, from an individual specimen’s point of view, which are more similar within the clade than outside the clade. This suggests that at nodes with low gCF values, it would be hard to find morphological features that are characteristic of the following clade (for a more in-depth explanation of this, see the example on bird phylogenomics presented in Lanfear & Hahn, 2024, pp. 10-11). In the nuclear orthologue species tree (Fig. 3), this reflects the especially low gCF values (gCF < 1) along the terminal nodes within clade 9 (*M. germanica*), where individuals show far less morphological diversity.

The phylogenies inferred from plastid data (Fig. 5, suppl. Fig. S5) differ considerably from the nuclear orthologue species tree (Fig. 3) in topology. This discrepancy is not unexpected, because plastid genes are maternally inherited and are generally non-recombining. Our plastid phylogeny shows similar results to previous studies using plastid loci (Hu & al., 2023; Sun & Werth, 2024). For example, the plastid trees in all these studies indicate that *M. elegans* is a well-supported clade. The phylogenetic trees inferred from whole chloroplast genomes by Hu & al. show the so-called “*M. squamosa complex*”, a large clade with individuals diagnosed as *M. bracteata*, *M. paniculata*, and *M. squamosa*. This is similar to the clade with LPP = 0.49 in our plastid orthologue species tree (Fig. 5) containing individuals previously diagnosed as *M. bracteata*, *M. germanica*, *M. paniculata*, and *M. squamosa*. This similarity in results suggests that plastid data on its own (regardless of number of base pairs) provide limited resolution for species-level relationships in *Myricaria*.

The SplitsTree analyses (Fig. 4 and Fig S4.) with the neighbour-net algorithm visualise all supported splits in the data, and where splits contradict, the analysis produces boxes. The analysis produced non-tree-like topologies among most of the groups (with the exception of zone D in Fig. 4, corresponding to clade 7, *M. longifolia* in the nuclear orthologue species tree in Fig. 3). Hybridisation or introgression could be a factor contributing to the discordance between nuclear and chloroplast trees, as could be incomplete lineage sorting. There is evidence of a gene duplication event in the evolution of Tamaricaceae (Yang & al., 2018), as well as evidence of reticulate evolution in the sister genus *Tamarix* (Villar & al., 2019). However, we have found little evidence of polyploidy within *Myricaria*, since the vast majority of our 27,188 specimen-loci combinations were single copy (suppl. Fig. S1). The chromosome number of *M. germanica* and *M. laxiflora* is reported in previous studies to be 2n=24 (Löve, 1980; Xiang & al. 2025, respectively).

### BIOGEOGRAPHIC AND EVOLUTIONARY PATTERNS

The nuclear orthologue species tree (Fig. 3) recovers basal clades (clades 0–4) with high support (LPP ≥ 0.9), whereas support for nodes nearer the tips is lower. These basally branching clades with strong support contain species collected in the QTP and Himalayan regions, near to the centre of origin of the genus (Sun & Werth, 2024). The next clades to branch off are clades 5 and 6, which include specimens collected in Hubei, Tibet, Qinghai, and Yunnan. Clades 7 (*M. longifolia*), 8 (*M. pulcherrima*), and 9 (*M. germanica*) were the last to diverge. *Myricaria germanica* and *M. longifolia* are found in particular regions which are geographical cul-de-sacs of the distribution. For example, *M. germanica* occupies the westernmost areas of Eurasia, where it is not possible to spread west any further. *M. longifolia* occupies the northeasternmost parts of the distribution, geographically isolated from the rest of the genus by the Taklamakan and Gobi Deserts. This is reflected in the plastid orthologue species tree (Fig. 5), where European *M. germanica* specimens as well as *M. longifolia* both have strong statistical support (both LPP = 1.0), whereas support for other clades indicated in the nuclear orthologue species tree (Fig. 3) in the rest of the tree is generally weak.

In areas closer to the QTP centre of origin, species boundaries are fuzzier, with multiple samples assigned different names occurring in the same clades (such as *M. paniculata*, *M. rosea*, and *M. wardii*), as well as non-treelike topology in the SplitsTree (Fig. 4) analysis indicative of reticulate evolution. This could be due to the higher interconnectedness between habitats in the QTP alpine regions, leading to gene flow. Further, the unique habitat type of montane riparian ecosystems might create strong selection pressure against divergent morphology in *Myricaria*. The similar morphological characteristics between individuals in clades 2, 3, 4 and 6 could thus be explained by this selection pressure. These similarities also make it difficult to assign clear species boundaries and contribute to the confusion surrounding current species names.

There were some indications of a signal of ancestral gene flow between clades. Firstly, some strongly supported clades in the nuclear orthologue species tree (clades 0, 1, 2, 4, 5, and 7) all contain specimens that were collected in similar geographical areas. Second, the aforementioned clades had a non-tree-like appearance with major reticulations in the SplitsTree analysis (Fig. 4). This topology could indicate ancestral gene flow between the clades.

There is evidence from across the Angiosperm Tree of Life (Zuntini & al., 2024) that diversification is associated with gene conflict. The major discrepancies between the nuclear orthologue species tree (Fig. 3) and the plastid orthologue species tree (Fig. 5), as well as the low gCF and sCF values in areas of the nuclear orthologue species tree, could be linked to recent diversification within *Myricaria*. Recently published divergence time estimates place the origin of extant *Myricaria* lineages at around 22.8 Ma (Sun & Werth, 2024); in fact, *Myricaria* lineages currently found in Europe diverged from their Asian relatives only around 8.8 Ma (late Miocene). Therefore, gene flow between populations may have been possible until recently (or might even still be possible between certain geographical areas).

### TAXONOMIC IMPLICATIONS

An overview of our taxonomic revision can be seen in Table 2, where we provide a list of species that we accept, as well as the infrageneric classifications and synonyms for each accepted species. A further discussion of these names can be found in the Taxonomic Treatment section.

**Table 2:**
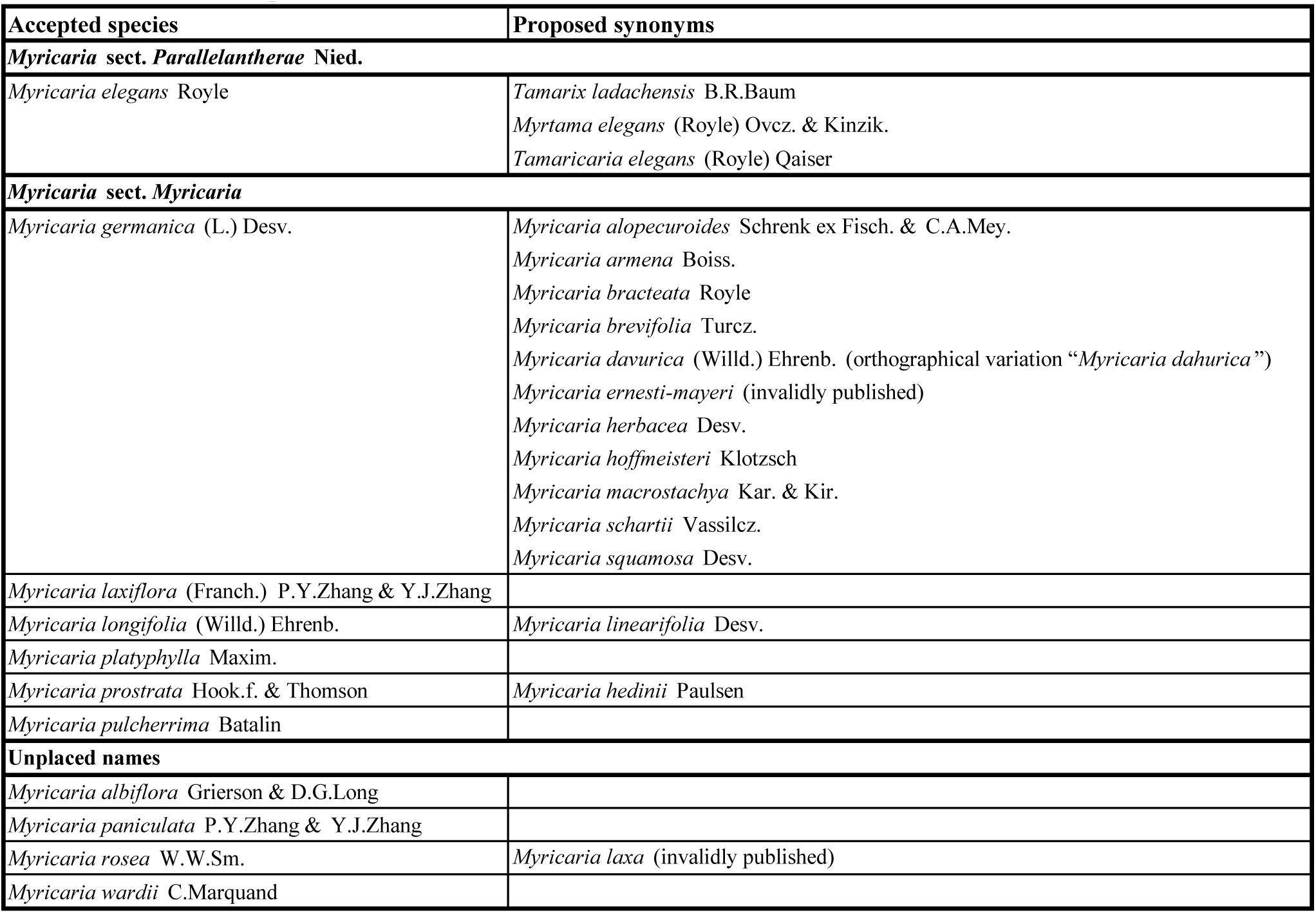
List of revised species names.

#### *Myricaria elegans* as sister to the remainder of the genus

Our analyses (Fig. 3; Fig. 5) consistently place *M. elegans* (clade 0) as sister to all other *Myricaria* species, in agreement with previous studies (Gaskin & al., 2004; Liu & al., 2006; Wang & al., 2009; Zhang & al., 2014; Hu & al., 2023; Sun & Werth, 2024). On the basis of its distinctive morphology, including free (unfused) stamens and differences in pollen structure, *M. elegans* has been segregated as a separate genus (*Myrtama*) by Ovchinnikov & Kinzikaeva (1977) and Qaiser & Ali (1978). Alternatively, infrageneric classifications have recognised its distinctiveness at sectional rank; for example, Niedenzu (1895) proposes the section *Parallelantherae* consisting of only *M. elegans*, with the remaining species placed in sect. *Renantherae*. In the revised infrageneric classification by Zhang & al. (2014), *M. elegans* continues to occupy an independent section.

*Myricaria elegans* has distinctive morphological features (unfused stamens, larger, lax leaves), which differ from those of other species within *Myricaria,* such as those found in the Asian highland clades (clades 2, 3, 4, and 6; Fig. 3), which have very similar morphologies. This species also consistently places as sister to the rest of *Myricaria* in molecular phylogenies. Phylogenetic evidence supports both taxonomic interpretations as monophyletic. However, we believe that the morphological and molecular differences are not sufficient to justify placement in its own genus. For example, other species which have distinctive features within *Myricaria* include *M. pulcherrima*, with large, cordate leaves, while *M. longifolia* and *M. platyphylla* are distinctive in their long, thin leaves and broad, lax leaves respectively.

Phylogenetically, the placement of *M. elegans* as sister to the remainder of the genus is stable across datasets, supporting its recognition as a distinct lineage. However, our SplitsTree analysis indicates some degree of gene tree conflict in this region of the phylogeny, which may reflect incomplete lineage sorting or historical gene flow, as expected in relatively recent radiations. The relatively young crown age of the genus (Sun & Werth, 2024), which includes *M. elegans*, is estimated at 22.8 Ma, another factor supporting the integration of *M. elegans* into the genus. We therefore retain *M. elegans* within *Myricaria*, but recognise its distinctiveness at the infrageneric level by reinstating *Myricaria* sect. *Parallelantherae* Nied.

Synonyms of *M. germanica*.

We observe that previously reported morphological differences in the species *M. germanica*, *M. bracteata*, and *M. squamosa* as described by Bobrov (1967) and other authorities (Gorshkova, 1974; Webb, 1968; Wu & al., 2007; Tsarenko & Peregrym, 2013) are neither supported by the molecular data nor by our observations from herbarium specimens. In our nuclear orthologue phylogeny, clade 9 shows 3 subgroups (Fig. 3, clade 9, A, B and C). These groupings broadly correspond to geographic regions, though LPP support is low for subgroups B and C, which consist of specimens collected east of the Caucasus (A with LPP = 1.0; B with LPP = 0.49; C with LPP = 0.61). None of the assessed morphological features is consistent between these three groupings. Imbricate scales, a defining feature of *M. squamosa* (Wu & al., 2007), occur throughout clade 9 and are not restricted to any subgroup (Fig. 3; Fig. 6). Raceme length is also not indicative, as specimens with varying raceme sizes can also be found across all three subgroups in the clade (Fig. 3; Fig. 7). Single terminal flowers were also observed on specimens across all subgroups of clade 9, ranging from Georgia to Siberia (Fig. 3; Fig. 8), suggesting that this trait may be environmentally influenced. In addition, variations in leaf size also do not sort by subgroups in the clade (Fig. 3; Fig. 9). Finally, bract shapes and sizes also vary across subgroups in clade 9 (Fig. 3), indicating that this feature is also variable (Fig. 10). The lectotype of *M. bracteata*, held at LIV (barcode LIV 1952.121.1759), was collected by J. F. Royle “on the road to Cashmere”; some specimens in clade 9 were also collected in nearby areas, such as Kumaon (specimen 08), Choglamsar (specimen 09), and Gilgit (specimen 48). This erect specimen shows small leaves, and a short raceme of around 3 cm, falling within the range of morphological variability within the sampled individuals of clade 9.

The protologue of *M. germanica* (basionym *Tamarix germanica*; Linne, 1753) does not provide much detail on the morphology of the species, only that its habitat is sometimes flooded, and that this species can be found in Germany. In his publication introducing the genus *Myricaria*, Desvaux (1825) describes *M. germanica* as having short, sharp, subimbricated leaves; flowers on peduncles; and ascending capsules. This basic description covers all the individuals in clade 9 and *M. germanica sensu lato*.

Plastid phylogenies have so far proved inconclusive for delimiting species boundaries. Hu & al. (2023) demonstrate that specimens diagnosed as *M. bracteata* and *M. squamosa* form unresolved polytomies on both their ITS and plastome trees, the so-called *M. squamosa* complex. However, Hu & al. (2023) did not include any European specimens, the type location of *M. germanica*, so their study could not test whether *M. germanica* would have been placed in the same polytomy. It could be argued that the differences in our plastid gene tree (Fig. 5) compared to the nuclear orthologue tree (Fig. 3) also support synonymising *M. bracteata* with *M. germanica*. In the nuclear orthologue tree (Fig. 3), specimens 49 and 50 from Turkey and specimen 46 from Iran form a clade with western European specimens and the Caucasian specimens (04 and 39) with strong support for specimens near the Caucasus. Meanwhile, on the plastid orthologue species tree (Fig. 5), specimens 46, 49, and 50 form a clade with the Asian specimens with strong support (LPP = 0.99); this indicates that there may well be intermixing among populations traditionally treated as *M. bracteata* and *M. germanica*.

In addition to the lack of phylogenetic and morphological differentiation between the putative species *M. germanica* and *M. bracteata*, there is also little ecological differentiation. Previous studies on *M. germanica* and *M. bracteata* populations (Kudrnovsky & Stöhr, 2013; Tsarenko & Peregrym, 2013; Werth & Scheidegger, 2014; Astashenkov & Lyakh, 2020; Sitzia & al., 2021) consistently report similar life histories of *M. germanica* and *M. bracteata*, suggesting that there are no major differences in habitat, ecology, or growth. Where there have been attempts to diagnose new species based on ecology and morphometrics (Lakušić & Pavlović, 1971; Lakušić & al., 1974), our phylogenomic analyses show no genetic indication that the putative species *M. ernesti-mayeri* should be seen as a species separate from *M. germanica*. Individuals from the Balkans (specimens 22 and 69, Montenegro, Lim River, near locality of specimens collected by Lakušić & Pavlović, 1971) are nested within the *M. germanica* clade among species from other areas, including Germany, France, and the Caucasus.

Taken together, the absence of consistent morphological, phylogenetic, and ecological differentiation provides no support for maintaining these taxa as separate species. We therefore propose synonymising *M. bracteata* and *M. squamosa* with *M. germanica*. As the earliest validly published name (Linnaeus (1753) published the basionym), *M. germanica* has nomenclatural priority, and is therefore supported as an accepted species.

#### Myricaria in Siberia and the Altai: Myricaria longifolia and Myricaria germanica

The morphological studies of Lyakh (2013) on Siberian *Myricaria* populations reveal that there are only two species in the region. In that publication, these two species are referred to as *M. bracteata* and *M. longifolia*. This finding is reflected in our nuclear orthologue species tree (Fig. 3), which shows that specimens collected from Siberia fall either into clade 7 (*M. longifolia*) or clade 9 (*M. germanica,* referred to by Lyakh (2013) as *M. bracteata*).

As discussed earlier, we propose synonymising *M. bracteata* with *M. germanica*; therefore the two species present in this region are *M. germanica* and *M. longifolia*. We thereby support retaining *M. longifolia* as an accepted species. Meanwhile, *M. davurica* was first described (as *Tamarix davurica*) by Willdenow (1816) after a sheet collected by Pallas. These specimens have very small, blunt leaves and morphologically clearly belong to *M. germanica* rather than *M. longifolia*. Therefore, we also propose synonymising *M. davurica* with *M. germanica*.

#### *Myricaria laxiflora* - the Three Gorges Dam region

*Myricaria laxiflora* (Fig. 3: clade 5) is consistently recovered as a well-supported and distinct lineage in phylogenetic studies (Liu & al., 2009; Wang & al., 2009; Zhang & al., 2014; Hu & al., 2023). Morphologically, however, it is only weakly differentiated from its closest relatives (Fig. 3: clades 6 and 9), with the primary diagnostic character being its lax inflorescences, as already noted in the protologue. The type specimen of this species was collected in “Chine orientalis, in faucibus Ou-Chan, vallis fluminis Yang-tze-Kiang” (near modern-day Wushan on the Yangtze Valley), and held at herbarium A (barcode 00063217), though this type consists only of a photograph and a small sample of plant material. We support retaining *M. laxiflora* as an accepted species because of its clear, well-supported phylogenetic placement and morphological differentiation via its lax inflorescences.

This is the only species of *Myricaria* found in central China at lower altitudes (70–160 m; Zhang & Zhang, 1984; Liu & al. 2009), in contrast to other *Myricaria* species, which are typically found at higher altitudes. The species is endemic to the Three Gorges Dam region of the Yangtze River. The controversial construction of the Three Gorges Dam in Hubei Province has therefore been a threat to *M. laxiflora* (Liu & al., 2009), as well as affecting the hydrochory of all plants in this section of the Yangtze River system (Liu & al., 2006).

#### *Myricaria pulcherrima* and *Myricaria platyphylla*: two large-leaved species found in Asia

In the nuclear orthologue species tree (Fig. 3), *M. pulcherrima* (clade 8) is sister to clade 9 (*M. germanica*); the combined clades of 8 and 9 received high support (LPP = 1.0). However, *Myricaria pulcherrima* is morphologically very distinct (larger, lax, and cordate leaves), so from that evidence we maintain the designation of *M. pulcherrima* as an accepted, separate species. Unfortunately, we were only able to source a single specimen of *M. pulcherrima*. The inclusion of this single individual in the nuclear orthologue tree would explain the relatively low LPP support (LPP = 0.82) of clade 9 (*M. germanica*).

Two specimens of *M. platyphylla* were sequenced, but unfortunately neither specimen had sufficient DNA quality to be included in the nuclear orthologue species tree. One specimen was included in the plastid species tree (Fig. 5), where it branches off after *M. laxiflora* (specimen 74). This is unsurprising, since this specimen (91) was collected in the Taklamakan Desert in modern-day Xinjiang (China), which is relatively close to the provenance location of the *M. laxiflora* specimens and *M. paniculata* (specimen 02, Tibet), the next specimen to branch off. *Myricaria platyphylla* specimens are also morphologically distinguished from other species by their large, non-cordate, lax leaves. Based on morphology, we support retaining *M. platyphylla* as an accepted species.

#### *Myricaria prostrata* - a highland species with prostrate habit

*Myricaria prostrata* (clade 1) is an alpine species defined by its prostrate habit, as well as stamens that are fused 2/3 from the base. It forms a well-supported clade and is sister to the rest of the species in the section *Myricaria*. Our results support *M. prostrata* as an accepted species.

#### The alpine grade

Despite strong support between the alpine clades (Fig 3: clades 2, 3, 4, and 6), we found no major consistent morphological differences from herbarium specimens. A table of our morphological findings for this clade can be found in suppl. Table S2. Potential names that can be assigned to these clades include *M. paniculata*, *M. rosea*, *M. wardii*, and possibly *M. albiflora*. We have not been able to obtain specimens diagnosed as *M. albiflora* for sequencing, since we have not found any herbaria outside of Edinburgh (E) using this name.

#### Infrageneric sections

Our results suggest that the section *Alpinae* proposed by Zhang & al. (2014) is not monophyletic. This is also true for the four series proposed earlier by Bobrov (1967), whose series *Elegantes*, for instance, contains *M. elegans* and *M. longifolia*. Based on our molecular evidence, we therefore reinstate the division into two sections introduced by Niedenzu (1895). However, we adopt the autonym *Myricaria* sect. *Myricaria* for the section containing the type species of the genus, *M. germanica*.

### FURTHER WORK

However, herbariomics alone is not able to resolve all species boundaries in the genus. Based on our results, we reiterate that the species boundaries of the Asian highland groups are still unclear. Possible names of these species include *M. albiflora*, *M. paniculata*, *M. rosea*, and *M. wardii*. We suggest that future taxonomic research on these groups should be carried out in the field or with live specimens, as morphological work with herbarium specimens and molecular studies seems to provide insufficient detail to fully resolve their species boundaries. Other possible future research could include greater sampling of different taxa represented in our study; this could include greater sampling of *Tamarix* to understand the relationships of genera in Tamaricaceae as a whole. A more detailed exploration of *M. germanica* subgroups could be done, and whether there are population-level differences can be observed with greater sampling depth. Population genetics methods could be useful for understanding patterns of gene flow across populations across Eurasia. Further morphological work could include detailed studies of *Myricaria* salt glands across different species, similar to Dörken & al. (2017), or microscopy of the pollen structures in *Myricaria*, like Qaiser & Ali (1978). These studies could also be helpful for understanding the evolution and loss of salt and drought tolerance in Tamaricaceae (Fradera-Soler & al., 2026).

### TAXONOMIC TREATMENT

The following taxonomic treatment is based on integrative evidence from phylogenomic data and morphological observations and conforms to current ICN standards (Madrid Code: Turland & al., 2025).

***Myricaria*** Desv. in Ann. Sci. Nat. 4: 345. 1825. *Type*: *M. germanica* (L.) Desv. Typified by Qaiser (1976).

*Notes*: Qaiser (1976) designates *M. germanica* as the lectotype.

***Myricaria*** sect. ***Parallelantherae*** Nied. in E. & P., Nat. Pfl. Fam. 3, 6 (1895) 296 – Type: *M. elegans* Royle.

*Note*: This is a monotypic section containing only *Myricaria elegans*.

1. *Myricaria elegans* Royle in Ill. Bot. Himal. 214. 1835 ≡ *Myrtama elegans* (Royle) Ovcz. & Kinzik. in Dokl. Akad. Nauk Tadzhiksk. S.S.R. 20(7): 56 (1977) ≡ *Tamaricaria elegans* (Royle) Qaiser & Ali Blumea 24: 153 (1978) ≡ *Tamarix ladachensis* B.R.Baum Monogr. Rev. Gen. Tamar.: 141 (1966). *Holotype:* INDIA: Numbool, Lippa Kunawar (Himalaya), Royle s.n. (LIV barcode 1952.121.1760 [photo!]). *Notes: Tamarix elegans* Spach. is not a synonym of *Myricaria elegans*; instead, this is a synonym of the currently accepted species *Tamarix chinensis* Lour. (POWO, 2026).

***Myricaria*** sect. ***Myricaria.*** Type: *M. germanica* (L.) Desv. = *Myricaria* sect. *Laxiflorae* M.L. Zhang. Type: *M. laxiflora* (Franch.) P.Y.Zhang & Y.J.Zhang – *Myricaria* sect. *Renantherae* Nied. in E. & P., Nat. Pfl. Fam. 3, 6 (1895) 296, nom. inval.

*Notes*: The section *Renantherae* is not validly published, because according to its protologue, it includes all species listed for the genus *Myricaria* mentioned by Desvaux (1825) (Art. 22.2, Turland & al., 2025). The section *Alpinae* suggested by Zhang & al. (2014) is also not validly published, because they do not designate a type. The species assigned to these sections are assigned to sect. *Myricaria*.

1. *Myricaria germanica* (L.) Desv. in Ann. Sci. Nat. (Paris) 4: 348 (1825) ≡ *Tamarix germanica* L. *Lectotype:* GERMANY: Ingolstadt. Burser XXV: 37 (UPS barcode [photo!]). Lectotypified by Jonsell and Karlsson (2010). = *Myricaria squamosa* Desv. in Ann. Sci. Nat. (Paris) 4: 350 (1825). *Holotype*: “in Oriente”, s.n. (P barcode P01902095 [photo!]). = *Myricaria herbacea* Desv. in Ann. Sci. Nat. (Paris) 4: 350 (1825). *Holotype*: Pall. in Fl. Ross. 2: 73 (1789), Plate LXXX B. = *Myricaria davurica* (Willd.) Ehrenb. Linnaea 2: 278 (1827) *Holotype*: RUSSIA, “Habitat in Dauuria”, Pallas s.n. (B barcode B-W 06072-01 0 [photo!]) = *Myricaria bracteata* Royle in Ill. Bot. Himal. Mts.: 214, t.44, f. 2 (1835). *Lectotype* (designated here): INDIA/PAKISTAN: “Road to Cashmere”, Royle s.n. (LIV barcode LIV 1952.121.1759 [photo!]). = *Myricaria brevifolia* Turcz. in Bull. Soc. Imp. Naturalistes Moscou 14: 70 (1840) *Holotype:* RUSSIA/MONGOLIA: “In humidis subarenosis Mongoliae chinensis”, 1831, I. Kusnetzov s.n., (LE barcode LE 01016269 [photo!]) = *Myricaria alopecuroides* Schrenk ex Fisch. & C.A.Mey. in Enum. Pl. Nov. 1: 65 (1841). *Lectotype*: RUSSIA: “Koksu Flußufer”, 14th June 1840, A. Schrenk s.n; specimen LE 01304808 [photo!] designated as lectotype by Gangurde & al. (2026). = *Myricaria macrostachya* Kar. & Kir in Bull. Soc. Nat. Mosc. 14(3): 423 (1841). Holotype: RUSSIA: “In Insula fluvii Naryu Prope Malonary in Skoi redut ad fines Chinensis sub flumen Julli omnino defloratam legimus” (LE). = *Myricaria hoffmeisteri* Klotzsch in *Botanische Ergebnisse der Reise seiner Königlichen Hoheit des Prinzen Waldemar von Preußen in den Jahren 1845 und 1846*: 120. 1862. *Neotype* (designated here): Klotzsch (1862), pl. XXV. = *Myricaria armena* Boiss. in Diagn. Pl. Orient., ser. 2, 2: 58 (1856). Lectotypified by Qaiser (1976). ARMENIA: “In valle Kassuklu, inter Baibout et Erzeroum”, Mai 1853, Huet du Pavillon s.n. (G barcode G00506651 [photo!]). = *Myricaria schartii* Vassilcz. in Bot. Mater. Gerb. Bot. Inst. Komarova Akad. Nauk S.S.S.R. 21: 6 (1961). *Holotype*: UZBEKISTAN: “Jugi Hissarici pars occidentalis, in syst. fl. Sangardak, in valle flum. Schart, prope vic. Bakhcza”, 27th August 1956, I. Cassilczenko et B. Sabiriv 39 (LE). – “*Myricaria ernesti-mayeri*” Lakušić in Zbor. Prvog Simp. Sist. Jugosl.: 75–83 (1971), nom. illeg. (*Myricaria dahurica* DC. in Prodr. 3: 98 (1828), orth. var.) *Notes:* The *M. squamosa* holotype collection locality is unclear. Herbarium sheet label reads: “Je crois que ce *Myricaria*en feuille est celui en fruit” - indicating that only the specimen on the left bearing fruits is *Myricaria*, the right-hand specimen is unknown. *Myricaria herbacea* is a synonym of *M. germanica*. A specimen from Bongard’s herbarium can be found in LECB (Byalt & Bubyreva, 2014), but this is unlikely to be a syntype because while Desvaux explicitly mentions having seen other specimens of *Myricaria* (denoted by “v. v.” and “v. s.”), he does not do so for *M. herbacea*. Ehrenberg notes that “Die Willdenowschen, von Pallas herstammenden Exemplare zeigen keinen merklichen Unterschied in der Größe der Blumen von *germanica*” (the specimens in the Willdenow collection, which originated from Pallas, show no obvious differences in flower size to that of *M. germanica*); Niedenzu (1895) synonymises *M. herbacea* with *M. alopecuroides*, which itself has been synonymised with *M. bracteata* by Bobrov (1967). The holotype of *M. davurica* (B barcode B -W 06072 -01 0) consists of two specimens, A and B, which appear to be from the same individual. The label, which reads “Habitat in Dauuria”, is on the outside of the folder and appears to have been written by Willdenow. The label “*Myricaria davurica* Ehbg.”, written by Ehrenberg, reflects his decision to transfer this species from *Tamarix* to *Myricaria*. It is unclear what Willdenow (1816) meant by “…in Sibirien dauurischen Alpen”. However, he distinguishes this habitat from the area surrounding Lake Baikal, since in his protologue for the then-named *Tamarix longifolia*, he cites the habitat of this species to be “in Sibirien jenseits des Baical-Sees”. An orthographic variant of the synonym *M. davurica* is “*M. dahurica*”; the first usage of this variant was by de Candolle (1828), which clearly states that it is the same name published by Willdenow (1816); he writes “*Tamarix dahurica* Willd. l c. n. 16.” (p. 98). The subsequent lectotypification of the *M. bracteata* illustration by Gangurde & al. (2026) is invalid, because both a specimen collected by Royle and an illustration were mentioned by Royle (1835) in the protologue. In accordance with Art. 9.12 of the ICN (Turland & al., 2025), the specimen should be chosen as lectotype over the illustration. This cited specimen was clearly collected before 1831, because Royle returned from his travels to England in 1831 and never left again (Harrison, 1978; Stafleu & Cowan, 1983, p. 961). It is unclear what is meant in the protologue (Turczaninow, 1840) of *M. brevifolia* by “Mongoliae chinensis”. In the 1830s, vast areas of modern-day Mongolia and modern-day Russia were considered vassals of the Qing Dynasty. It is likely that the collection locality of this specimen came from this area (Stieler & Reichard, 1833; area shown in green, labelled “Mongolei”). *Myricaria brevifolia* has been considered a synonym of *M. davurica* (Qaiser 1976), which is here synonymised with *M. germanica*. There are three sheets in the gathering of *M. alopecuroides* collected at “Koksu Flußufer”. Other specimens of the same gathering (LE barcode LE 01304808 [photo!] and LE 01304811 [photo!], LE 01304813 [photo!]) remain as residual syntypes. The original *M. hoffmeisteri* material collected by Hoffmeister was housed at B and was largely destroyed during the Second World War. The original type specimen of *M. hoffmeisteri* is presumed lost; the illustration is here designated as the neotype. The protologue of *M. armena* (Boissier, 1856) mentions two specimens: “Hab. in Armeniaev alle Kassuklu cl. Huet du Pavillon et circa Erzeroum cl. Calvert” (p. 58). Both specimens are held at G; the specimen collected by Huet du Pavillon “In valle Kaßuklu” has barcode number G00506651 [photo!], while the specimen collected by Calvert at “Erzeroum” has barcode number G00506652 [photo!]. Therefore, we correct the holotype cited by Qaiser (1976) as the lectotype. An isolectotype of the Huet du Pavillon specimen can be found at K (barcode K000641953 [photo!]). *Myricaria ernesti-mayeri*, first introduced by Lakušić & Pavlović (1971), is not a validly published name, as this publication does not provide sufficient details to designate a type specimen (no collector, date or specific location is cited).
2. *Myricaria laxiflora* (Franch.) P.Y.Zhang & Y.J.Zhang in Bull. Bot. Res., Harbin 4(2): 76 (1984) ≡ *Myricaria germanica* var. *laxiflora* Franch. in Nouv. Arch. Mus. Hist. Nat., sér. 2, 8: 205 (1885 publ. 1886). *Holotype*: CHINA, “China orientalis, in faucibus Ou-Chan, vallis fluminis Yang-tze-Kiang, Fl. Nov.” (near modern-day Wushan in the Three Gorges region), 1868, A. David s.n. (A barcode 00063217 [photo!]).
3. *Myricaria longifolia* (Willd.) Ehrenb. in Linnaea 2: 279 (1827) ≡ *Tamarix longifolia* Willd. in Abh. Königl. Akad. Wiss. Berlin 1812-1813: 85 (1816). *Lectotype* designated here: RUSSIA, “Habitat in Siberia”, Pallas s.n. (B barcode B -W 06071 -01 0 [photo!], specimen labelled A). = *Myricaria linearifolia* Desv. in Ann. Sci. Nat. (Paris) 4: 349 (1825). (*Tamarix decandra* Pall. in Fl. Ross. 2: 73 (1789), non auct. Salisb. nom. illeg.) *Notes:* Willdenow (1816) cites an illustration by Pallas in the protologue of *M. longifolia*, but no specimen was explicitly cited. The Willdenow collection (B) contains specimens collected by Pallas and labelled by Willdenow. This protologue was published posthumously in 1816, four years after his death in 1812, so the annotations in Willdenow’s handwriting “Habitat in Siberia” on the outside of the folder must have preceded the publication of the protologue. On the herbarium sheet (B barcode B -W 06071 -01 0), there are two gatherings: A and B. These specimens should be from the same collection, although if they were to be considered separate, specimen A accords most closely with the protologue and the illustration referenced therein (Ehrenberg (1827), pg. 279). We therefore designate specimen A as the lectotype. Ehrenberg (1827) also synonymises *M. linearifolia*, writing: “*M. linearifolia* Desv. ist gleich *T. longifolia* Willd., beide beziehen sich auf dieselbe Figur von Pallas” (p. 276).
4. *Myricaria platyphylla* Maxim. in Bull. Acad. Imp. Sci. Saint-Pétersbourg, sér. 3, 27: 425 (1882). *Holotype*: CHINA: “Mongolia occidentalis, mont. Alaschan australis”, June 1872, N. M. Przewalski, (LE barcode LE 01016270 [photo!]).
5. *Myricaria prostrata* Hook.f. & Thomson in Gen. Pl. 1: 161 (1862). *Lectotype*: CHINA/INDIA: “Tibet occ., descent from Lanak Pass, 13 - 15,000 ft.”, 13th September 1847, Thomson s.n. (K barcode K000641963 [photo!]). Lectotypified by Hara (1972). = *Myricaria hedinii* Paulsen in S.Hedin, S. Tibet 6(3): 54 (1922). Lectotypified by Qaiser (1976): CHINA: Tibet-Xinjiang border (36°01’00.0” N 87°46’00.0” E according to Hedin (1920), pg. 20), “Läger XXVI Norra Tibet”, 29th June 1901, Sven A. Hedin s.n. (S barcode S11-36916 [photo!]). *Notes:* In the protologue of *M. hedinii*, Paulsen (1922) mentions two specimens; the first is the specimen mentioned above (S barcode S11-36916), and the second is “a specimen of Hb. Hooker f. et Thomson, named *M. germanica* Desf. var. *prostrata* [sic], from W. Tibet, 13–15000 feet, agreed with it” (p. 55). Since two specimens are mentioned to be in agreement, and the author does not use the word “type” to designate a specimen, these two specimens should be seen as syntypes. Qaiser (1976) cites the first specimen as the holotype; here we correct this to lectotype.
6. *Myricaria pulcherrima* Batalin in Trudy Imp. S.-Peterburgsk. Bot. Sada 11: 483 (1891). Lectotype: CHINA: “H. Tarim: Jarkend-Darja”, 21st June 1889, W. J. Roborowski, (LE barcode LE 01016271 [photo!]). Lectotypified by Grubov (2000). UNPLACED NAMES *Myricaria albiflora* Grierson & D.G.Long in Notes Roy. Bot. Gard. Edinburgh 40: 116 (1982). *Holotype*: BHUTAN: West Bhutan, Stony river beds, Ha, 9100 ft, 13th June 1933, Ludlow & Sherriff 85 (E barcode E00318077 [photo!]). *Myricaria paniculata* P.Y.Zhang & Y.J.Zhang in Acta Phytotax. Sin. 22: 224 (1984). *Holotype*: CHINA: Yangsha, Lintan County, Gansu Province, alt. 2250 m, Zhang Peng-Yun 81053, (LZU barcode LZU 82011 [photo!]). *Note:* It is unclear what the collection date of this specimen is, as the date cited in the protologue (16^th^ July 1981) is not included on the specimen label (herbarium, location, collector, and collection numbers match). *Myricaria rosea* W.W.Sm. in Notes Roy. Bot. Gard. Edinburgh 10: 52 (1917). *Lectotype*: CHINA: “Mountains in the N.E. of the Yangtze bend, Yunnan. Lat. 27 degrees 45’ N. Alt. 10,000 ft.”, June 1913, G. Forrest 10609, (E barcode E00318099 [photo!]). Lectotypified by Hara (1972); subsequent lectotypification by Gangurde & al. (2026) is superfluous. – “*Myricaria laxa*” W.W.Sm in Notes Roy. Bot. Gard. Edinburgh 14: 123 (1923). nom illeg. *Note:* This name, *M. laxa,* is not validly published because there is insufficient description in the original publication (Articles 38.4 and 38.5, Turland & al., 2025; see examples). The single specimen mentioned in this publication, collected in July 1921 at “Tsarong, Eastern Tibet” and held at E (E barcode E00117579 [photo!]), was diagnosed as *M. rosea* by H. F. Comber (on herbarium label) and by Hara (1972). *Myricaria wardii* C.Marquand in J. Linn. Soc., Bot. 48: 166 (1929). *Holotype*: CHINA: “S.E. Tibet: Growing in the sand along the margins of pastures in the riverbed at Tsela Dzong, 9600 ft” (2900 m), 28th May 1924, F. Kingdon Ward 5708 (K barcode K000641961 [photo!]).

## CONCLUSIONS

Herbariomics provides an excellent opportunity to study plants with a wide distribution from habitats that are difficult to reach, as historical specimens can be sequenced without new fieldwork. There are a number of key findings that arise out of our molecular phylogenomic analyses, in conjunction with morphological observations. Firstly, we demonstrate that the series or sectional classifications proposed by Bobrov (1967) and Zhang & al. (2014) are not monophyletic, and we reinstate the sections introduced by Niedenzu (1895). We confirm that *M. elegans* (sect. *Parallelantherae*) is consistently sister to the rest of *Myricaria*, while the rest of the species (sect. *Myricaria*) include *M. germanica*, *M. laxiflora*, *M. longifolia*, *M. platyphylla*, *M. pulcherrima* and *M. prostrata*. Morphological observations indicate that individuals previously designated *M. bracteata* and *M. squamosa* are not distinguishable from individuals designated *M. germanica*, and that the names *M. bracteata* and *M. squamosa* should be seen as synonyms of *M. germanica*, which can be found in Europe through to the central Asian mountain ranges. We affirm the conclusion of Lyakh (2013) that there are only two species of *Myricaria* in Siberia - namely *M. germanica* and *M. longifolia*. *Myricaria laxiflora*, *M. platyphylla*, *M. pulcherrima* and *M. prostrata* should also be accepted as species, based on both morphological and molecular results. Unplaced names, including *M. albiflora*, *M. paniculata*, *M. rosea*, and *M. wardii*, are associated with the Asian highlands, and require further investigation. We also provide a thorough taxonomic overview of the published species names within *Myricaria*, and outline areas for potential further research.

## DATA AVAILABILITY

The data that support the findings of this study are openly available in the National Center for Biotechnology Information at https://www.ncbi.nlm.nih.gov/bioproject/1224993, reference number PRJNA1224993.

## AUTHOR CONTRIBUTIONS

**K.W.** and **S.W.** conceptualised this study; **K.W.** sourced herbarium material; **E.L.** collected and curated specimens; **K.W.** extracted DNA and sequenced the plant material; **D.M.-B.** and **F.W.** developed data analysis methodology; **K.W.**, **D.M.-B.**, and **F.W.** analysed the data; **K.W.** wrote the original draft; all authors reviewed and edited the manuscript; **K.W.** and **S.W.** acquired the funding.

## CURATORS AND COLLECTIONS

We thank the following curators and herbaria for their expertise and assistance: Hans-Joachim Esser, Andreas Fleischmann, Carla Novoa Sepúlveda, and Camila Uribe-Holguin (M and MSB); Anthony Brach (A and GH); Dmitar Lakušić (BEOU); Jiří Danihelka (BRNU); Marc Appelhans (GOET); Guangwan Hu (HIB); Rachel McCarthy (K); Irina Illarionova and Mariya Sheludyakova (LE); Donna Young (LIV); Jianbin Pan (LZU); George Miehe (MB); Vanessa Invernon (P); Chien-Wen Chen (TAIF); and Mats Hjertson (UPS).

## TECHNICAL AND LANGUAGE SUPPORT

We thank the Helmholtz Zentrum Munich for their high-quality Illumina sequencing service. In addition, we are grateful for the support of Andrea Brandl, Tanja Ernst, Lilia Galvez, Alina Höwener, Elizabeth Joyce, Agnes Scheunert, and Silvia Wienken in the lab, and for the provision of computing resources by Johannes Kinder, Martin Josko, and Sven Strickroth. We also thank Marc Gottschling for his academic support. We thank Luo Chen for his assistance with Chinese texts, and Marina Lyakh for her assistance in interpreting between Russian and English. Finally, we thank the anonymous reviewers for their constructive comments and feedback.

---

**Appendix S1: List of vouchers, new diagnosed species name with authority, location description, collection date, collector and collection number, herbarium code, NCBI BioSample Accession number. All sequences were newly generated for this study.**

***M. elegans*** Royle, India: Himachal Pradesh, 20. Jul. 1987, *H. Hartmann 3127*, (M), SAMN46873452; ***M. elegans*** Royle, Pakistan: Karakorum, 28. Jul. 1974, *F. Gartner s.n.*, (M), SAMN46873472; ***M. elegans*** Royle, Tajikistan: Gorno-badakhshan, 21. Aug. 2013, *E. M. Lyakh 52*, (NSK), SAMN46873539; ***M. prostrata*** Hook.f. & Thomson, China: Tibet, 13. Aug. 2005, *G. Miehe, S. Miehe NX05-094-01*, (MB), SAMN46873490; ***M. prostrata*** Hook.f. & Thomson, China: Xinjiang, 06. Jul. 1992, *G. Miehe, S. Miehe 8250*, (MB), SAMN46873491; ***Myricaria* sp.** China: Tibet, 14. Sep. 1989, *B. Dickoré 5261*, (M), SAMN46873457; ***Myricaria* sp.** China: Tibet, 13. Aug. 2004, *D. E. Bou ord, J. H. Chen, S. L. Kelley, J. Li, R. H. Ree, H. Sun, J. P. Yue, Y. H. Zhang s.n.*, (A), SAMN46873492; ***Myricaria* sp.** China: Tibet, 20. Aug. 1994, *B. Dickoré 11449*, (M), SAMN46873447; ***Myricaria* sp.** China: Tibet, 28. Sep. 1989, *B. Dickoré 5810*, (M), SAMN46873469; ***Myricaria* sp.** Nepal: Gandaki Zone, 26. Jun. 2003, *Y. Iokawa, T. Miyazaki, K. Yonekura, Y. Ibaragi, R. K. Uprety s.n.*, (A), SAMN46873495; ***Myricaria* sp.** China: Sichuan, 26. May. 2012, *X. Li 1271*, (TAIF), SAMN46873484; ***Myricaria* sp.** China: Sichuan, 16. Aug. 1934, *H. Smith 11085*, (UPS), SAMN46873487; ***Myricaria* sp.** China: Sichuan, 25. Jul. 2007, *D. E. Bou ord, K. Fujikawa, S. L. Kelley, R. H. Ree, Bo Xu, J. W. Zhang, T. C. Zhang, W. D. Zhu s.n.*, (A), SAMN46873493; ***Myricaria*** sp., China: Yunnan, 15. Jul. 2000, *E. M. Lyakh 58*, (NSK), SAMN46873536; ***M. laxiflora*** (Franch.) P.Y.Zhang & Y.J.Zhang, China: Hubei, 04. Dec. 2011, *D. Wang 952*, (TAIF), SAMN46873483; ***M. laxiflora*** (Franch.) P.Y.Zhang & Y.J.Zhang, China: no specific locality, no date recorded, *no author*, *s.n*., (HIB), SAMN46873486; ***Myricaria* sp.** China: Tibet, 02. Aug. 2009, *D.E. Bou ord, B. Bartholomew, D.A. Eaton, X.H. Li, R.H. Ree, B. Xu, J.P. Yue, J.W. Zhang, X.X. Zhu 41694*, (M), SAMN46873448; ***Myricaria* sp.** China: Qinghai, 01. Sep. 1996, *T. N. Ho, B. Bartholomew, M. F. Watson, M. G. Gilbert s.n.*, (GH), SAMN46873494; ***Myricaria* sp.** China: Yunnan, 13. Jul. 2000, *E. M. Lyakh 59*, (NSK), SAMN46873537; ***Myricaria* sp.** China: Yunnan, 13. Jul. 2000, *E. M. Lyakh 60*, (NSK), SAMN46873538; ***M. longifolia*** (Willd.) Ehrenb., Russia: Sayan Mountains, 15. Jun. 1968, *L. Malyschev, Y Petroczenko, S. Andrulaytis 5482*, (M), SAMN46873451; ***M. longifolia*** (Willd.) Ehrenb., Russia: Transbaikal, 28. Aug. 1968, *L. Nikolaeva, A. Tkaczenko 5481*, (M), SAMN46873455; ***M. longifolia*** (Willd.) Ehrenb., Russia: Tuva, 05. Jun. 1972, *V. Khanminczun 5690*, (M), SAMN46873456; ***M. longifolia*** (Willd.) Ehrenb., Russia: Tuva, 06. Aug. 2004, *J. Danihelka 2004/0256*, (BRNU), SAMN46873463; ***M. longifolia*** (Willd.) Ehrenb., Russia: Tuva, 29. Jul. 1987, *E. M. Lyakh 1*, (NSK), SAMN46873498; ***M. longifolia*** (Willd.) Ehrenb., Russia: Tuva, 27. Jul. 1987, *D. N. Shaulo 2*, (NSK), SAMN46873499; ***M. longifolia*** (Willd.) Ehrenb., Russia: Tuva, 29. Jul. 1987, *E. M. Lyakh 3*, (NSK), SAMN46873500; ***M. longifolia*** (Willd.) Ehrenb., Russia: Tuva, 05. Aug. 1995, *E. M. Lyakh 5*, (NSK), SAMN46873501; ***M. longifolia*** (Willd.) Ehrenb., Russia: Tuva, 05. Aug. 1994, *E. M. Lyakh 6*, (NSK), SAMN46873502; ***M. longifolia*** (Willd.) Ehrenb., Russia: Tuva, 16. Jul. 1994, *E. M. Lyakh 7*, (NSK), SAMN46873503; ***M. longifolia*** (Willd.) Ehrenb., Russia: Tuva, 18. Aug. 1995, *E. M. Lyakh 9*, (NSK), SAMN46873504; ***M. longifolia*** (Willd.) Ehrenb., Russia: Buryatia, 28. Jul. 2003, *E. M. Lyakh 10*, (NSK), SAMN46873505; ***M. longifolia*** (Willd.) Ehrenb., Russia: Buryatia, 22. Aug. 2004, *E. M. Lyakh 11’*, (NSK), SAMN46873506; ***M. longifolia*** (Willd.) Ehrenb., Russia: Buryatia, 18. Jul. 1995, *E. M. Lyakh 13*, (NSK), SAMN46873507; ***M. longifolia*** (Willd.) Ehrenb., Russia: Buryatia, 23. Aug. 2004, *E. M. Lyakh 15’*, (NSK), SAMN46873508; ***M. longifolia*** (Willd.) Ehrenb., Russia: Khakassia, 26. Jul. 1987, *E. M. Lyakh 18*, (NSK), SAMN46873509; ***M. longifolia*** (Willd.) Ehrenb., Russia: Altai, 10. Aug. 2016, *E. M. Lyakh 19’*, (NSK), SAMN46873510; *M. pulcherrima* Batalin, China: Xinjiang, 17. Aug. 1986, *B. Dickoré 0126*, (GOET), SAMN46873489; ***M. germanica*** Desv., Georgia: Svaneti, 07. Aug. 2009, *A. Gröger, W. Lobin, C. Lobin, Z. Schewardnadse 1716*, (M), SAMN46873449; ***M. germanica*** Desv., Russia: Kurai, 12. Jul. 1971, *E. Bobrov 5480*, (M), SAMN46873450; ***M. germanica*** Desv., India: Uttaranchal, 04. Oct. 2004, *B. Dickoré 19452*, (M), SAMN46873453; ***M. germanica*** Desv., India: Jammu and Kashmir, 26. Jul. 1992, *H. Hartmann 4191*, (M), SAMN46873454; ***M. germanica*** Desv., Afghanistan: Wakhan, 26. Jul. 1975, *H. Huss 23 C*, (M), SAMN46873458; ***M. germanica*** Desv., Germany: Bavaria, 12. Jul. 2020, *H. Krause 7*, (M), SAMN46873459; ***M. germanica*** Desv., Kazakhstan: Almaty, 07. Jul. 2014, *M. Chytry, M. Koci 2014 no. 525 duplum*, (BRNU), SAMN46873460; ***M. germanica*** Desv., Russia: Altai, 05. Aug. 2005, *M. Hajek, P. Hajkova 2005/81*, (BRNU), SAMN46873461; ***M. germanica*** Desv., Russia: Altai, 05. May. 2005, *P. Lustyk 2005/1283*, (BRNU), SAMN46873462; ***M. germanica*** Desv., Montenegro: Andrijevica, 29. Jul. 2014, *V. Kalnikova, T. Peterka, S. Palpurina, Z. Pleskova VK14/098*, (BRNU), SAMN46873464; ***M. germanica*** Desv., Romania: Alba, 16. Jun. 2010, *V. Grulich et al. VG10/157*, (BRNU), SAMN46873465; ***M. germanica*** Desv., Norway: Sor-Trondelag, 31. Aug. 2014, *V. Kalnikova, T. Peterka, T. Ruzickova, O. Knapek VK14/007*, (BRNU), SAMN46873466; ***M. germanica*** Desv., France: Laragne, 11. May. 1970, *A. Charpin, W. Greuter, O. Monthoux s.n.*, (M), SAMN46873467; ***M. germanica*** Desv., Georgia: Mtskheta-Mtianeti, 14. Aug. 2007, *L. Meierott Georg07/65*, (M), SAMN46873468; ***M. germanica*** Desv., Afghanistan: Parwan, 26. May. 1973, *O. Anders 9937*, (M), SAMN46873470; ***M. germanica*** Desv., France: Moustiers Sainte-Marie, 13. Apr. 1973, *E. Berger 4575*, (M), SAMN46873471; ***M. germanica*** Desv., Iran: Azerbaijan Province, 06. Jun. 1971, *K.H. Rechinger 4/270*, (M), SAMN46873473; ***M. germanica*** Desv., Uzbekistan: Tian Shan, 06. Jul. 1973, *V. Vašák s.n.*, (M), SAMN46873474; ***M. germanica*** Desv., Pakistan: Gilgit, 02. Jul. 1974, *S.M.A. Kazmi 5684*, (M), SAMN46873475; ***M. germanica*** Desv., Turkey: Ağrı, 24. Aug. 1976, *K.P. Buttler, W. Schmidt, I. Turgut, S. Yüce 21760*, (M), SAMN46873476; ***M. germanica*** Desv., Turkey: Oltu, 09. Jul. 1992, *M. Nydegger 46664*, (M), SAMN46873477; ***M. germanica*** Desv., Afghanistan: Maidan, 24. Aug. 1970, *D. Podlech 19489*, (M), SAMN46873478; ***M. germanica*** Desv., Serbia: Knjazevac-Kalna, 08. Jun. 1997, *“Vuk.S., Zlat.B, Rand.V.” 5694*, (BEOU), SAMN46873479; ***M. germanica*** Desv., Serbia: Knjazevac, 17. Jul. 1998, *V. Stevanovic, M. Niketic, G. Tomovic 12781*, (BEOU), SAMN46873480; ***M. germanica*** Desv., Russia: Ullu Tau, 17. Jul. 2007, *D. Milanovic 39334*, (BEOU), SAMN46873481; ***M. germanica*** Desv., Montenegro: Berane, 30. Jul. 2016, *D. Lakusic, N. Kuzmanovic, I. Jankovic 46516*, (BEOU), SAMN46873482; ***M. germanica*** Desv., Russia: Altai, 27. Jul. 1992, *M. Wakabayashi, M. Amano, N. V. Friesen, Y. Kadota, A. A. Krasnikov, I. M. Krasnoborov, M. N. Lomonosova, M. Minaki, M. M. Tamura 9280528*, (TAIF), SAMN46873485; ***M. germanica*** Desv., Kazakhstan: Almaty, 12. Aug. 2001, *“J. Homeier & Kazakhstan Exkursion 2001” 824*, (GOET), SAMN46873488; ***M. germanica*** Desv., Russia: Altai, 15. Aug. 1987, *E. M. Lyakh 20*, (NS), SAMN46873511; ***M. germanica*** Desv., Russia: Altai, 30. Jun. 1990, *E. M. Lyakh 24*, (NSK), SAMN46873512; ***M. germanica*** Desv., Russia: Altai, 30. Jun. 1990, *E. M. Lyakh 25*, (NSK), SAMN46873513; ***M. germanica*** Desv., Russia: Altai, 03. Sep. 1993, *E. M. Lyakh 28*, (NSK), SAMN46873514; ***M. germanica*** Desv., Russia: Altai, 04. Sep. 1993, *E. M. Lyakh 29*, (NSK), SAMN46873515; ***M. germanica*** Desv., Russia: Altai, 25. Jul. 2022, *E. M. Lyakh 31*, (NSK), SAMN46873516; ***M. germanica*** Desv., Russia: Altai, 11. Aug. 2016, *E. M. Lyakh 33*, (NSK), SAMN46873517; ***M. germanica*** Desv., Russia: Altai, 17. Aug. 1987, *E. M. Lyakh 44*, (NS), SAMN46873518; ***M. germanica*** Desv., Russia: Altai, 28. Jul. 2022, *E. M. Lyakh 34*, (NSK), SAMN46873519; ***M. germanica*** Desv., Russia: Altai, 22. Jul. 2022, *E. M. Lyakh 36*, (NSK), SAMN46873520; ***M. germanica*** Desv., Russia: Altai, 24. Jul. 2022, *E. M. Lyakh 37*, (NSK), SAMN46873521; ***M. germanica*** Desv., Russia: Khakassia, 27. Jul. 1987, *E. M. Lyakh 39*, (NSK), SAMN46873522; ***M. germanica*** Desv., Kazakhstan: Abai, 31. May. 2002, *E. M. Lyakh 46*, (NSK), SAMN46873523; ***M. germanica*** Desv., Kazakhstan: Jetisu, 24. May. 2016, *E. M. Lyakh 41*, (NSK), SAMN46873525; ***M. germanica*** Desv., Russia: Altai, 25. Aug. 2010, *E. M. Lyakh 38*, (NSK), SAMN46873526; ***M. germanica*** Desv., Kazakhstan: Almaty, 25. May. 2016, *E. M. Lyakh 42*, (NSK), SAMN46873527; ***M. germanica*** Desv., Russia: Altai, 12. Aug. 2016, *E. M. Lyakh 32*, (NSK), SAMN46873528; ***M. germanica*** Desv., Russia: Altai, 18. Aug. 2005, *E. M. Lyakh 30’*, (NSK), SAMN46873529; ***M. germanica*** Desv., Tajikistan: Hissar Valley, 17. Aug. 2013, *E. M. Lyakh 49*, (NSK), SAMN46873530; ***M. germanica*** Desv., Tajikistan: Gorno-Badakhshan, 21. Aug. 2013, *E. M. Lyakh 50*, (NSK), SAMN46873531; ***M. germanica*** Desv., Tajikistan: Gorno-Badakhshan, 21. Aug. 2013, *E. M. Lyakh 51*, (NSK), SAMN46873532; ***M. germanica*** Desv., Tajikistan: Gorno-Badakhshan, 22. Aug. 2013, *E. M. Lyakh 53*, (NSK), SAMN46873533; ***M. germanica*** Desv., Tajikistan: Gorno-Badakhshan, 23. Aug. 2013, *E. M. Lyakh 54*, (NSK), SAMN46873534; ***M. germanica*** Desv., Tajikistan: Gorno-Badakhshan, 23. Aug. 2013, *E. M. Lyakh 55*, (NSK), SAMN46873535; ***M. platyphylla*** Maxim., China: Inner Mongolia, 1872, *N.M. Przewalski 39*, (P), SAMN46873496; ***M. platyphylla*** Maxim., China: Taklamakan Desert, 1885, *N.M. Przewalski s.n.*, (P), SAMN46873497; ***M. germanica*** Desv., Kazakhstan: Abai, 01. Jun. 2002, *E. M. Lyakh 43*, (NSK), SAMN46873524.

## LIST OF SUPPLEMENTARY MATERIALS

### FIGURES

**Figure S1: Gene recovery statistics**

**Figure S2: Homologue nuclear species tree inferred using ASTRAL-PRO.** Tip labels show specimen number, library preparation number, new species identifications, and locality. Node labels are LPP. Specimen numbers correspond to maps (Fig. 1), nuclear orthologue species tree (Fig. 3), SplitsTree analysis (Fig. 4), plastid trees (Fig. 5; suppl. Fig. S4), photographs (Fig. 6-10), and lists of specimens sampled (suppl. Table S1; Appendix S4).

**Figure S3:** Supermatrix tree of Angiosperms353 loci inferred using IQ-TREE. Tip labels show specimen number, new species identifications, and locality. Node labels are bootstraps. Specimen numbers correspond to maps (Fig. 1), nuclear orthologue species tree (Fig. 3), SplitsTree analysis (Fig. 4), plastid trees (Fig. 5; suppl. Fig. S4), photographs (Fig. 6-10), and lists of specimens sampled (suppl. Table S1; Appendix S4).

**Figure S4:** SplitsTree analysis with neighbour-net algorithm using nuclear Angiosperms353 supermatrix alignment, with outgroup *Reaumuria songarica* and *Tamarix gallica*. Specimen numbers correspond to maps (Fig. 1), nuclear orthologue species tree (Fig. 3), other nuclear trees (Fig. S2; Fig. S3), plastid trees (Fig. 5; suppl. Fig. S5), photographs (Fig. 6-10), and lists of specimens sampled (suppl. Table S1; Appendix S4).

**Figure S5:** Supermatrix tree of plastid loci inferred using IQ-TREE. Tip labels show specimen number, new species identifications, and locality. Node labels are bootstraps. Specimen numbers correspond to maps (Fig. 1), nuclear orthologue species tree (Fig. 3), SplitsTree analysis (Fig. 4), plastid species tree calculated with ASTRAL-IV (Fig. 5), photographs (Fig. 6-10), and lists of specimens sampled (suppl. Table S1; Appendix S4).

### TABLES

**Table S1: Overview of sampled *Myricaria* specimens**

**Table S2: Morphological overview of specimens from clades 2, 3, 4, and 6.**

## APPENDICES

**Appendix S1: Herbarium voucher specimens**

**Appendix S2: Detailed description of bioinformatics and phylogenetic analyses**

**Appendix S3: Bioinformatics scripts Appendix S4(A-F): CAPTUS reports**

