## Appendix S2 for "Towards an understanding of the taxonomy of the Eurasian montane riparian genus *Myricaria* (Tamaricaceae) through Angiosperms353 herbariomics"

### Supplementary materials S2: Detailed bioinformatics pipeline

This workflow was done with Ubuntu 20.04 and Ubuntu 22.04. For ease of understanding, colour-coding is used in this document:

Scripts are denoted in blue: `pxclsq_wrapper.py`

Folders are denoted in red: `~/flowchart_1124/MAFFT031124`

Path to installation (location of program in computer) is denoted in grey: `$PATH_to_ASTRAL_Installation$`

Programs used in the pipeline are in bold: **ASTRAL-IV**

This document illustrates the workflow for the Angiosperms353 nuclear genes with exons. The workflow for the plastid genes with exons is the same, but instead using the PTD output from **Captus**.

First, install all programs needed for this pipeline

- **Captus** v.1.0.1 (Ortiz et al., 2023)
- **MAFFT** v.7.526 (Katoh and Standley, 2013)
- **Phyx** v.1.3.1 (Brown et al., 2017)
- **IQ-TREE** v.2.3.6 (Chernomor et al., 2016; Kalyaanamoorthy et al., 2017; Minh et al., 2020b; Mo et al., 2022)
- **TreeShrink** v.1.3.9 (Mai and Mirarab, 2018)
- **ASTRAL-Pro3** v.1.19.3.5 (Zhang et al., 2020; Zhang and 231 Mirarab, 2022a; Tabatabaee et al., 2023)
- **ASTRAL-IV** v.1.19.4.5 (Zhang et al., 2018; Zhang and Mirarab, 2022b; Tabatabaee et al., 2023)
- **Seaview** v.5.0.5 (Gouy et al., 2010)

Scripts needed to execute workflow (see Supplementary Materials 2)

Move these scripts to a separate scripts folder.

- `reformatCaptus.sh`
- `pxclsq_wrapper.py`
- `IQtreeatperl.sh`
- `mask_tips_by_taxonID_transcripts.py`
- `outgroup.txt`
- `tree_shrink_1.3.9_wrapper.py`
- `write_homolog_fasta_from_multiple_aln.py`
- `prune_paralogs_MO_1to1_MO.py`
- `write_ortholog_fasta_from_multiple_aln.py`
- `concatenate_matrices_phyx.py`
- `slice_supermatrix.sh`
- `outgroup.txt`

Documentation webpages:

- **Captus:** https://edgardomortiz.github.io/captus.docs/assembly/clean/preparation/index.html
- Dr. Diego F. Morales-Briones' BitBucket: https://bitbucket.org/dfmoralesb/target_enrichment_orthology/src/master/

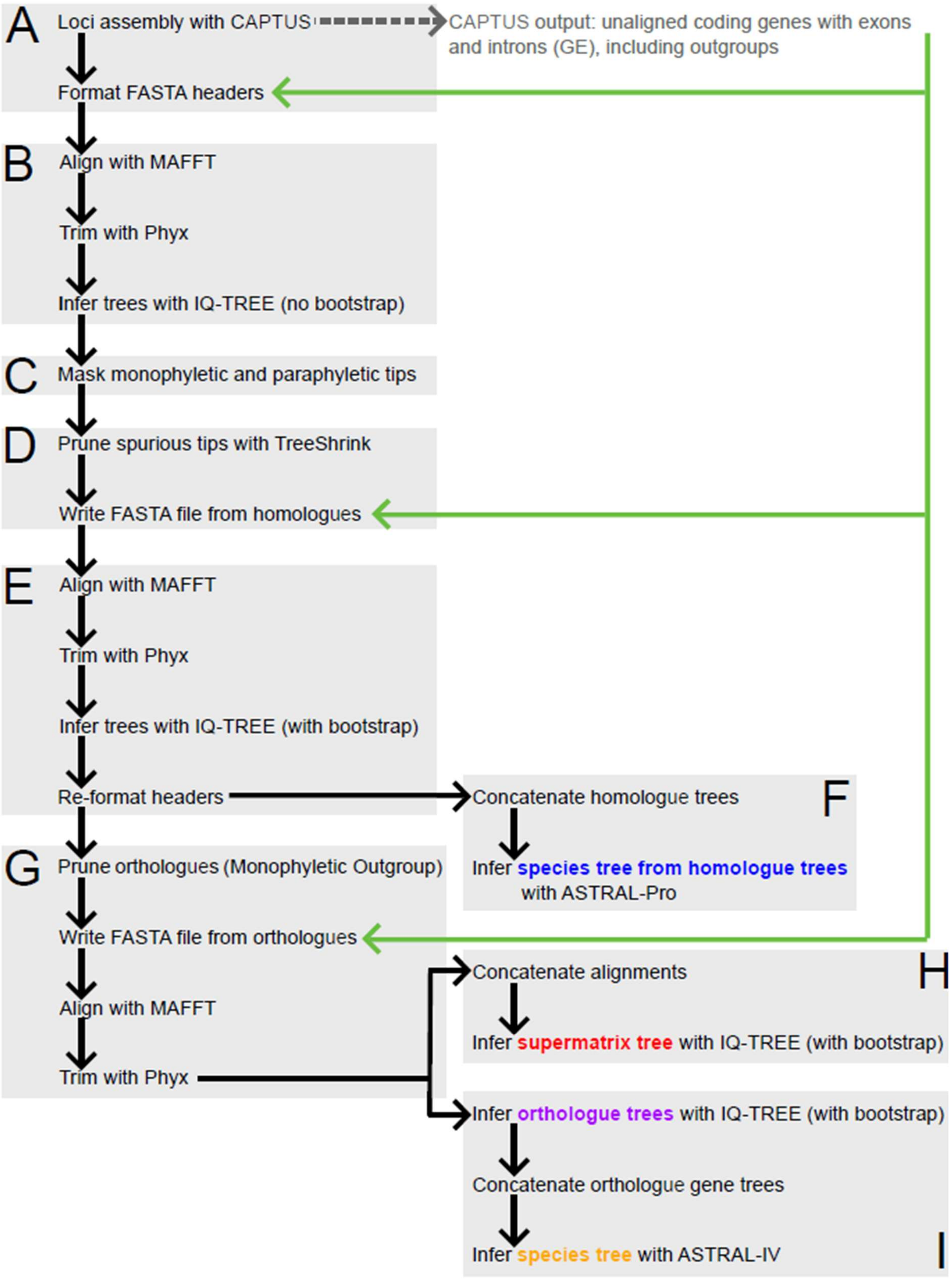


Reprint of **Figure 1** for ease of reference**:** Summary of bioinformatics pipeline. The black text names each step in the process, while the black arrows represents output of the previous step being used in the next step. The grey arrows show that the new FASTA files being written has been inputted directly from the original **Captus** output. The coloured text show the outputs of this pipeline.

**Steps A to I are described below.**

---

# A

<u>Loci assembly with **Captus**</u>

Input is raw sequence data from Helmholtz Institute. Outgroups (*Reaumuria songarica* and *Tamarix gallica*) was downloaded from GenBank (Sayers et al., 2021) and the relevant genes were also retrieved using the **Captus** pipeline.

Step 1: Clean using **Captus** default settings:

```
captus_assembly clean -r
```

Step 2: Assembly using **Captus** default settings:

```
captus_assembly assemble -r 01_clean_reads
```

Step 3: Extract:

```
captus_assembly extract -a 02_assemblies/ -n target.fasta --nuc_min_score 0.15 --nuc_min_identity 75 --nuc_min_coverage 50 -o 03
```

Step 4: "Align" (collect genes):

```
captus_assembly align -e 03_Scipio_Tamaricaceae_outgroup_extractions/ -m NUC -f NT,GE --filter_method none -o 04_Scipio_Tamaricaceae_outgroups_alignments
```

This step is necessary to collect the genes in the correct format for downstream analysis with **MAFFT**.

---

# B

<u>Align with **MAFFT**</u>

Input is non-aligned output of **Captus** with flanking exons:

`04_Scipio_Tamaricaceae_outgroups_alignments/01_unalign/01_coding_NT/03_genes`

Copy `reformatCaptus.sh` into that folder and run the following command there:

```
bash reformatCaptus.sh
```

Run the following command in the folder:

`04_Scipio_Tamaricaceae_outgroups_alignments/01_unalign/01_coding_NT/03_genes`

To align using **MAFFT** (with GNU parallel to optimise for multi-threading):

```
parallel mafft --nuc --auto --maxiterate 1000 --reorder {} '>' {}.mafft.aln ::: *.fna
```

Move the output files to separate folder (e.g. `~/flowchart_1124/MAFFT031124`).

Trim alignments using **Phyx**

Trim alignments using **Phyx** with Python 2 wrapper:

```
python2 pxclsq_wrapper.py ~/flowchart_1124/MAFFT031124 0.1 dna
```

Infer Trees using **IQ-TREE**

Induce phylogenetic trees using **IQ-TREE** with GNU parallel (without bootstrap) by running the following command in the folder `~/flowchart_1124/MAFFT031124`:

```
parallel iqtree2 -m MFP --seqtype DNA -s ::: *.aln-cln
```

Reformat the FASTA files with bash script to replace @ removed by **IQ-TREE**:

```
bash IQtreeatperl.sh
```

Move the output to a separate folder, e.g.

`~/flowchart_1124/IQTrees1_reformat_041124.`

---

# C

Mask monophyletic and paraphyletic tips

Mask monophyletic and paraphyletic tips using Python 2 script:

```
python2 mask_tips_by_taxonID_transcripts.py
~/flowchart_1124/IQTrees1_reformat_041124 .treefile
~/flowchart_1124/MAFFT031124 y ~/flowchart_1124/IQTrees1_mask_041124
```

---

# D

Prune spurious tips with **TreeShrink**

Create exceptions file in scripts folder "`outgroup.txt`" to avoid pruning outgroups.

Test **TreeShrink** with quantile 0.05, 0.10 and 0.50:

```
python tree_shrink_1.3.9_wrapper.py
~/flowchart_1124/IQTrees1_mask_041124 .treefile.mm 0.05
~/flowchart_1124/IQTrees1_shrink005_041124 outgroup.txt

python tree_shrink_1.3.9_wrapper.py
~/flowchart_1124/IQTrees1_mask_041124 .treefile.mm 0.10
~/flowchart_1124/IQTrees1_shrink010_041124 outgroup.txt

python tree_shrink_1.3.9_wrapper.py
~/flowchart_1124/IQTrees1_mask_041124 .treefile.mm 0.50
~/flowchart_1124/IQTrees1_shrink050_041124 outgroup.txt
```

Quantile 0.10 seems to be the best.

Remove those sequences from the original unaligned files that **TreeShrink** removed from the trees:

```
python write_homolog_fasta_from_multiple_aln.py ~/IQTree/04_Scipio_Tamaricaceae_outgroupsNUC_alignments/01_unaligned/01_coding_NUC/03_genes ~/flowchart_1124/IQTrees1_shrink010_041124 .fna .ts ~/flowchart_1124/Unaligned_shrunk_041124
```

---

# E

<u>Align with **MAFFT**</u>

Run the following command in the folder `~/flowchart_1124/Unaligned_shrunk_041124`

Align using **MAFFT** with GNU parallel to optimise multi-threading:

```
parallel mafft --nuc --auto --maxiterate 1000 --reorder {} '>' {}.mafft.aln ::: *.fa
```

Move the output to a separate folder (e.g. `~/flowchart_1124/MAFFT2_041124`)

<u>Trim alignments using **Phyx**</u>

Trim alignments using **Phyx** with Python 2 wrapper:

```
python2 pxclsq_wrapper.py ~/flowchart_1124/MAFFT2_041124 0.3 dna
```

<u>Infer Trees using **IQ-TREE**</u>

Run the following command in the folder `~/flowchart_1124/MAFFT2_041124` .

Induce phylogenetic trees using **IQ-TREE** with GNU parallel (with bootstrap):

```
parallel iqtree2 -m MFP --seqtype DNA -B 1000 -wbtl -s ::: *.aln-cln
```

Move the output to a separate folder (e.g. `~/flowchart_1124/IQTree2_041124`).

with bash script to replace @ removed by **IQ-TREE** by moving the script `IQtreeatperl.sh` to the folder just created and executing the command

```
bash IQtreeatperl.sh
```

<u>Infer tree with **ASTRAL-Pro** from homolog trees</u>

Create a folder for the **ASTRAL-Pro** analysis (e.g. `~/flowchart_1124/ASTRALPro3`).

Adapt **Captus' ASTRAL-Pro** .tsv output to account for the @ notation by using sed to add @paralog-fields (one line at a time) from the folder `~/flowchart_1124/ASTRALPro3`:

```
cp ~/Captus/04_Scipio_Tamaricaceae_outgroups_alignments/astral-pro.tsv  astral-pro-at.tsv

sed -i -e 's/__00/@paralog_00/g' astral-pro-at.tsv

sed -i -e 's/__01/@paralog_01/g' astral-pro-at.tsv
```

```
sed -i -e 's/__02/@paralog_02/g' astral-pro-at.tsv
sed -i -e 's/__03/@paralog_03/g' astral-pro-at.tsv
sed -i -e 's/__04/@paralog_04/g' astral-pro-at.tsv
sed -i -e 's/__05/@paralog_05/g' astral-pro-at.tsv
sed -E 's/^([^\t]+)\t\1$/\1@paralog_unique\t\1/' astral-pro-at.tsv
```

---

# F

Concatenate homolog gene trees by executing the following command :

```
cat IQTree2_041124/*.treefile > homologgenetrees.nw
```

Run **ASTRAL-Pro** from `~/flowchart_1124/ASTRALPro3` :

```
$PATH_to_ASTRAL_Installation$/astral-pro3 -a astral_table.tsv -i IQTree2_041124/homologgenetrees.nw -o astral-pro-tree.tre -t 8 --root Reaumuria_songarica_outgroup1 2> astralpro.log
```

---

# G

<u>Prune orthologs (Monophyletic Outgroup; MO)</u>

Prune orthologs using Python 2 script:

```
python2 prune_paralogs_MO_1to1_MO.py ~/flowchart_1124/IQTree2_041124 .treefile 20 ~/flowchart_1124/IQTree2_MO_041124 ~/flowchart_1124/taxonID.txt
```

Remove those sequences from the original unaligned files that MO removed from the trees:

```
python write_ortholog_fasta_from_multiple_aln.py ~/IQTree/04_Scipio_Tamaricaceae_outgroupsNUC_alignments/01_unaligned/01_coding_NUC/03_genes ~/flowchart_1124/IQTree2_MO_041124 .fna .tre ~/flowchart_1124/unaligned_prunedMO_041124
```

<u>Align with **MAFFT**</u>

Run the following command from `~/flowchart_1124/unaligned_prunedMO_04112`:

Align using **MAFFT** with GNU parallel to optimise multi-threading:

```
parallel mafft --nuc --auto --maxiterate 1000 --reorder {} '>' {}.mafft.aln ::: *.fa
```

Move the output to a separate folder (e.g. `~/flowchart_1124/MAFFT3_041124`)

<u>Trim alignment with **Phyx**</u>

Trim alignments using **Phyx** with Python 2 wrapper:

```
python2 pxclsq_wrapper.py ~/flowchart_1124/MAFFT3_041124 0.3 dna
```

Move the output to a separate folder (e.g. `~/flowchart_1124/final_clean_ortholog_alignments_041124`)

---

# H

<u>Concatenate alignments to infer Supermatrix tree</u>

Concatenate all high quality sequences of the final ortholog alignment:

```
python2 concatenate_matrices_phyx.py ~/flowchart_1124/final_clean_ortholog_alignments_041124 600 70 ~/flowchart_1124/final_trimmed_ortholog_supermatrix_041124
```

Infer Supermatrix tree with **IQ-TREE** by running the following command from `~/flowchart_1124/final_trimmed_ortholog_supermatrix_041124`:

```
iqtree2 -s final_trimmed_ortholog_supermatrix_041124.phy -p final_trimmed_ortholog_supermatrix_041124.model -B 1000 -wbtl -T AUTO
```

---

# I

<u>Infer species tree with **ASTRAL-IV:**</u>

Infer all the gene trees by running the following command from `~/flowchart_1124/final_clean_ortholog_alignments_041124`:

```
parallel iqtree2 -m MFP --seqtype DNA -B 1000 -wbtl -s ::: *.aln-cln
```

Create a folder for the ASTRAL-IV analysis (e.g. `~/flowchart_1124/ASTRAL4_also_in_supermatrix_131124`).

Use sed to extract only the names of the alignments from the model file:

```
sed 's/.*\(\/home[^ ]*aln-cln\).*/\1/' final_trimmed_ortholog_supermatrix_041124.model > included_alignments.txt
```

Use find-and-replace in a text editor to alter this list to a list of gene trees.

Concatenate all gene trees occurring in that list:

```
{ xargs cat < genetrees_also_in_supermatrix.txt ; } > genetrees_also_in_supermatrix.nw
```

Run **ASTRAL-IV** to make species tree from only those orthologs that are also in the Supermatrix alignment:

```
$PATH_to_ASTRAL_IV_Installation$/astral4 -o
astral4_concat_ortholog.stree -i genetrees_also_in_supermatrix.nw -t
30 --root Reaumuria_songarica_outgroup1 2> astral4.log
```

---

#### Calculating site (sCF) and gene concordance (gCF) values

First, calculate the sCF values with **IQ-TREE,** using the species tree calculated with **ASTRAL-IV** (step I) and the Supermatrix alignment (step H):

```
iqtree2 -te
astral_species_tree_071124/astral4_concat_also_in_supermatrix.stree
-s MAFFT3_041124/final_trimmed_ortholog_supermatrix_041124.fa --scfl
100 --prefix scfl_concord -nt 30
```

Then with **IQ-TREE** write gCF to the same tree with sCF values generated from previous step and the gene trees (step I) as input:

```
iqtree2 -te scfl_concord.cf.tree --gcf
ortholog_genetrees_051124/concat_genetrees_in_supermatrix.treefile -
-prefix full_concord
```

---

#### Plastid sequence assembly and tree inference

<u>Loci assembly with **Captus**</u>

Input is the same raw sequence data from Helmholtz Institute as for the nuclear Angiosperms353 loci pipeline. Outgroups (*Reaumuria songarica* and *Tamarix gallica*) was downloaded from GenBank (Sayers et al., 2021) and the relevant genes were also retrieved using the **Captus** pipeline.

Step 1: Clean using **Captus** default settings:

```
captus_assembly clean -r
```

Step 2: Assembly using **Captus** default settings:

```
captus_assembly assemble -r 01_clean_reads
```

Step 3: Extract using PTD-specific settings:

```
captus_assembly extract -a 02_assemblies/ -p SeedPlantsPTD --
ptd_min_coverage 50 -f Outgroup_GenBank_plastid/ -o
03_Scipio_Tamaricaceae_outgroupsPTD_extractions/
```

Step 4: “Align” (collect genes) using PTD-specific settings:

```
captus_assembly align -e
03_Scipio_Tamaricaceae_outgroupsPTD_extractions/ -m PTD -f NT,GE --
filter_method none -o 04_Scipio_Tamaricaceae_outgroupsPTD_alignments
```

This step is necessary to collect the genes in the correct format for downstream analysis with **MAFFT**.

Infer homologue gene trees:

Use output from:

```
~/Captus/04_Scipio_Tamaricaceae_outgroupsPTD_alignments/01_unaligned/02_coding_PTD/03_genes
```

Check **Captus** reports and remove sequences with less than 50% loci retrieval.

Rename these FASTA file headings with @ using `reformatCaptus.sh`:

```
bash reformatCaptus.sh
```

Run **MAFFT** with GNU parallel:

```
parallel mafft --nuc --auto --maxiterate 1000 --reorder {} '>' {}.mafft.aln ::: *.fna
```

Trimming with `python2 pxclsq_wrapper.py`:

```
python2 pxclsq_wrapper.py PTD_flowchart_1124/MAFFT_PTD1 0.1 dna
```

Run **IQ-TREE**:

```
parallel iqtree2 -m MFP --seqtype DNA -s ::: *.aln-cln
```

Reformat @ in FASTA file:

```
bash IQtreeatperl.sh
```

Mask monophyletic and paraphyletic tips:

```
python2 mask_tips_by_taxonID_transcripts.py ~/PTD_flowchart_1124/trees_PTD1 .treefile ~/PTD_flowchart_1124/trimmed_MAFFT_PTD1 y ~/PTD_flowchart_1124/IQTreesPTD1_mask_231124
```

Run **TreeShrink** with setting 0.1, using file called `outgroup.txt` containing the names of non-*Myricaria* specimens included in the analysis, to prevent false positives on **TreeShrink**:

```
python tree_shrink_1.3.9_wrapper.py ~/PTD_flowchart_1124/IQTreesPTD1_mask_231124 .treefile.mm 0.10 ~/PTD_flowchart_1124/IQTreesPTD1_shrink010_231124 outgroup.txt
```

Run `write_homolog_fasta_from_multiple_aln.py` to remove those sequences from the original unaligned files that treeshrink removed from the trees:

```
python write_homolog_fasta_from_multiple_aln.py ~/PTD_flowchart_1124/unaligned_codingPTD_genes/no_lyakh ~/PTD_flowchart_1124/IQTreesPTD1_shrink010_231124 .fna .ts ~/PTD_flowchart_1124/PTD_Unaligned_shrunk_231124
```

Run **MAFFT** (2nd round):

```
parallel mafft --nuc --auto --maxiterate 1000 --reorder {} '>' {}.mafft.aln ::: *.fa
```

Run **Phyx** (2nd round):

```
python2 pxclsq_wrapper.py ~/PTD_flowchart_1124/PTD_MAFFT2_231124 0.3 dna
```

Run **IQ-TREE**:

```
	parallel iqtree2 -m MFP --seqtype DNA -B 1000 -wbtl -s ::: 
*.aln-cln
```

Homologue trees in output folder: `PTD_IQTree2_231124`

Reformat @ in FASTA file:

```
	bash IQtreeatperl.sh
```

Infer tree from homologue trees with **ASTRAL-Pro-3**:

Adapt **Captus**' **ASTRAL-Pro-3** `.tsv` output to account for the @ notation by using sed to add @paralog-fields (do the following one line at a time):

```
	sed -i -e 's/__00/@paralog_00/g' astral-pro-at.tsv

	sed -i -e 's/__01/@paralog_01/g' astral-pro-at.tsv

	sed -i -e 's/__02/@paralog_02/g' astral-pro-at.tsv

	sed -i -e 's/__03/@paralog_03/g' astral-pro-at.tsv

	sed -i -e 's/__04/@paralog_04/g' astral-pro-at.tsv

	sed -i -e 's/__05/@paralog_05/g' astral-pro-at.tsv

	sed -E 's/^([^\t]+)\t\1$/\1@paralog_unique\t\1/' astral-pro-
	at.tsv
```

Concatenate homologue  gene trees:

```
	cat *.treefile > PTD_concat_homolog_trees.nw
```

Run **ASTRAL-Pro-3**:

```
	$PATH_to_ASTRAL_Installation$/astral4 -a astral-pro-at.tsv -I 
PTD_concat_homolog_trees.nw -o astral-pro-tree.tre -t 8 -root 
Reaumuria_songarica_chloroplastgenome 2> astralpro.log
```

output tree is `PTD_astral-pro-tree_231124.tre` in folder `PTD_astral_pro_tree_231124`

Create orthologue alignments:

Run prune monophyletic outgroups:

```
	python2 prune_paralogs_MO_1to1_MO.py 
~/PTD_flowchart_1124/PTD_IQTree2_231124 .treefile 20 
~/PTD_flowchart_1124/PTD_IQTree2_MO_231124 
~/PTD_flowchart_1124/PTDtaxonID.txt
```

Write FASTA file from orthologue again:

To remove those sequences from the original unaligned files that MO removed from the trees, run:

```
	python write_ortholog_fasta_from_multiple_aln.py 
~/PTD_flowchart_1124/unaligned_codingPTD_genes
```

```
~/PTD_flowchart_1124/PTD_IQTree2_MO_231124 .fna .tre
~/PTD_flowchart_1124/PTD_unaligned_prunedMO_231124
```

Run **MAFFT** (3rd round):

```
    parallel mafft --nuc --auto --maxiterate 1000 --reorder {} '>'
{}.mafft.aln ::: *.fa
```

Run **Phyx** (3rd round):

```
    python2 pxclsq_wrapper.py PTD_flowchart_1124/PTD_MAFFT3_231124
0.3 dna
```

Infer supermatrix tree from orthologue alignments:

Concatenate all high quality sequences of the final orthologue alignment:

```
    python2 concatenate_matrices_phyx.py
~/PTD_flowchart_1124/PTD_final_clean_ortholog_alignments_231124/ 10
25
~/PTD_flowchart_1124/PTD_final_trimmed_ortholog_supermatrix_231124
```

Run **IQ-TREE**:

```
    iqtree2 -s PTD_final_trimmed_ortholog_supermatrix_231124.phy -p
PTD_final_trimmed_ortholog_supermatrix_231124.model -B 1000 -wbtl -T
AUTO
```

Look at the **IQ-TREE** stats and remove individuals with low quality sequences (removed those that failed the Chi-Squared composition test with *P*-value 0.01)

Infer species tree from orthologue gene trees with low quality individuals removed:

Run script to split it into gene regions again:

```
    slice_supermatrix.sh
```

Infer species tree with **IQ-TREE**:

```
    parallel iqtree2 -m MFP --seqtype DNA -B 1000 -wbtl -s :::
*.fasta
```

Concatenate orthologue gene trees:

```
    cat *.treefile > PTD_genetrees_also_in_supermatrix.nw
```

Infer species tree from orthologue gene trees also in the supermatrix, with *Reaumuria songarica* as the outgroup for rooting:

```
    $PATH_to_ASTRAL_Installation$/astral4 -o
astral4_concat_PTD_ortholog.stree -i
PTD_genetrees_also_in_supermatrix.nw -t 30 --root
Reaumuria_songarica_chloroplastgenome 2> astral4_PTD.log
```
