## Appendix S3 for "Towards an understanding of the taxonomy of the Eurasian montane riparian genus *Myricaria* (Tamaricaceae) through Angiosperms353 herbariomics"

**Supplementary Materials S3: Scripts and Wrappers used in the pipeline**

These scripts are required for the pipeline to function. Most of these scripts were imported and updated from https://bitbucket.org/dfmoralesb/target_enrichment_orthology/src/master/ . New scripts written for this particular pipeline are indicated.

reformatCaptus.sh

```
for i in $(ls *.fna); do

sed -i -E 's/(>.+)(__)([[:digit:]]+)( .+)/\1@paralog_\3/' $i

sed -i -E 's/(>.+)( \[query=.+)/\1@paralog_unique/' $i

done
```

pxclsq_wrapper.py

```
"""

pxclsq (from phyx) wrapper. adapted from Ya's phyutility_wrapper.py

make sure that pxclsq is in the path

"""


import sys,os

from seq import read_fasta_file


def pxclsq(DIR,alignment,min_col_occup,seqtype,min_chr=10):
        """
        remove columns with occupancy lower than
MIN_COLUMN_OCCUPANCY
        remove seqs shorter than MIN_CHR after filter columns
        """
        if DIR[-1] != "/": DIR += "/"
        cleaned = alignment+"-cln"
        if os.path.exists(DIR+cleaned): return cleaned
        assert alignment.endswith(".aln"),\
                "pxclsq infile "+alignment+" not ends with .aln"
        assert os.stat(DIR+alignment).st_size > 0,
DIR+alignment+"empty"
        assert seqtype == "aa" or seqtype == "dna","Input data type:
dna or aa"
```

```
        if seqtype == "aa":
                cmd = ["pxclsq","-a","-p",str(min_col_occup),"-s",\
                           DIR+alignment,"-o",DIR+alignment+"-pht"]
        else:
                cmd = ["pxclsq","-p",str(min_col_occup),"-s",\
                           DIR+alignment,"-o",DIR+alignment+"-pht"]
        print " ".join(cmd)
        os.system(" ".join(cmd))
        assert os.path.exists(DIR+alignment+"-pht"),"Error pxclsq"

        #remove empty and very short seqs
        outfile = open(DIR+cleaned,"w")
        for s in read_fasta_file(DIR+alignment+"-pht"):
                if len(s.seq.replace("-","")) >= min_chr:
                        outfile.write(s.get_fasta())
        outfile.close()
        os.remove(DIR+alignment+"-pht")
        return cleaned

def main(DIR,min_col_occup,seqtype):
        if DIR[-1] != "/": DIR += "/"
        filecount = 0
        for i in os.listdir(DIR):
                if i.endswith(".aln"):
                        filecount += 1
                        pxclsq(DIR,i,min_col_occup,seqtype)
        assert filecount > 0, "No file end with .aln found in "+DIR


if __name__ == "__main__":
        if len(sys.argv) != 4:
                print "python pxclsq_wrapper.py DIR
min_column_occupancy dna/aa"
```

```
                sys.exit(0)

        DIR,min_col_occup,seqtype = sys.argv[1:]
        main(DIR,min_col_occup,seqtype)
```

IQtreeatperl.sh (new)

```
for i in $(ls *.treefile)
do
perl -p -i -e "s/_paralog/\@paralog/g" $i
done
```

mask tips by taxonID transcripts.py

```
"""
Input is a dir of trees that end with ".tt". Change
INTREE_FILE_ENDING if needed

Mask both mono- and paraphyletic tips that belong to the same taxon
If only mask monophyletic tips, comment out this line:
curroot = paraphyly_masking(curroot,unamb_chrDICT)
Keep the tip that has the most un-ambiguous, well-aligned charactors
in the trimmed alignment
"""

import newick3,phylo3,os,sys
from tree_utils import get_name,remove_kink
from seq import read_fasta_file

def get_clusterID(filename):
        """given a file name return the cluster id"""
        return filename.split(".")[0]+'.'+filename.split(".")[1]

def mask_monophyletic_tips(curroot,unamb_chrDICT):
        going = True
        while going and len(curroot.leaves()) >= 4:
```

```
                going = False
                for node in curroot.iternodes(): #walk through nodes
                        if not node.istip: continue #only look at
tips
                        for sister in node.get_sisters():
                                if sister.istip and
get_name(node.label)==get_name(sister.label): #masking
                                        #print
node.label,unamb_chrDICT[node.label],sister.label,unamb_chrDICT[sist
er.label]
                                        if unamb_chrDICT[node.label]
> unamb_chrDICT[sister.label]:
                                                node =
sister.prune()
                                        else: node = node.prune()
                                        if len(curroot.leaves()) >=
4:
                                                if (node==curroot
and node.nchildren==2) or (node!=curroot and node.nchildren==1):
                                                        node,curroot
= remove_kink(node,curroot)
                                        going = True
                                        break
        return curroot


def mask_paraphyletic_tips(curroot,unamb_chrDICT):
        going = True
        while going and len(curroot.leaves()) >= 4:
                going = False
                for node in curroot.iternodes(): #walk through nodes
                        if not node.istip: continue #only look at
tips
                        parent = node.parent
                        if node == curroot or parent == curroot:
                                continue #no paraphyletic tips for
the root
                        for para in parent.get_sisters():
```

```
                                if para.istip and
get_name(node.label)==get_name(para.label):
                                        if unamb_chrDICT[node.label]
> unamb_chrDICT[para.label]:
                                                node = para.prune()
                                        else: node = node.prune()
                                        if len(curroot.leaves()) >=
4:
                                                if (node==curroot
and node.nchildren==2) or (node!=curroot and node.nchildren==1):
                                                        node,curroot
= remove_kink(node,curroot)
                                        going = True
                                        break
        return curroot

def main(treDIR,intree_file_ending,clnDIR,para,outDIR):
        if treDIR[-1] != "/": treDIR += "/"
        if clnDIR[-1] != "/": clnDIR += "/"
        if outDIR[-1] != "/": outDIR += "/"
        assert para=="y" or para=="n", "mask paraphyletic tips?
(y/n)"
        mask_para = True if para == "y" else False
        filecount = 0

        filematch = {} #key is clusterID, value is the .aln-cln file
        for i in os.listdir(clnDIR):
                if i.endswith(".aln-cln"):
                        clusterID = get_clusterID(i)
                        assert clusterID not in filematch, \
                                "The clusterID "+clusterID+" repeats
in "+clnDIR
                        filematch[clusterID] = i

        for i in os.listdir(treDIR):
                if i.endswith(intree_file_ending):
```

```
                        with open(treDIR+i,"r") as infile:
                                intree =
newick3.parse(infile.readline())
                        print i
                        clusterID = get_clusterID(i)
                        filecount += 1
                        chrDICT = {} #key is seqid, value is number
of unambiguous chrs
                        for s in
read_fasta_file(clnDIR+filematch[clusterID]):
                                for ch in ['-','X',"x","?","*"]:
                                        s.seq = s.seq.replace(ch,"")
#ignore gaps, xs and Xs
                                chrDICT[s.name] = len(s.seq)
                        curroot =
mask_monophyletic_tips(intree,chrDICT)
                        if mask_para: curroot =
mask_paraphyletic_tips(curroot,chrDICT)
                        with open(outDIR+i+".mm","w") as outfile:

outfile.write(newick3.tostring(curroot)+";\n")
        assert filecount > 0, \
                "No file ends with "+intree_file_ending+" found in
"+treDIR


if __name__ == "__main__":
        if len(sys.argv) != 6:
                print "python mask_tips_by_taxonID_transcripts.py
treDIR tree_file_ending aln-clnDIR mask_paraphyletic(y/n) outDIR"
                sys.exit(0)


        treDIR,intree_file_ending,clnDIR,para,outDIR = sys.argv[1:]
        main(treDIR,intree_file_ending,clnDIR,para,outDIR)
```

outgroup.txt (sister taxa of the genus *Myricaria*)

```
Tamarix_gallica_outgroup2
Reaumuria_songarica_outgroup1
```

tree_shrink_1.3.9_wrapper.py

```
"""
Detects and trim abnormally long branches using TreeShrink v1.3.9

TreeShrink must be installed and on path
Phyx (pxrr) must be installed and on path

"""


import sys, os, shutil, glob, re
from Bio import Phylo
import pandas as pd


def trim_exceptions(inDIR,tree_file_ending,q,outDIR,exceptions):

        if os.path.isabs(inDIR) == False: inDIR = 
os.path.abspath(inDIR)
        if inDIR[-1] != "/": inDIR += "/"
        if os.path.isabs(outDIR) == False: outDIR = 
os.path.abspath(outDIR)
        if outDIR[-1] != "/": outDIR += "/"
        if outDIR == ".": outDIR = os.getcwd()


        filecount = 0

        with open(exceptions) as f:
            taxa_to_not_remove = f.read().splitlines()
            print(taxa_to_not_remove)
```

```
        for i in os.listdir(inDIR):
                if i.endswith(tree_file_ending):
                        original_tre = Phylo.read(inDIR+i, "newick")
                        original_tre_tips = []
                        for leaf in original_tre.get_terminals():
original_tre_tips.append(leaf.name)
                        ori_tip_db = pd.DataFrame(original_tre_tips)
                        #print ori_tip_db



                        newlist = []
                        for taxa in taxa_to_not_remove:
                                r = re.compile(str(taxa)+".*")
                                newlist.append(filter(r.match,
original_tre_tips))


                        flatlist = []
                        for sublist in newlist:
                                for item in sublist:
                                        flatlist.append(item)


                        print(flatlist)


                        if len(flatlist) >= 1:


                                exception_file =
outDIR+i+".exceptions.ext"
                                with open(exception_file, "w") as
file:
                                        for item in flatlist:
                                                file.write("%s\n" %
item)
```

```
                                os.path.splitext(outDIR+i)[0]

                                #print os.path.splitext(outDIR+i)[0]

                                filecount += 1

                                cmd= ["run_treeshrink.py","-t",
inDIR+i ,"-c","-m per-gene", "-q "+str(q), "-o",outDIR+i+".ts_dir",
"-O "+os.path.splitext(outDIR+i)[0], "-x",exception_file]

                                print(" ".join(cmd))

                                os.system(" ".join(cmd))



                        else:


                                os.path.splitext(outDIR+i)[0]

                                #print os.path.splitext(outDIR+i)[0]

                                filecount += 1

                                cmd= ["run_treeshrink.py","-t",
inDIR+i ,"-c","-m per-gene", "-q "+str(q), "-o",outDIR+i+".ts_dir",
"-O "+os.path.splitext(outDIR+i)[0]]

                                print(" ".join(cmd))

                                os.system(" ".join(cmd))



        #moves output files to DIR and delete treeshrink individual
folders

        for j in os.listdir(outDIR):

                if j.endswith(".ts_dir"):

                        source = outDIR+j

                        dest = outDIR

                        files = os.listdir(source)

                        for f in files:

                                shutil.move(source+"/"+f, dest)

                        shutil.rmtree(outDIR+j)
```

```
        #removes single quotes from tip labels from treeshrink
output trees
        for k in os.listdir(outDIR):
                if k.endswith(".mm"):
                        with open(outDIR+k, 'r+') as f:
                                content = f.read()
                                f.seek(0)
                                f.truncate()
                                f.write(content.replace("'", ""))
                        f.close()



        #removes remove root edge (if present). This avoid issues
with ortholog prunning scripts.les
        for n in os.listdir(outDIR):
                if n.endswith(".mm"):
                        cmd= ["pxcltr","-r","-t", outDIR+n,"-
o",outDIR+n+".tr"]
                        #print(" ".join(cmd))
                        os.system(" ".join(cmd))



        #unroot treeshrink ouput trees
        for l in os.listdir(outDIR):
                if l.endswith(".tr"):
                        cmd= ["pxrr","-u","-t", outDIR+l,"-
o",outDIR+l+".ts"]
                        #print(" ".join(cmd))
                        os.system(" ".join(cmd))



        #delete tr files
        for m in os.listdir(outDIR):
                        if m.endswith(".tr"):
                                os.remove(outDIR+m)
```

```
        #delete ts files
        for m in os.listdir(outDIR):
                        if m.endswith(".mm"):
                                os.remove(outDIR+m)

        #delete extension files
        for o in os.listdir(outDIR):
                        if o.endswith(".ext"):
                                os.remove(outDIR+o)

        #delete summary files
        for p in os.listdir(outDIR):
                        if p.endswith("treefile_summary.txt"):
                                os.remove(outDIR+p)



def trim(inDIR,tree_file_ending,q,outDIR):

        if os.path.isabs(inDIR) == False: inDIR =
os.path.abspath(inDIR)
        if inDIR[-1] != "/": inDIR += "/"
        if os.path.isabs(outDIR) == False: outDIR =
os.path.abspath(outDIR)
        if outDIR[-1] != "/": outDIR += "/"
        if outDIR == ".": outDIR = os.getcwd()


        filecount = 0


        for i in os.listdir(inDIR):
                if i.endswith(tree_file_ending):
```

```
                        os.path.splitext(outDIR+i)[0]

                        #print os.path.splitext(outDIR+i)[0]

                        filecount += 1

                        cmd= ["run_treeshrink.py","-t", inDIR+i ,"-
c","-m per-gene", "-q "+str(q), "-o",outDIR+i+".ts_dir", "-O
"+os.path.splitext(outDIR+i)[0]]

                        print(" ".join(cmd))

                        os.system(" ".join(cmd))



        #moves output files to DIR and delete treeshrink individual
folders

        for j in os.listdir(outDIR):

                if j.endswith(".ts_dir"):

                        source = outDIR+j

                        dest = outDIR

                        files = os.listdir(source)

                        for f in files:

                                shutil.move(source+"/"+f, dest)

                        shutil.rmtree(outDIR+j)



        #removes single quotes from tip labels from treeshrink
output trees

        for k in os.listdir(outDIR):

                if k.endswith(".mm"):

                        with open(outDIR+k, 'r+') as f:

                                content = f.read()

                                f.seek(0)

                                f.truncate()

                                f.write(content.replace("'", ""))

                        f.close()
```

```
        #removes remove root edge (if present). This avoid issues
with ortholog prunning scripts.les

        for n in os.listdir(outDIR):

                if n.endswith(".mm"):

                        cmd= ["pxcltr","-r","-t", outDIR+n,"-
o",outDIR+n+".tr"]

                        #print(" ".join(cmd))

                        os.system(" ".join(cmd))



        #unroot treeshrink ouput trees

        for l in os.listdir(outDIR):

                if l.endswith(".tr"):

                        cmd= ["pxrr","-u","-t", outDIR+l,"-
o",outDIR+l+".ts"]

                        #print(" ".join(cmd))

                        os.system(" ".join(cmd))



        #delete tr files

        for m in os.listdir(outDIR):

                        if m.endswith(".tr"):

                                os.remove(outDIR+m)


        #delete ts files

        for m in os.listdir(outDIR):

                        if m.endswith(".mm"):

                                os.remove(outDIR+m)



        #delete summary files

        for p in os.listdir(outDIR):

                        if p.endswith("treefile_summary.txt"):

                                os.remove(outDIR+p)
```

```
        assert filecount > 0, \
                "No file end with "+tree_file_ending+" found in 
"+inDIR


if __name__ == "__main__":


        if len(sys.argv) == 6:
                inDIR,tree_file_ending,q,outDIR,exceptions = 
sys.argv[1:]

trim_exceptions(inDIR,tree_file_ending,q,outDIR,exceptions)


        elif len(sys.argv) == 5:
                inDIR,tree_file_ending,q,outDIR = sys.argv[1:]
                trim(inDIR,tree_file_ending,q,outDIR)


        else:
                print("Usage:")
                print("python tree_shrink_wrapper.py inDIR 
tree_file_ending quantile outDIR exceptions(optional)")
                sys.exit(0)
```

write homolog fasta from multiple aln.py

```
"""
This script take a folder of fasta files and
write fasta files given a folder of trees
"""


import os, sys, glob
from Bio import Phylo, SeqIO


def write_fasta(fasta_DIR, tree_DIR, fasta_file_ending,
tree_file_ending, outDIR):
	""""Takes a fasta file and tree and write fasta file only
the taxa from the tree"""


	if os.path.isabs(tree_DIR) == False: tree_DIR =
os.path.abspath(tree_DIR)
	if tree_DIR[-1] != "/": tree_DIR += "/"


	if os.path.isabs(fasta_DIR) == False: fasta_DIR =
os.path.abspath(fasta_DIR)
	if fasta_DIR[-1] != "/": fasta_DIR += "/"


	if os.path.isabs(outDIR) == False: outDIR =
os.path.abspath(outDIR)
	if outDIR[-1] != "/": outDIR += "/"
	if outDIR == ".": outDIR = os.getcwd()


	filecount = 0



	for i in os.listdir(tree_DIR):
		if i.endswith(tree_file_ending):
			print(i)
			filecount += 1
```

```
                        original_tre = Phylo.read(tree_DIR+i, 
"newick")
                        original_tre_tips = []
                        for leaf in original_tre.get_terminals(): 
original_tre_tips.append(leaf.name)
                        print(original_tre_tips)


                        path_tree, files_tree = os.path.split(i)
                        tree_name = str(files_tree)
                        base_name_tree = tree_name.split( "." )
                        print(base_name_tree)


                        fasta_file = glob.glob(fasta_DIR + 
base_name_tree[0]+ fasta_file_ending)
#"."+base_name_tree[1] +
                        print(fasta_file)


                        new_fasta_file = 
outDIR+base_name_tree[0]+"."+base_name_tree[1]+".homolog.fa"
                        #print(new_fasta_file)
                        file=open(new_fasta_file,'w')


                        for record in 
SeqIO.parse(fasta_file[0],"fasta"):
                            if record.id in original_tre_tips:
                                id =record.id
                                seq=str(record.seq)
                                file.write(">" + id + "\n" + seq + 
"\n")
                        file.close()


if __name__ == "__main__":
        if len(sys.argv) != 6:
                print("python 
write_ortholog_fasta_from_multiple_aln.py fasta_DIR tree_DIR 
fasta_file_ending(no_dot) tree_file_ending(not_dot) outDIR")
```

```
                sys.exit(0)

        fasta_DIR, tree_DIR, fasta_file_ending, tree_file_ending, outDIR = sys.argv[1:]
        write_fasta(fasta_DIR, tree_DIR, fasta_file_ending, tree_file_ending, outDIR)
```

prune_paralogs_MO_1to1_MO.py

```
"""
Prepare a taxon file, with each line look like (separated by tab):
IN	taxonID1
IN	taxonID2
OUT	taxonID3

Taxon duplication? --No--> output one-to-one orthologs
                |
            Yes
             |
Outgroup present? --No--> ignore this homolog
                |
            Yes
             |
Outgroup taxon duplication? --Yes--> ignore this homolog
                |
             No
             |
Outgroup monophyletic? --No--> ignore this homolog
                |
            Yes
             |
Infer orthologs by using monophyletic, non-repeating outgroups

If not to output 1-to-1 orthologs, for example, already analysed these
```

```
set OUTPUT_1to1_ORTHOLOGS to False
"""

import phylo3,newick3,os,sys

OUTPUT_1to1_ORTHOLOGS = True

#if pattern changes, change it here
#given tip label, return taxon name identifier
def get_name(label):
	return label.split("@")[0]

def get_clusterID(filename):
	return filename.split(".")[0]+"."+filename.split(".")[1]

def get_front_labels(node):
	leaves = node.leaves()
	return [i.label for i in leaves]

def get_back_labels(node,root):
	all_labels = get_front_labels(root)
	front_labels = get_front_labels(node)
	return set(all_labels) - set(front_labels) #labels do not 
repeat

def get_front_names(node): #may include duplicates
	labels = get_front_labels(node)
	return [get_name(i) for i in labels]

def get_front_outgroup_names(node):
	names = get_front_names(node)
	return [i for i in names if i in OUTGROUPS]

def get_back_names(node,root): #may include duplicates
```

```
		back_labels = get_back_labels(node,root)
		return [get_name(i) for i in back_labels]


def remove_kink(node,curroot):
		if node == curroot and curroot.nchildren == 2:
				#move the root away to an adjacent none-tip
				if curroot.children[0].istip: #the other child is not tip
						curroot = phylo3.reroot(curroot,curroot.children[1])
				else: curroot = phylo3.reroot(curroot,curroot.children[0])
		#---node---< all nodes should have one child only now
		length = node.length + (node.children[0]).length
		par = node.parent
		kink = node
		node = node.children[0]
		#parent--kink---node<
		par.remove_child(kink)
		par.add_child(node)
		node.length = length
		return node,curroot


#check if outgroups are monophyletic and non-repeating and reroot
#otherwise return None
def reroot_with_monophyletic_outgroups(root):
		lvs = root.leaves()
		outgroup_matches = {} #key is label, value is the tip node object
		#Since no taxon repeat in outgroups name and leaf is one-to-one
		outgroup_labels = []
		for leaf in lvs:
				label = leaf.label
				name = get_name(label)
```

```
		if name in OUTGROUPS:
			outgroup_matches[label] = leaf
			outgroup_labels.append(leaf.label)
	if len(outgroup_labels) == 1: #one single outgroup
		#cannot reroot on a tip so have to go one more node into the ingroup
		new_root = outgroup_matches[outgroup_labels[0]].parent
		return phylo3.reroot(root,new_root)
	else: #has multiple outgroups. Check monophyly and reroot
		newroot = None
		for node in root.iternodes():
			if node == root: continue #skip the root
			front_names = get_front_names(node)
			back_names = get_back_names(node,root)
			front_in_names,front_out_names,back_in_names,back_out_names = 0,0,0,0
			for i in front_names:
				if i in OUTGROUPS: front_out_names += 1
				else: front_in_names += 1
			for j in back_names:
				if j in OUTGROUPS: back_out_names += 1
				else: back_in_names += 1
			if front_in_names==0 and front_out_names>0 and back_in_names>0 and back_out_names==0:
				newroot = node #ingroup at back, outgroup in front
				break
			if front_in_names>0 and front_out_names==0 and back_in_names==0 and back_out_names>0:
				newroot = node.parent #ingroup in front, outgroup at back
				break
		if newroot != None:
```

```
                        return phylo3.reroot(root,newroot)
                else: return None



def prune_paralogs_from_rerooted_homotree(root):
        if len(get_front_names(root)) ==
len(set(get_front_names(root))):
                return root #no pruning needed
        #check for duplications at the root first
        #one or two of the trifurcating root clades are ingroup
clades
        node0,node1,node2 =
root.children[0],root.children[1],root.children[2]

out0,out1,out2=len(get_front_outgroup_names(node0)),len(get_front_ou
tgroup_names(node1)),len(get_front_outgroup_names(node2))
        if out0 == 0 and out1 == 0: #0 and 1 are the ingroup clades
                name_set0 = set(get_front_names(node0))
                name_set1 = set(get_front_names(node1))
                if len(name_set0.intersection(name_set1)) > 0:
                        if len(name_set0) > len(name_set1): #cut the
side with less taxa
                                root.remove_child(node1)
                                node1.prune()
                        else:
                                root.remove_child(node0)
                                node0.prune()
        elif out1 == 0 and out2 == 0: #1 and 2 are the ingroup
clades
                name_set1 = set(get_front_names(node1))
                name_set2 = set(get_front_names(node2))
                if len(name_set1.intersection(name_set2)) > 0:
                        if len(name_set1) > len(name_set2): #cut the
side with less taxa
                                root.remove_child(node2)
                                node2.prune()
```

```
                        else:
                                root.remove_child(node1)
                                node1.prune()
        elif out0 == 0 and out2 == 0: #0 and 2 are the ingroup clades
                name_set0 = set(get_front_names(node0))
                name_set2 = set(get_front_names(node2))
                if len(name_set0.intersection(name_set2)) > 0:
                        if len(name_set0) > len(name_set2): #cut the side with less taxa
                                root.remove_child(node2)
                                node2.prune()
                        else:
                                root.remove_child(node0)
                                node0.prune()
        while len(get_front_names(root)) > len(set(get_front_names(root))):
                for node in root.iternodes(order=0): #PREORDER, root to tip
                        if node.istip or node == root: continue
                        child0,child1 = node.children[0],node.children[1]
                        name_set0 = set(get_front_names(child0))
                        name_set1 = set(get_front_names(child1))
                        if len(name_set0.intersection(name_set1)) > 0:
                                if len(name_set0) > len(name_set1): #cut the side with less taxa
                                        node.remove_child(child1)
                                        child1.prune()
                                else:
                                        node.remove_child(child0)
                                        child0.prune()
                                node,root = remove_kink(node,root) #no rerooting here
                                break
```

```
        return root


if __name__ == "__main__":
        if len(sys.argv) != 6:
                print "python prune_paralogs_MO.py homoTreeDIR 
tree_file_ending minimal_taxa outDIR taxon_file"
                sys.exit(0)


        inDIR = sys.argv[1]+"/"
        tree_file_ending = sys.argv[2]
        MIN_TAXA = int(sys.argv[3])
        outDIR = sys.argv[4]+"/"
        taxon_file = sys.argv[5]


        INGROUPS = []
        OUTGROUPS = []
        with open(taxon_file,"r") as infile:
                for line in infile:
                        if len(line) < 3: continue
                        spls = line.strip().split("\t")
                        if spls[0] == "IN": INGROUPS.append(spls[1])
                        elif spls[0] == "OUT": 
OUTGROUPS.append(spls[1])
                        else:
                                print "Check taxon_code_file file 
format"
                                sys.exit()
        if len(set(INGROUPS) & set(OUTGROUPS)) > 0:
                print "Taxon ID",set(INGROUPS) & set(OUTGROUPS),"in 
both ingroups and outgroups"
                sys.exit(0)
        print len(INGROUPS),"ingroup taxa 
and",len(OUTGROUPS),"outgroup taxa read"
        print "Ingroups:",INGROUPS
        print "Outgroups:",OUTGROUPS
```

```
	for i in os.listdir(inDIR):
		if not i.endswith(tree_file_ending): continue
		print i

		#read in the tree and check number of taxa
		outID = outDIR+get_clusterID(i)
		with open(inDIR+i,"r") as infile:
			 intree = newick3.parse(infile.readline())
		curroot = intree
		names = get_front_names(curroot)
		num_tips,num_taxa = len(names),len(set(names))
		if num_taxa < MIN_TAXA:
			continue #not enough taxa

		#If the homolog has no taxon duplication, no cutting 
is needed
		if num_tips == num_taxa:
			if OUTPUT_1to1_ORTHOLOGS:
				curroot = 
reroot_with_monophyletic_outgroups(curroot)
				if curroot != None:
					os.system("cp "+inDIR+i+" 
"+outID+".1to1ortho.tre")
				else: print "outgroup non-
monophyletic in 1_to_1"
		else:
			#now need to deal with taxon duplications
			#check to make sure that the ingroup and 
outgroup names were set correctly
			for name in names:
				if name not in INGROUPS and name not 
in OUTGROUPS:
					print "check name",name
					print INGROUPS
```

```
                                        print OUTGROUPS
                                        sys.exit()
                        outgroup_names =
get_front_outgroup_names(curroot)

                        #if no outgroup at all, do not attempt to
resolve gene duplication
                        if len(outgroup_names) == 0:
                                print "duplicated taxa in unrooted
tree"

                        #skip the homolog if there are duplicated
outgroup taxa
                        elif len(outgroup_names) >
len(set(outgroup_names)):
                                print "outgroup contains taxon
repeats"

                        else: #at least one outgroup present and
there's no outgroup duplication
                                if curroot.nchildren == 2: #need to
reroot
                                        temp,curroot =
remove_kink(curroot,curroot)
                                curroot =
reroot_with_monophyletic_outgroups(curroot)
                                #only return one tree after prunning
                                if curroot != None:
                                        #with
open(outID+".reroot","w") as outfile:
                                        #
outfile.write(newick3.tostring(curroot)+";\n")
                                        ortho =
prune_paralogs_from_rerooted_homotree(curroot)
                                        if
len(set(get_front_names(curroot))) >= MIN_TAXA:
                                                with
open(outID+".ortho.tre","w") as outfile:
```

```
outfile.write(newick3.tostring(ortho)+";\n")
                                        else: print "not enough taxa
after pruning"
                                else: print "outgroup non-
monophyletic"
```

write_ortholog_fasta_from_multiple_aln.py

```
"""
This script take a folder of fasta files and
write fasta files given a folder of trees
"""

import os, sys, glob
from Bio import Phylo, SeqIO

def write_fasta(fasta_DIR, tree_DIR, fasta_file_ending,
tree_file_ending, outDIR):
        """"Takes a fasta file and tree and write fasta file only
the taxa from the tree"""

        if os.path.isabs(tree_DIR) == False: tree_DIR =
os.path.abspath(tree_DIR)
        if tree_DIR[-1] != "/": tree_DIR += "/"

        if os.path.isabs(fasta_DIR) == False: fasta_DIR =
os.path.abspath(fasta_DIR)
        if fasta_DIR[-1] != "/": fasta_DIR += "/"

        if os.path.isabs(outDIR) == False: outDIR =
os.path.abspath(outDIR)
        if outDIR[-1] != "/": outDIR += "/"
        if outDIR == ".": outDIR = os.getcwd()

        filecount = 0
```

```
	for i in os.listdir(tree_DIR):
		if i.endswith(tree_file_ending):
			print(i)
			filecount += 1
			original_tre = Phylo.read(tree_DIR+i,
"newick")
			original_tre_tips = []
			for leaf in original_tre.get_terminals():
original_tre_tips.append(leaf.name)
			#print(original_tre_tips)

			path_tree, files_tree = os.path.split(i)
			tree_name = str(files_tree)
			base_name_tree = tree_name.split( "." )
			#print(base_name_tree)

			fasta_file = glob.glob(fasta_DIR +
base_name_tree[0]+ fasta_file_ending)
			#print(fasta_file)

			new_fasta_file =
outDIR+base_name_tree[0]+"."+base_name_tree[1]+".ortho.fa"
			#print(new_fasta_file)
			file=open(new_fasta_file,'w')

			for record in
SeqIO.parse(fasta_file[0],"fasta"):
			    if record.id in original_tre_tips:
			        id =record.id
			        species_id,copy_id = id.split("@")
			        seq=str(record.seq)
			        file.write(">" + species_id + "\n" +
seq + "\n")
			file.close()
```

```
if __name__ == "__main__":
        if len(sys.argv) != 6:
                print("python write_ortholog_fasta_from_multiple_aln.py fasta_DIR tree_DIR fasta_file_ending(no_dot) tree_file_ending(not_dot) outDIR")
                sys.exit(0)

        fasta_DIR, tree_DIR, fasta_file_ending, tree_file_ending, outDIR = sys.argv[1:]
        write_fasta(fasta_DIR, tree_DIR, fasta_file_ending, tree_file_ending, outDIR)
```

concatenate_matrices_phyx.py

```
"""
Filter the aligned and cleaned ortholog matrices by number of taxa and characters
Write out name of matrices that passed the filter
Also write out supermatrix stats
"""
from seq import read_fasta_file
from tree_utils import get_name
import sys,os

def concatenate(clnDIR,numofsitesFilter,numoftaxaFilter,outname):
        """filter cleaned alignments and concatenate"""
        if clnDIR[-1] != "/": clnDIR += "/"
        sites_filter = int(numofsitesFilter)
        taxa_filter = int(numoftaxaFilter)

        print "Filtering ortholog matrixes"
        selected = [] # list of alignment file names that pass the filters
        for i in os.listdir(clnDIR):
                if i.endswith(".aln-cln"):
```

```
                        print i
                        seqlist = read_fasta_file(clnDIR+i)
                        num_seq = len(seqlist)
                        num_col = len(seqlist[0].seq)
                        if num_seq >= taxa_filter and num_col >=
sites_filter:
                                selected.append(i)
        print len(selected),"matrices passed the filter"


        print "Getting matrix occupancy stats"
        taxon_occupancy = {}
        #key is taxon name, value is [times present in a
matrix,total length for this taxon]
        total_aligned_len = 0 #record how long the final
concatenated matrix is


        cmd = "pxcat"+" -o "+outname+".fa"+" -p "+outname+".model"+"
-s "
        for i in selected:
                cmd += clnDIR+i+" "
                seqlist = read_fasta_file(clnDIR+i)
                total_aligned_len += len(seqlist[0].seq)
                for s in seqlist:
                        taxonid = get_name(s.name)
                        if taxonid not in taxon_occupancy:
                                taxon_occupancy[taxonid] = [0,0]
                        taxon_occupancy[taxonid][0] += 1
                        taxon_occupancy[taxonid][1] +=
len((s.seq.replace("-","")).replace("?",""))
        cmd += "\n"


        total_ortho = len(selected)
        with open(outname+"_taxon_occupancy_stats","w") as outfile:

outfile.write("taxon\t#orthologs\t#total_charactors\tperc_orthologs\
tperc_charactors\n")
```

```
                sum_char = 0
                for taxon in taxon_occupancy:
                        times,chars =
taxon_occupancy[taxon][0],taxon_occupancy[taxon][1]
                        sum_char += chars
                        out =
taxon+"\t"+str(times)+"\t"+str(chars)+"\t"
                        out +=
str(times/float(total_ortho))+"\t"+str(chars/float(total_aligned_len
))+"\n"
                        outfile.write(out)
                total_taxa = len(taxon_occupancy)
                out = "\nSupermatrix dimension "+str(total_taxa)+"
taxa, "
                out += str(total_ortho)+" loci and
"+str(total_aligned_len)+" aligned columns\n"
                out += "Overall matrix occupancy
"+str(sum_char/float(total_taxa*total_aligned_len))+"\n"
                outfile.write(out)


        print "Supermatrix taxon occupancy stats written
to",outname+"_taxon_occupancy_stats"
        print "Waiting for concatenation to finish. This may take
several minutes..."
        with open(outname+".temp.sh","w") as f: f.write(cmd)
        os.system("bash "+outname+".temp.sh")


        #writes phy file
        cmd_pxs2phy = ["pxs2phy","-o",outname+".phy","-
s",outname+".fa"]
        print (" ".join(cmd_pxs2phy))
        os.system(" ".join(cmd_pxs2phy))


        #writes nex file
        cmd_pxs2nex = ["pxs2nex","-o",outname+".nex","-
s",outname+".fa"]
        print (" ".join(cmd_pxs2nex))
```

```
        os.system(" ".join(cmd_pxs2nex))


        assert os.path.exists(outname+".phy") and 
os.path.exists(outname+".nex") and os.path.exists(outname+".fa"), 
"error concatenate"
        os.system("rm "+outname+".temp.sh")
        print "outfiles written",outname+".phy",outname+".model"


if __name__ == "__main__":
        if len(sys.argv) != 5:
                print "usage: python concatenate_matrices_phix.py 
aln-clnDIR numofsitesFilter numoftaxaFilter outname"
                sys.exit()


        clnDIR,numofsitesFilter,numoftaxaFilter,outname = 
sys.argv[1:]
        concatenate(clnDIR,numofsitesFilter,numoftaxaFilter,outname)
```

slice_supermatrix.sh

```
#!/bin/bash

# Check if both arguments are provided
if [ "$#" -ne 2 ]; then
    echo "Usage: $0 <master_alignment_file> <partition_file>"
    echo "Example: $0 supermatrix.fasta partitions.txt"
    exit 1
fi

INPUT_ALIGN="$1"
PARTITION_FILE="$2"
OUT_FORMAT="fasta" # Output format: fasta, nexus, phylip, etc.

# Verify both input files actually exist
if [ ! -f "$INPUT_ALIGN" ]; then
    echo "Error: Master alignment file '$INPUT_ALIGN' not found."
```

```
    exit 1
fi

if [ ! -f "$PARTITION_FILE" ]; then
    echo "Error: Partition file '$PARTITION_FILE' not found."
    exit 1
fi

echo "Starting SeaView alignment slicing..."

# Read the partition file line by line
while IFS= read -r line || [ -n "$line" ]; do
    # Skip empty lines
    [[ -z "${line// }" ]] && continue

    # Parse out the coordinate range (everything after the '=' sign)
    COORDINATES=$(echo "$line" | cut -d'=' -f2 | tr -d '[:space:]')

    # Extract the file path/name from the left side of the '=' sign
    RAW_PATH=$(echo "$line" | cut -d'=' -f1 | cut -d',' -f2 | tr -d
'[:space:]')

    # Strip paths to get a clean base filename
    GENE_NAME=$(basename "$RAW_PATH")

    # Set output name
    OUTPUT_FILE="${GENE_NAME}.${OUT_FORMAT}"

    echo "Slicing ${GENE_NAME} [Sites: ${COORDINATES}] ->
${OUTPUT_FILE}"

    # Run seaview headless to isolate the site selection
    seaview -convert "$OUT_FORMAT" \
            -def_site_selection "$GENE_NAME","$COORDINATES" \
```

```
            -sites "$GENE_NAME" \
            -o "$OUTPUT_FILE" \
            "$INPUT_ALIGN"

done < "$PARTITION_FILE"

echo "All partitions successfully extracted!"
```

outgroup.txt

```
Tamarix_gallica_outgroup2
Reaumuria_songarica_outgroup1
```
