## Supplementary figures and images for "Towards an understanding of the taxonomy of the Eurasian montane riparian genus *Myricaria* (Tamaricaceae) through Angiosperms353 herbariomics"

### Figure S1

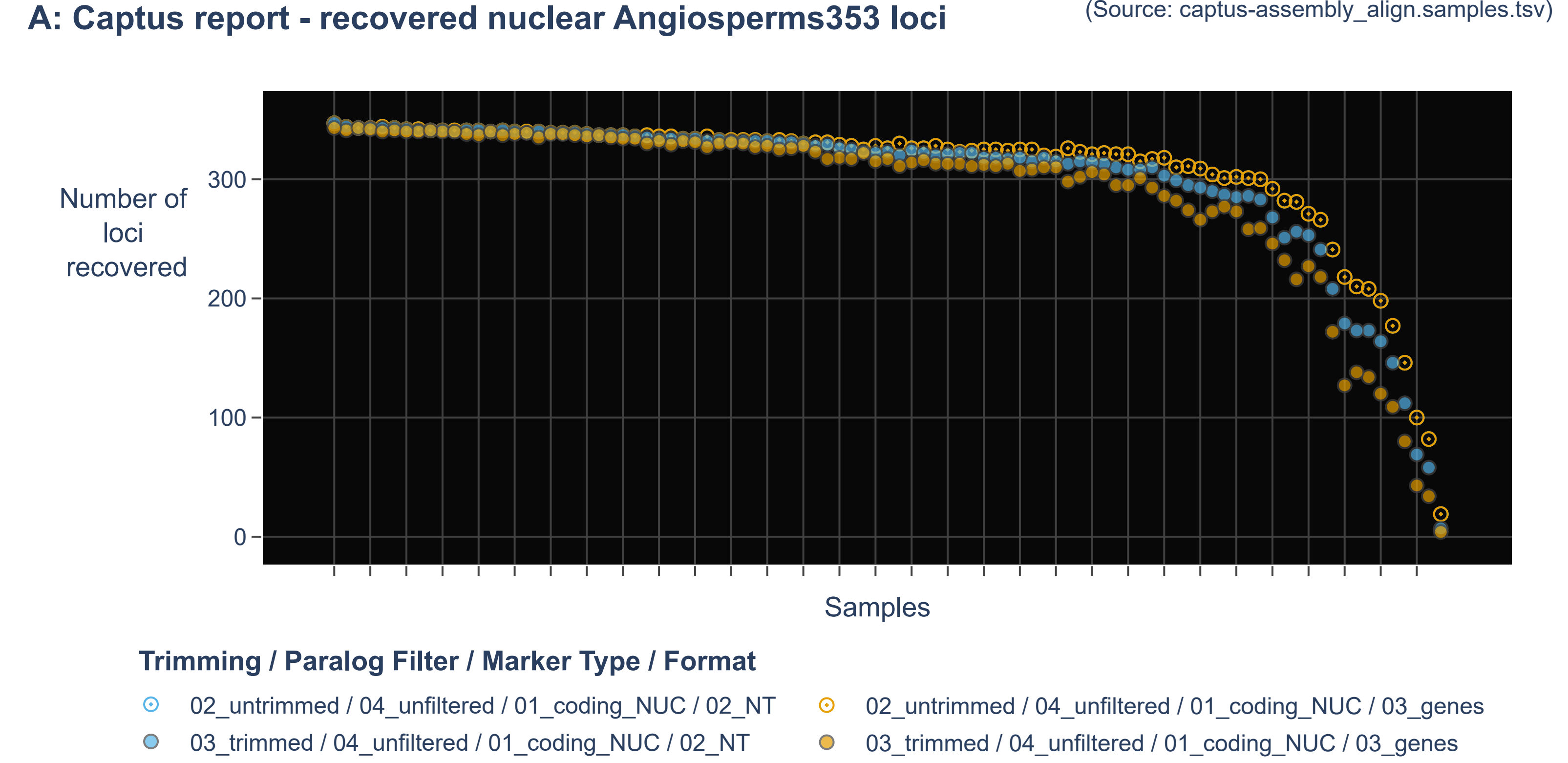

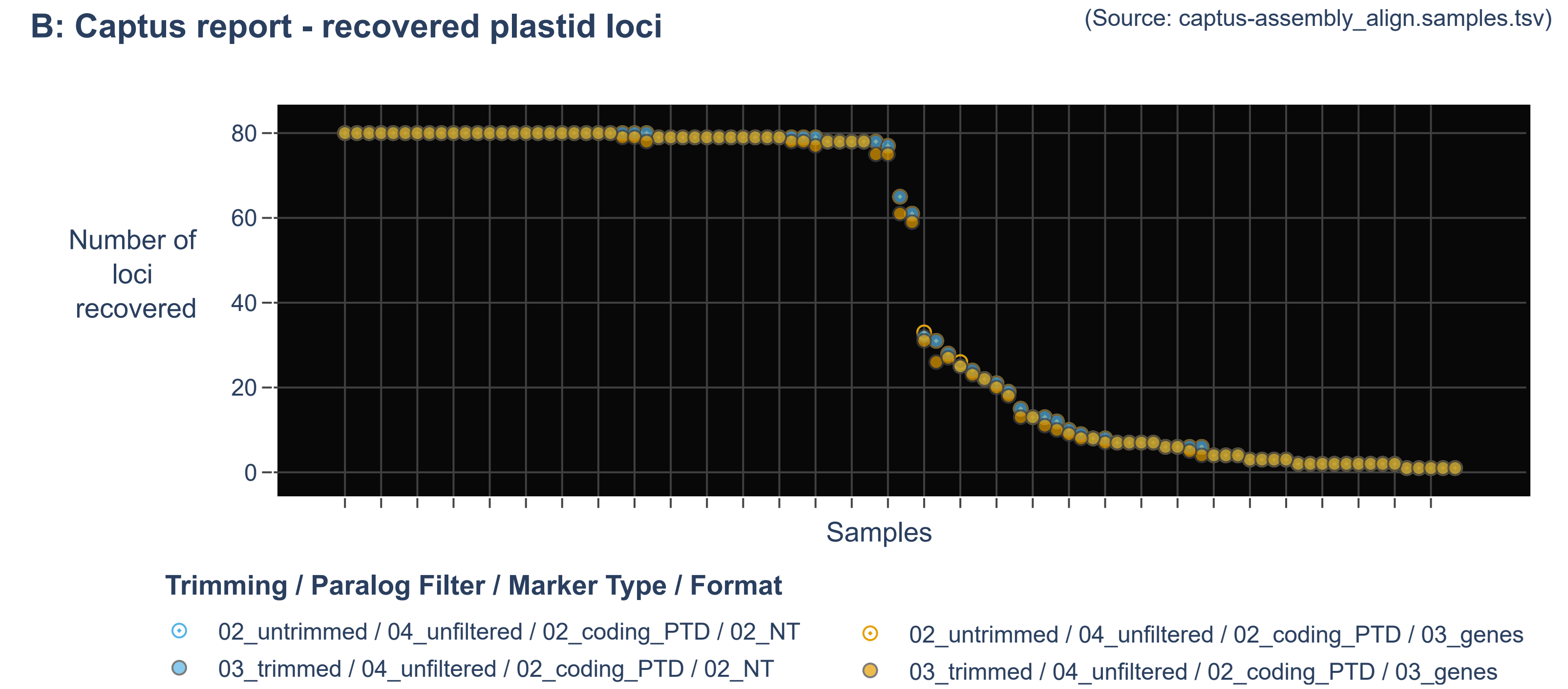

### Figure S2

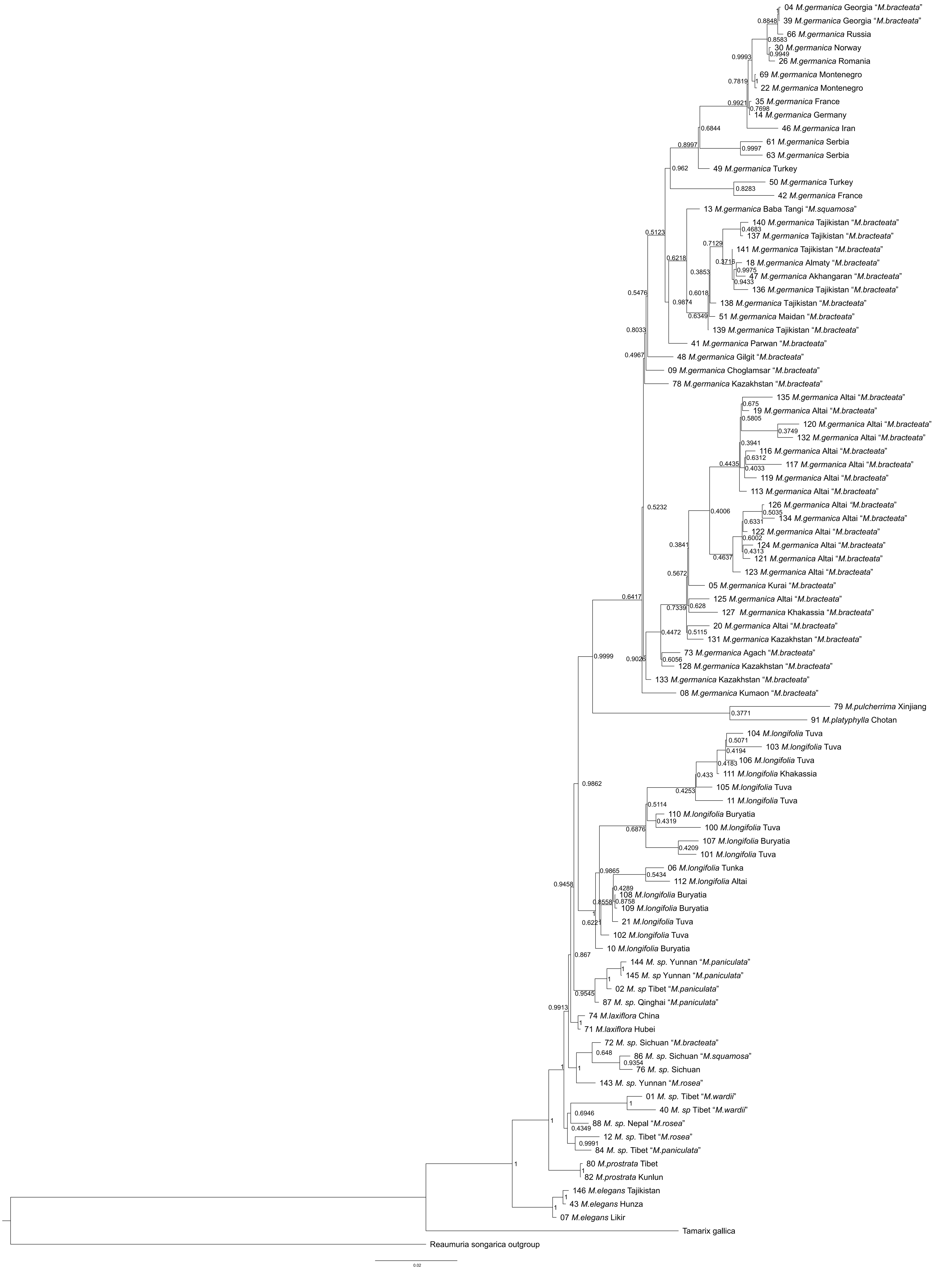

### Figure S3

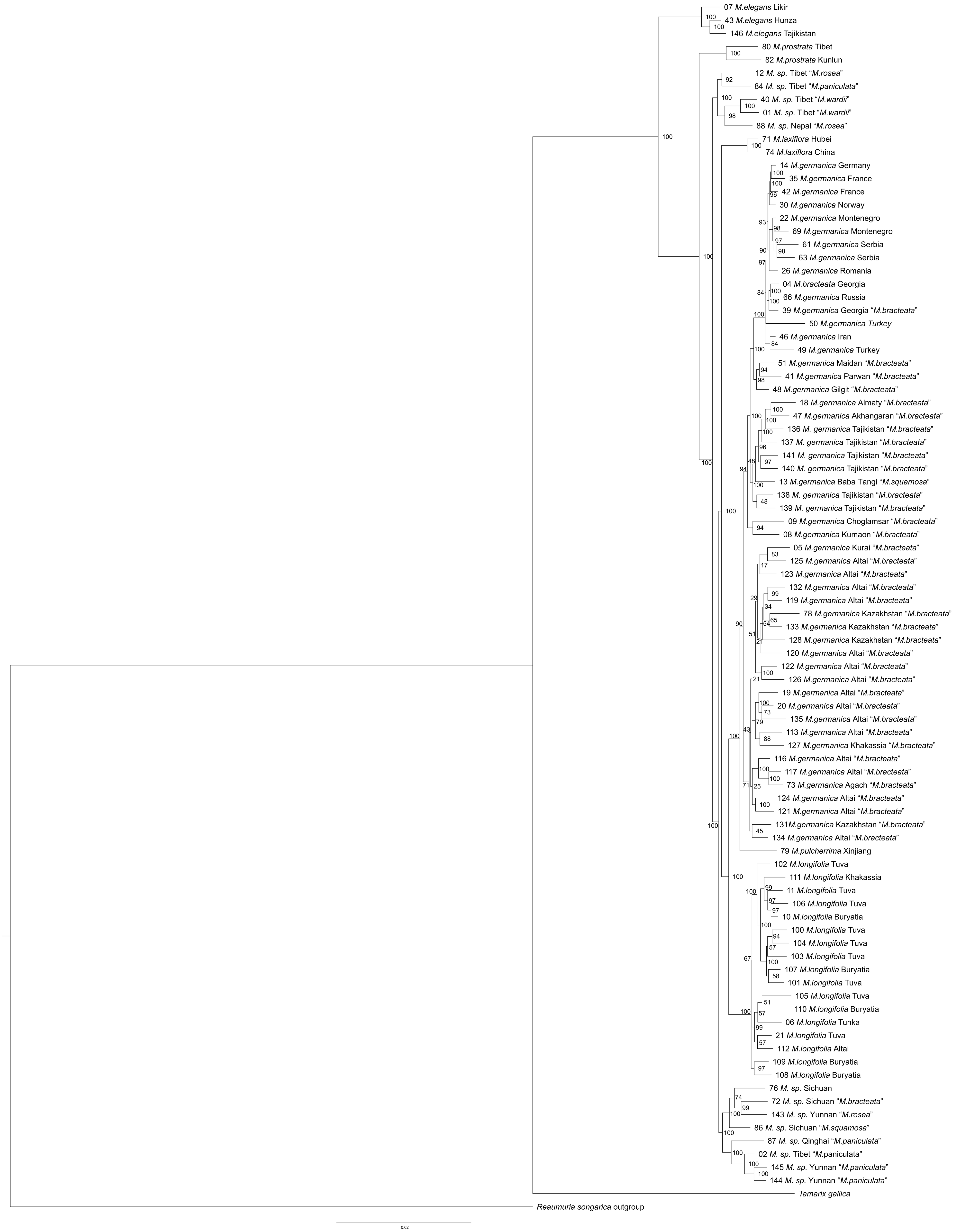

### Figure S4

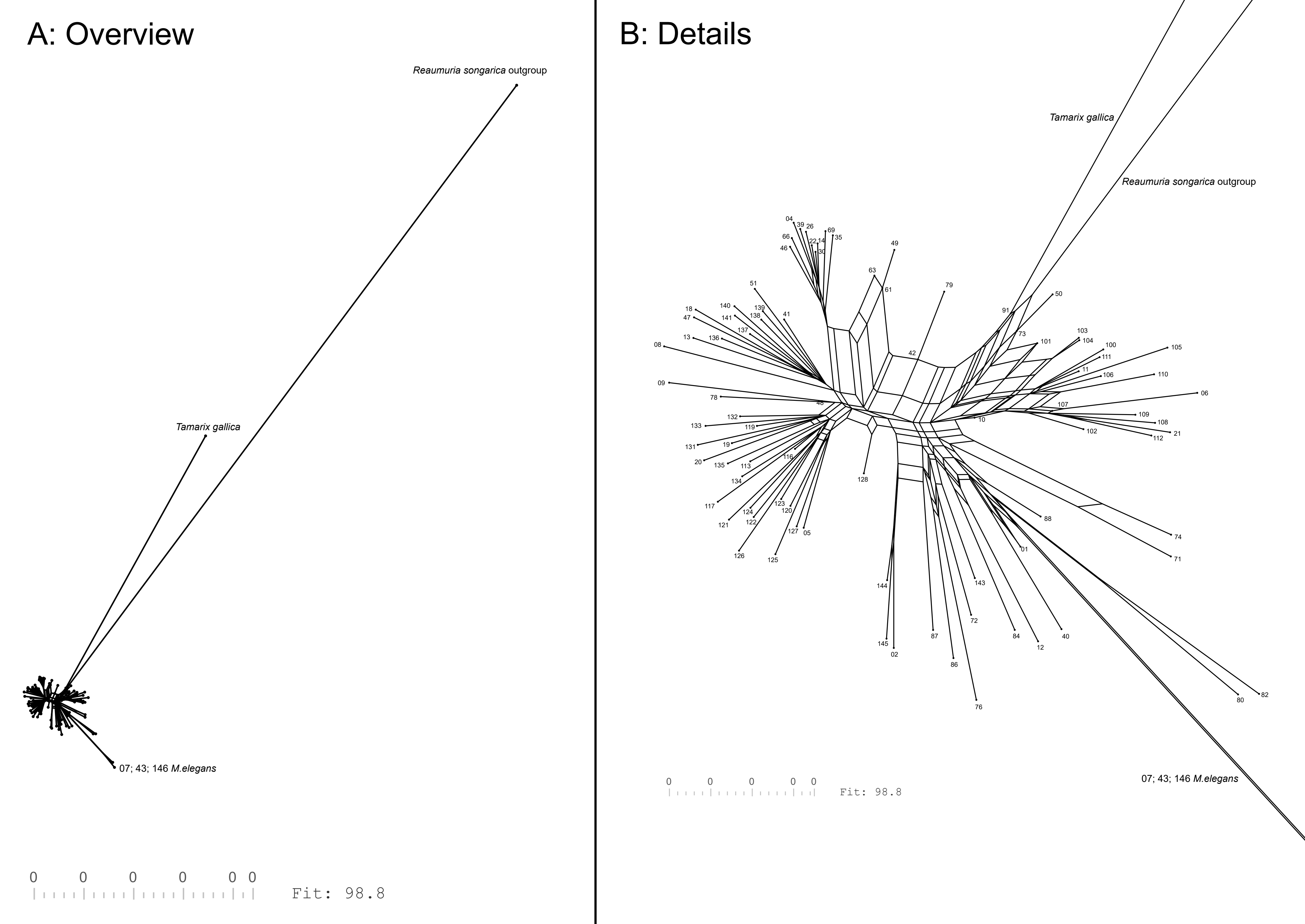

### Figure S5

#### Plastid Supermatrix Tree

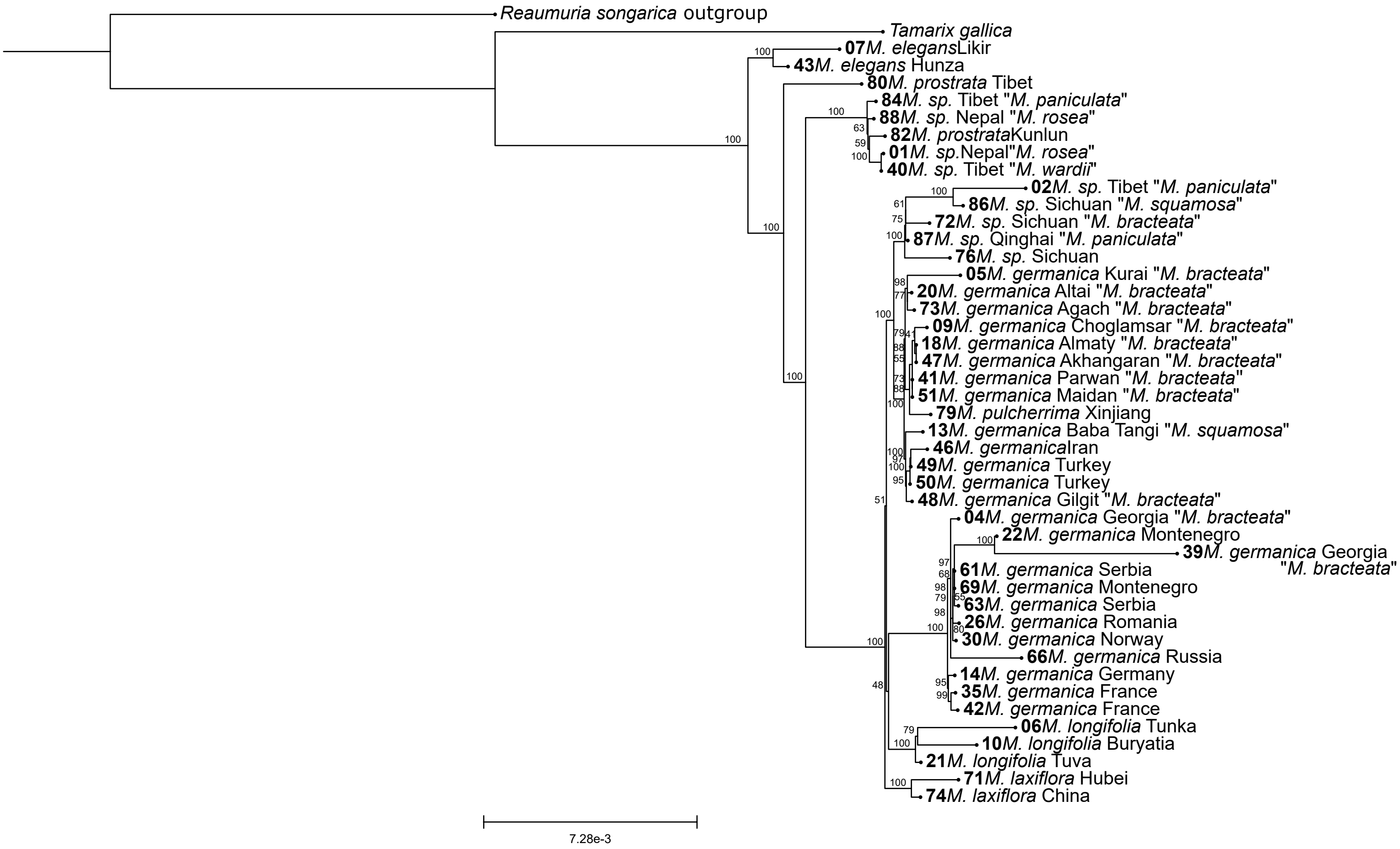
