## Supplementary material for "Towards an understanding of the taxonomy of the Eurasian montane riparian genus *Myricaria* (Tamaricaceae) through Angiosperms353 herbariomics": Table S1

### Table S1: Overview of sampled *Myricaria* specimens

| Specimen identifiers | | | | | | Location Data from herbarium label (coordinates where available) | | | NCBI BioSample | Notes |
|---|---|---|---|---|---|---|---|---|---|---|
| **Specimen number** | **New diagnosed species name** | **Collector or citation** | **Collection number** | **Collection date (DD.MM. YYYY)** | **Herbarium Code** | **Location Description** | **N Co-ordinate** | **E Co-ordinate** | **NCBI BioSample Accession number** | |
| 01 | *Myricaria* sp. | B. Dickoré | 11449 | 20.08.1994 | M | China: Tibet; Gyala Peri-N Glacier W (High Camp 15'), 3720m, subalp. Rhodedendron-Salix dwarf-scrub, lateral moraine | 29.9 | 94.88 | SAMN46873447 | |
| 02 | *Myricaria* sp. | D.E. Boufford, B. Bartholomew, D.A. Eaton, X.H. Li, R.H. Ree, B. Xu, J.P. Yue, J.W. Zhang, X.X. Zhu | 41694 | 02.08.2009 | M | China: Tibet; Gongjue Xian: Just N of the city of Gongjue on highway S501 (road to highway 314, and to Toba and Jiangda (Gyamda)). 3620m. Disturbed gravel borrow pit adjacent to highway. Disturbed gravel area adjacent to road. | 30.88 | 98.26 | SAMN46873448 | |
| 04 | *M. germanica* (L) Desv. | A. Gröger, W. Lobin, C. Lobin, Z. Schewardnadse | 1716 | 07.08.2009 | M | Georgia: Svaneti; zwischen Mestia und Ushguli; 1500m | 43.05 | 42.81 | SAMN46873449 | |
| 05 | *M. germanica* (L) Desv. | E. Bobrov | 5480 | 12.07.1971 | M | Russia: Kurai; Altai occidentalis, via Czujski tract dicta, Kuraj, in lapidosis fl. Kurajka | | | SAMN46873450 | |
| 06 | *M. longifolia* (Willd.) Ehrenb. | L. Malyschev, Y Petroczenko, S. Andrulaytis | 5482 | 15.06.1968 | M | Russia: Sayan Mountains; Montes Sajanenses orientales, vallis Tunkensis. prope pag. Tunka in arena mobili. | | | SAMN46873451 | |
| 07 | *M. elegans* Royle | H. Hartmann | 3127 | 20.07.1987 | M | India: Himachal Pradesh; am Bachlauf unterhalb Yangtang (zwischen Likir und Nurka) ca. 3550 | | | SAMN46873452 | |
| 08 | *Myricaria* sp. | B. Dickoré | 19452 | 04.10.2004 | M | India: Uttaranchal; Kumaon, Pithoragarh Dt., Gori Valley, Milam - Milam Glacier, 3520m | 30.45 | 80.12 | SAMN46873453 | |
| 09 | *Myricaria* sp. | H. Hartmann | 4191 | 26.07.1992 | M | India: Jammu and Kashmir; am Bewässerungsgraben östlich Choglamsar (südlich Leh) | | | SAMN46873454 | |
| 10 | *M. longifolia* (Willd.) Ehrenb. | L. Nikolaeva, A. Tkaczenko | 5481 | 28.08.1968 | M | Russia: Transbaikal; resp. aut. Burjatica, distr. Barguzin, prope pag. Pervomajsk (olim Schamanka), in fructicetis in valle fl. Schamanka. | | | SAMN46873455 | |
| 11 | *M. longifolia* (Willd.) Ehrenb. | V. Khanminczun | 5690 | 05.06.1972 | M | Russia: Tuva; distr. Tess-Khem, jugum Tannu-Ola orientalis, declivium australe, 1200 m.s.m. | | | SAMN46873456 | |
| 12 | *Myricaria* sp. | B. Dickoré | 5261 | 14.09.1989 | M | China: Tibet; Namchabarwar NW slope S above Nam La Tso - Nam La Pass 4550m | 29.58 | 95.02 | SAMN46873457 | |
| 13 | *M. germanica* (L) Desv. | H. Huss | 23 C | 26.07.1975 | M | Afghanistan: Wakhan; Baba Tangi, Im Schotter am Ufer eines Gletchscherbachs. 3000m. | | | SAMN46873458 | |
| 14 | *M. germanica* (L) Desv. | H. Krause | 7 | 12.07.2020 | M ( living specimen) | Germany: Bavaria; Munich Botanical Garden. Planted in the species conservation bed, A4. Bush, with white flowers. Original specimen collected from the wild in the Allgäu. | 47.65 | 10.8 | SAMN46873459 | |
| 18 | *M. germanica* (L) Desv. | M. Chytrý, M. Koci | 2014 no. 525 duplum | 07.07.2014 | BRNU | Kazakhstan: Almaty; distr. Karasai. Kaskelen, Kaskelen stream valley near the southern edge of the town. 1106 m.a.s.l. | 48.14 | 76.61 | SAMN46873460 | |
| 19 | *M. germanica* (L) Desv. | M. Hajek, P. Hajkova | 2005/81 | 05.08.2005 | BRNU | Russia: Altai; Altai Mts., Kosh-Agach distr., Kurai, Northern Chuya Ridge, Bolshoi Akturu river valley 18km SSW of the village; 2063 m.a.s.l. | 50.1 | 87.79 | SAMN46873461 | |
| 20 | *M. germanica* (L) Desv. | P. Lustyk | 2005/1283 | 05.08.2005 | BRNU | Russia: Altai; Altai Mts., Kosh-Agach distr., Kurai: Bolshoi Akturu river valley 18 km SSW of the vilalge; 2094 m.a. Hemiboreal forest with Pinus Sibirica on a steep SE facing slope | 50.09 | 87.79 | SAMN46873462 | |
| 21 | *M. longifolia* (Willd.) Ehrenb. | J. Danihelka | 2004/0256 | 06.08.2004 | BRNU | Russia: Tuva; Tuva Basin, Khemchik River Valley. Left bank in the floodplain of the Khemchik River 4km SSW of Sug-Agsy Town; 720 m.a.s.l. | 51.37 | 91.25 | SAMN46873463 | |
| 22 | *M. germanica* (L) Desv. | V. Kalníková, T. Peterka, S. Palpurina, Z. Pleskova | VK14/098 | 29.07.2014 | BRNU | Montenegro: Andrijevica; obstina Andrijevica. Andrijevica: Lim River, gravel bar, left bank, 788 m.s.m. | 42.7 | 19.84 | SAMN46873464 | |
| 26 | *M. germanica* (L) Desv. | V. Grulich et al. | VG10/157 | 16.06.2010 | BRNU | Romania: Alba; distr. Abrud. Posaga de Jos: Cheile Posegii, in dumetis secundum rivulum 4,3 km sept.-occ a pago versus. | | | SAMN46873465 | |
| 30 | *M. germanica* (L) Desv. | V. Kalníková, T. Peterka, T. Ruzickova, O. Knapek | VK14/007 | 31.08.2014 | BRNU | Norway: Sor-Trondelag; Storas: Orkla River, gravel bar, island. S. m. 130m. | 63.05 | 9.68 | SAMN46873466 | |
| 35 | *M. germanica* (L) Desv. | A. Charpin, W. Greuter, O. Monthoux | s.n. | 11.05.1970 | M | France: Laragne; secus amnem "Veragne" inter pagos "Laragne" et "Lazer" provinciae Alpinum superiorum 600m supra maris aequor. In ripis sabulosis. | | | SAMN46873467 | |
| 39 | *M. germanica* (L) Desv. | L. Meierott | Georg07/65 | 14.08.2007 | M | Georgia: Mtskheta-Mtianeti; Kaukasus, 2km s Kazbegi, felsige Hänge, ca 1600, | 42.38 | 44.38 | SAMN46873468 | |
| 40 | *Myricaria* sp. | B. Dickoré | 5810 | 28.09.1989 | M | China: Tibet; Tsangpo tributary, Nijiang/Lower Gyamda Chu, Bayi, SE of town. Lower mont., *Myricaria* shrub, moist to dry open riverside gravel and sand, ditches | 29.67 | 94.37 | SAMN46873469 | |
| 41 | *M. germanica* (L) Desv. | O. Anders | 9937 | 26.05.1973 | M | Afghanistan: Parwan; Panjir Tal, Sandbank oberhalb Kharu, 2150 m | | | SAMN46873470 | |
| 42 | *M. germanica* (L) Desv. | E. Berger | 4575 | 13.04.1973 | M | France: Moustiers Sainte-Marie; im Bett des Verdon, östlich des Pont d'Aiguines 510m | | | SAMN46873471 | |

| Specimen identifiers | | | | | | Location Data from herbarium label (coordinates where available) | | | NCBI BioSample | Notes |
|---|---|---|---|---|---|---|---|---|---|---|
| Specimen number | New diagnosed species name | Collector or citation | Collection number | Collection date (DD.MM. YYYY) | Herbarium Code | Location Description | N Co-ordinate | E Co-ordinate | NCBI BioSample Accession number | |
| 01 | *Myricaria* sp. | B. Dickoré | 11449 | 20.08.1994 | M | China: Tibet; Gyala Peri-N Glacier W (High Camp 15'), 3720m, subalp. Rhodedendron-Salix dwarf-scrub, lateral moraine | 29.9 | 94.88 | SAMN46873447 | |
| 43 | *M. elegans* Royle | F. Gartner | s.n. | 28.07.1974 | M | Pakistan: Karakorum; Königreich Nagar, Hunza-Tal, Pisan, Almwiese, Moränengeröll, 2600m | | | SAMN46873472 | |
| 46 | *M. germanica* (L) Desv. | K.H. Rechinger | 4/270 | 06.06.1971 | M | Iran: Azerbaijan Province; Prov. Azerbaijan orient.: In jugo inter Marand et Sufian, 1600-1750m | | | SAMN46873473 | |
| 47 | *Myricaria* sp. | V. Vašák | s.n. | 06.07.1973 | M | Uzbekistan: Tian Shan; Asia centralis, Tian-shan, apud flumen Akhangaran in vicinitate pagi Karakhtai. Alt. 600 m.s.m. | | | SAMN46873474 | |
| 48 | *Myricaria* sp. | S.M.A. Kazmi | 5684 | 02.07.1974 | M | Pakistan: Gilgit; Gopis & surrounding areas | | | SAMN46873475 | |
| 49 | *M. germanica* (L) Desv. | K.P. Buttler, W. Schmidt, I. Turgut, S. Yüce | 21760 | 24.08.1976 | M | Turkey: Ağrı; Ala dag S Diyadin, Bachtal 0-500 m unterhalb Güdük köyü, 2030m Bachalluvionen | | | SAMN46873476 | |
| 50 | *M. germanica* (L) Desv. | M. Nydegger | 46664 | 09.07.1992 | M | Turkey: Oltu; A8 Erzurum: Oltu-Narman skm s Oltu | | | SAMN46873477 | |
| 51 | *Myricaria* sp. | D. Podlech | 19489 | 24.08.1970 | M | Afghanistan: Maidan; Darrah-i-Sanglakh (Seitental des Maidan-Tales) bei Dahane Urmun, 2650m | | | SAMN46873478 | |
| 61 | *M. germanica* (L) Desv. | "Vuk.S., Zlat.B, Rand.V." | 5694 | 08.06.1997 | BEOU | Serbia: Knjazevac-Kalna; | | | SAMN46873479 | |
| 63 | *M. germanica* (L) Desv. | V. Stevanovic, M. Niketic, G. Tomovic | 12781 | 17.07.1998 | BEOU | Serbia: Knjazevac; (istoncna), Knjazevac, Kalna obale reke | | | SAMN46873480 | |
| 66 | *M. germanica* (L) Desv. | Đ. Milanovic | 39334 | 17.07.2007 | BEOU | Russia: Ullu Tau; Kavkaz, Ullu Tau dolina uz reku Adir su aluvijalni nanosi (on the Georgian border) | | | SAMN46873481 | |
| 69 | *M. germanica* (L) Desv. | D. Lakusic, N. Kuzmanovic, I. Jankovic | 46516 | 30.07.2016 | BEOU | Montenegro: Berane; Lim, Berane, Buce, 670 alt, gravel river banks (Salicion eleagni), limestone | 42.82 | 19.87 | SAMN46873482 | |
| 71 | *M. laxiflora* (Franch.) P.Y.Zhang & Y.J.Zhang | D. Wang | 952 | 28.07.2016 | BEOU | China: Hubei; Zhijiang City, Dongshi Town. | | | SAMN46873483 | |
| 72 | *Myricaria* sp. | X. Li | 1271 | 26.05.2012 | TAIF | China: Sichuan; Jiulong County, Mt. Jichou | | | SAMN46873484 | |
| 73 | *M. germanica* (L) Desv. | M. Wakabayashi, M. Amano, N. V. Friesen, Y. Kadota, A. A. Krasnikov, I. M. Krasnoborov, M. N. Lomonosova, M. Minaki, M. M. Tamura | 9280528 | 27.07.1992 | TAIF | Russia: Altai; Kosh-Agach District, the North Chuya Range, basin on the Akturu River alt. 2150m | | | SAMN46873485 | |
| 74 | *M. laxiflora* (Franch.) P.Y.Zhang & Y.J.Zhang | No data recorded | No data recorded | No data recorded | HIB | China: | | | SAMN46873486 | |
| 76 | *Myricaria* sp. | H. Smith | 11085 | 16.08.1934 | UPS | China: Sichuan; Sikang: Kangting (Tachienlu) distr.: Chungo valley: Chungo. Ad rivum. 3000m | | | SAMN46873487 | |
| 78 | *M. germanica* (L) Desv. | "J. Homeier & Kazakhstan Exkursion 2001" | 824 | 12.08.2001 | GOET | Kazakhstan: Almaty; valley of the Talghar-river; | | | SAMN46873488 | |
| 79 | *M. pulcherrima* Batalin | B. Dickoré | 0126 | 17.08.1986 | GOET | China: Xinjiang; SW-Kashgaria (SW Takla Makan desert). Yechen, southeastern margin of the town. On damp loess, close to an irrigation pond | 37.88 | 77.42 | SAMN46873489 | |
| 80 | *M. prostrata* Hook.f. & Thomson | G. Miehe, S. Miehe | NX05-094-01 | 13.08.2005 | MB | China: Tibet; West of Lake Xiketang, 4630 m | 32.1 | 90.68 | SAMN46873490 | |
| 82 | *M. prostrata* Hook.f. & Thomson | G. Miehe, S. Miehe | 8250 | 06.07.1992 | MB | China: Xinjiang; Kunlun, between Dahoongliutan and the Karakax-Aksai Chin watershed. Open subalpine/alpine steppe patches (Carex moorcroftii) and cushion communities flushes | 35.8 | 79.35 | SAMN46873491 | |
| 84 | *Myricaria* sp. | D. E. Boufford, J. H. Chen, S. L. Kelley, J. Li, R. H. Ree, H. Sun, J. P. Yue, Y. H. Zhang | 32248 | 13.08.2004 | A | China: Tibet; Riwoqe | 31.08 | 96.94 | SAMN46873492 | |
| 86 | *Myricaria* sp. | D. E. Boufford, K. Fujikawa, S. L. Kelley, R. H. Ree, Bo Xu, J. W. Zhang, T. C. Zhang, W. D. Zhu | 38168 | 25.07.2007 | A | China: Sichuan; Dawu | 30.47 | 101.66 | SAMN46873493 | |
| 87 | *Myricaria* sp. | T. N. Ho, B. Bartholomew, M. F. Watson, M. G. Gilbert | 2786 | 01.09.1996 | GH | China: Qinghai; Nangqen, along W side of the Za Qu (upper Mekong), on road between Nangqen and Yushu | 32.28 | 96.47 | SAMN46873494 | |
| 88 | *Myricaria* sp. | Y. Iokawa, T. Miyazaki, K. Yonekura, Y. Ibaragi, R. K. Uprety | 20310001 | 26.06.2003 | A | Nepal: Gandaki Zone; Mustang, Dhawalagiri Zone | 28.8 | 83.75 | SAMN46873495 | |

| Specimen identifiers | | | | | | Location Data from herbarium label (coordinates where available) | | | NCBI BioSample | Notes |
|---|---|---|---|---|---|---|---|---|---|---|
| Specimen number | New diagnosed species name | Collector or citation | Collection number | Collection date (DD.MM. YYYY) | Herbarium Code | Location Description | N Co-ordinate | E Co-ordinate | NCBI BioSample Accession number | |
| 01 | *Myricaria* sp. | B. Dickoré | 11449 | 20.08.1994 | M | China: Tibet; Gyala Peri-N Glacier W (High Camp 15'), 3720m, subalp. Rhodedendron-Salix dwarf-scrub, lateral moraine | 29.9 | 94.88 | SAMN46873447 | |
| 90 | *M. platyphylla* Maxim. | N.M. Przewalski | 39 | 1872 | P | China: Inner Mongolia; "Mongolia occidentalis mont. Alaschan" - in modern-day China, Inner Mongolia | | | SAMN46873496 | Not included in analysis due to poor gene extraction |
| 91 | *M. platyphylla* Maxim. | N.M. Przewalski | s.n. | 1885 | P | China: Taklamakan Desert; "Mongolia occidentalis Chotan" - in modern-day China, Taklamakan desert | | | SAMN46873497 | Not included in analysis due to poor gene extraction |
| 100 | *M. longifolia* (Willd.) Ehrenb. | E. M. Lyakh | 1 | 29.07.1987 | NSK | Russia: Tuva; river Kara Sug; | 51.61434 | 90.074897 | SAMN46873498 | |
| 101 | *M. longifolia* (Willd.) Ehrenb. | D. N. Shaulo | 2 | 27.07.1987 | NSK | Russia: Tuva; river Ak Sug; | 51.57803 | 90.098132 | SAMN46873499 | |
| 102 | *M. longifolia* (Willd.) Ehrenb. | E. M. Lyakh | 3 | 29.07.1987 | NSK | Russia: Tuva; river Hemchik; | 51.15448 | 90.576568 | SAMN46873500 | |
| 103 | *M. longifolia* (Willd.) Ehrenb. | E. M. Lyakh | 5 | 05.08.1995 | NSK | Russia: Tuva; river Erzin near river Ular; | 50.38993 | 95.603371 | SAMN46873501 | |
| 104 | *M. longifolia* (Willd.) Ehrenb. | E. M. Lyakh | 6 | 05.08.1994 | NSK | Russia: Tuva; river Tes-khem; | 49.9913 | 95.484254 | SAMN46873502 | |
| 105 | *M. longifolia* (Willd.) Ehrenb. | E. M. Lyakh | 7 | 16.07.1994 | NSK | Russia: Tuva; river Shivilig-khem; | 50.72621 | 94.557988 | SAMN46873503 | |
| 106 | *M. longifolia* (Willd.) Ehrenb. | E. M. Lyakh | 9 | 18.08.1995 | NSK | Russia: Tuva; river Hondergey, Chadan; | 51.15725 | 91.569844 | SAMN46873504 | |
| 107 | *M. longifolia* (Willd.) Ehrenb. | E. M. Lyakh | 10 | 28.07.2003 | NSK | Russia: Buryatia; river Shamanka Kurumkanskij reg. | 54.5054 | 110.4856 | SAMN46873505 | |
| 108 | *M. longifolia* (Willd.) Ehrenb. | E. M. Lyakh | 11' | 22.08.2004 | NSK | Russia: Buryatia; river Zun-Murin | 51.72867 | 102.86257 | SAMN46873506 | |
| 109 | *M. longifolia* (Willd.) Ehrenb. | E. M. Lyakh | 13 | 18.07.1995 | NSK | Russia: Buryatia; river Djida | 50.60274 | 105.28975 | SAMN46873507 | |
| 110 | *M. longifolia* (Willd.) Ehrenb. | E. M. Lyakh | 15' | 23.08.2004 | NSK | Russia: Buryatia; river Irkut | 51.67316 | 101.00423 | SAMN46873508 | |
| 111 | *M. longifolia* (Willd.) Ehrenb. | E. M. Lyakh | 18 | 26.07.1987 | NSK | Russia: Khakassia; river Bol'shoy On. | 52.06893 | 89.758116 | SAMN46873509 | |
| 112 | *M. longifolia* (Willd.) Ehrenb. | E. M. Lyakh | 19' | 10.08.2016 | NSK | Russia: Altai; river Yustyt, Aktal village. | 49.94462 | 88.817468 | SAMN46873510 | |
| 113 | *M. germanica* (L) Desv. | E. M. Lyakh | 20 | 15.08.1987 | NS | Russia: Altai; river Chuuya | 50.32322 | 87.526693 | SAMN46873511 | |
| 116 | *M. germanica* (L) Desv. | E. M. Lyakh | 24 | 30.06.1990 | NSK | Russia: Altai; river Sema | 51.57741 | 85.578622 | SAMN46873512 | |
| 117 | *M. germanica* (L) Desv. | E. M. Lyakh | 25 | 30.06.1990 | NSK | Russia: Altai; river Cherginka | 51.56545 | 85.540707 | SAMN46873513 | |
| 119 | *M. germanica* (L) Desv. | E. M. Lyakh | 28 | 03.09.1993 | NSK | Russia: Altai; river Katanda | 50.14325 | 86.17849 | SAMN46873514 | |
| 120 | *M. germanica* (L) Desv. | E. M. Lyakh | 29 | 04.09.1993 | NSK | Russia: Altai; river Charysh | 50.85486 | 84.879711 | SAMN46873515 | |
| 121 | *M. germanica* (L) Desv. | E. M. Lyakh | 31 | 25.07.2022 | NSK | Russia: Altai; river Chulyshman Ulaganskij reg | 51.3437 | 87.7448 | SAMN46873516 | |
| 122 | *M. germanica* (L) Desv. | E. M. Lyakh | 33 | 11.08.2016 | NSK | Russia: Altai; river Chibitka | 50.31812 | 87.542547 | SAMN46873517 | |
| 123 | *M. germanica* (L) Desv. | E. M. Lyakh | 44 | 17.08.1987 | NS | Russia: Altai; river Bol'shoj Yaloman | 50.52083 | 86.562011 | SAMN46873518 | |

| Specimen identifiers | | | | | | Location Data from herbarium label (coordinates where available) | | | NCBI BioSample | Notes |
|---|---|---|---|---|---|---|---|---|---|---|
| **Specimen number** | **New diagnosed species name** | **Collector or citation** | **Collection number** | **Collection date (DD.MM. YYYY)** | **Herbarium Code** | **Location Description** | **N Co-ordinate** | **E Co-ordinate** | **NCBI BioSample Accession number** | |
| 01 | *Myricaria* sp. | B. Dickoré | 11449 | 20.08.1994 | M | China: Tibet; Gyala Peri-N Glacier W (High Camp 15'), 3720m, subalp. Rhodedendron-Salix dwarf-scrub, lateral moraine | 29.9 | 94.88 | SAMN46873447 | |
| 124 | *M. germanica* (L) Desv. | E. M. Lyakh | 34 | 28.07.2022 | NSK | Russia: Altai; river Okporok, Teletskoye lake | 51.45597 | 87.38103 | SAMN46873519 | |
| 125 | *M. germanica* (L) Desv. | E. M. Lyakh | 36 | 22.07.2022 | NSK | Russia: Altai; river Chechenek, Teletskoye lake | 51.46054 | 87.36106 | SAMN46873520 | |
| 126 | *M. germanica* (L) Desv. | E. M. Lyakh | 37 | 24.07.2022 | NSK | Russia: Altai; river Kyga, Teletskoye lake | 51.21229 | 87.50569 | SAMN46873521 | |
| 127 | *M. germanica* (L) Desv. | E. M. Lyakh | 39 | 27.07.1987 | NSK | Russia: Khakassia; river Ona near the mouth of the river Bol'shoj On | 52.0712 | 89.751151 | SAMN46873522 | |
| 128 | *M. germanica* (L) Desv. | E. M. Lyakh | 46 | 31.05.2002 | NSK | Kazakhstan: Abai; Urdzharskij, river Karabuta | 46.6708 | 82.6611 | SAMN46873523 | |
| 129 | *M. germanica* (L) Desv. | E. M. Lyakh | 43 | 01.06.2002 | NSK | Kazakhstan: Abai; Urdzharskij; river Karabuta | 46.7391 | 82.2773 | SAMN46873524 | |
| 131 | *M. germanica* (L) Desv. | E. M. Lyakh | 41 | 24.05.2016 | NSK | Kazakhstan: Jetisu; river Aksu | 45.63834 | 79.45404 | SAMN46873525 | |
| 132 | *M. germanica* (L) Desv. | E. M. Lyakh | 38 | 25.08.2010 | NSK | Russia: Altai; river Terekta | 50.18491 | 85.52862 | SAMN46873526 | |
| 133 | *M. germanica* (L) Desv. | E. M. Lyakh | 42 | 25.05.2016 | NSK | Kazakhstan: Almaty; river Charyn | 43.76511 | 79.431079 | SAMN46873527 | |
| 134 | *M. germanica* (L) Desv. | E. M. Lyakh | 32 | 12.08.2016 | NSK | Russia: Altai; river Ursul | 50.84154 | 85.69485 | SAMN46873528 | |
| 135 | *M. germanica* (L) Desv. | E. M. Lyakh | 30' | 18.08.2005 | NSK | Russia: Altai; river Chuuya | 50.07351 | 87.767477 | SAMN46873529 | |
| 136 | *M. germanica* (L) Desv. | E. M. Lyakh | 49 | 17.08.2013 | NSK | Tajikistan: Hissar Valley; | 38.9427 | 69.3089 | SAMN46873530 | |
| 137 | *M. germanica* (L) Desv. | E. M. Lyakh | 50 | 21.08.2013 | NSK | Tajikistan: Gorno-Badakhshan;. river Pyanj | 37.4736 | 71.5234 | SAMN46873531 | |
| 138 | *M. germanica* (L) Desv. | E. M. Lyakh | 51 | 21.08.2013 | NSK | Tajikistan: Gorno-Badakhshan; kishlak Avj (village) | 36.9245 | 71.507 | SAMN46873532 | |
| 139 | *M. germanica* (L) Desv. | E. M. Lyakh | 53 | 22.08.2013 | NSK | Tajikistan: Gorno-Badakhshan; river Gunt | 37.7016 | 72.3768 | SAMN46873533 | |
| 140 | *M. germanica* (L) Desv. | E. M. Lyakh | 54 | 23.08.2013 | NSK | Tajikistan: Gorno-Badakhshan; river Sokhcharv | 37.7219 | 71.5533 | SAMN46873534 | |
| 141 | *M. germanica* (L) Desv. | E. M. Lyakh | 55 | 23.08.2013 | NSK | Tajikistan: Gorno-Badakhshan; river Pasthur | 37.6626 | 71.5993 | SAMN46873535 | |
| 143 | *Myricaria* sp. | E. M. Lyakh | 58 | 15.07.2000 | NSK | China: Yunnan; | 27.1609 | 100.0565 | SAMN46873536 | |
| 144 | *Myricaria* sp. | E. M. Lyakh | 59 | 13.07.2000 | NSK | China: Yunnan; | 26.9221 | 100.3003 | SAMN46873537 | |
| 145 | *Myricaria* sp. | E. M. Lyakh | 60 | 13.07.2000 | NSK | China: Yunnan; | 26.9152 | 100.0344 | SAMN46873538 | |
| 146 | *M. elegans* Royle | E. M. Lyakh | 52 | 21.08.2013 | NSK | Tajikistan: Gorno-badakhshan; kishlak Avj (village) | 36.9245 | 71.507 | SAMN46873539 | |
