## Supplementary material for "Towards an understanding of the taxonomy of the Eurasian montane riparian genus *Myricaria* (Tamaricaceae) through Angiosperms353 herbariomics": Table S2

Table S2: Morphological overview of specimens from clades 2, 3, 4, and 6

| Specimen identifier | | Observations (where available) | | | | | | | | | | | | | | | | | |
|---|---|---|---|---|---|---|---|---|---|---|---|---|---|---|---|---|---|---|---|
| **Specimen number** | **Clade nuclear ortholog species tree** | **a** | **b** | **c** | **d** | **e** | **f** | **g** | **h** | **i** | **j** | **k** | **l** | **m** | **n** | **o** | **p** | **q** | **r** |
| 01 | 3 | erect | 1 | 1 | 1 | dense | 0 | 3 | 1 | NA | 4 | 1 | 1 | 2 | 0.5 | 3 | 1.5 | purple to white | |
| 02 | 6 | erect | 0 | 0 | 1 | dense | 0 | 1 | <0.5 | 2/3 united at base | 4.5 | 1 | 0 | 3 | 1 | 4.5 | 2 | pink | |
| 12 | 2 | prostrate | 0 | 0 | 1 | dense | 0 | 2 | 0.5 | feature not available - wrong developmental stage | 5 | 2 | 0 | 2.5 | 0.5 | 3.5 | 1.5 | white or pink | |
| 40 | 3 | erect | 1 | 1 | 1 | dense | 0 | 2 | <0.5 | | 2 | 0.5 | 1 | 1.5 | 0.5 | 2.5 | 1 | purple to white | |
| 71 | 5 | erect | 0 | 0 | 1 | dense | 0 | 2.5 | <0.5 | | 5 | 1 | 0 | 4 | 1 | 5 | 2 | white | |
| 72 | 4 | erect | 0 | 0 | 1 | dense | 0 | 1.5 | <0.5 | | unsure | unsure | 1 | 2 | 1.5 | 3 | 1.5 | pink | Papery sepals with dark centre |
| 74 | 5 | feature not available | NA | NA | NA | dense | 0 | 3 | 1 | NA | NA | NA | NA | NA | NA | NA | NA | NA | |
| 76 | 4 | erect | 1 | 0 | 0 | dense | 0 | 1.5 | <0.5 | NA - wrong developmental stage | 1,5 (early development) | 0,5 (early development) | 1 | buds not open | buds not open | buds not open | buds not open | purple to white | |
| 84 | 2 | erect | 1 | 0 | 1 | dense | 0 | 1.5 | <0.5 | feature not available | 5 | 1 | 1 | 2.5 | 1 | 3 | 1.5 | white or pink | Papery sepals with dark centre, slightly purple |

| Specimen number | Clade nuclear ortholog species tree | a | b | c | d | e | f | g | h | i | j | k | l | m | n | o | p | q | r |
|---|---|---|---|---|---|---|---|---|---|---|---|---|---|---|---|---|---|---|---|
| 86 | 4 | erect | 1 | 0 | 1 | dense | 0 | 1.5 | <0.5 | 2/3 united at base | 4 | 1 | 1 | 2 | <0.5 | 3 | 1.5 | purple to white | Papery sepals with dark centre |
| 87 | 6 | erect | 0 | 0 | 1 | dense | 0 | 3 | 0.5 | | 3 | 2 | 0 | 3 | 1.5 | 5 | 2.5 | pink | |
| 88 | 3 | prostrate | 1 | 1 | 1 | dense | 0 | 2.5 | 0.5 | 2/3 united at base | 4 | 1.5 | none visible | 2 | 0.5 | 3.5 | 1 | white | Papery sepals with dark centre |
| 143 | 4 | erect | 1 | 0 | 1 | dense | 0 | 1.5 | <0.5 | | | | 1 | 2 | 1 | 3 | 2.5 | purple | |
| 144 | 6 | erect | 0 | 0 | 1 | dense | 0 | 1 | <0.5 | | | | 0 | unsure | unsure | 4 | 2 | unsure | |
| 145 | 6 | erect | 0 | 0 | 1 | dense | 0 | 1 | <0.5 | | | | 0 | unsure | unsure | 4 | 2 | unsure | |

**Key:**

a) Habit; b) Terminal single flowers; c) Few-flowered racemes; d) Many-flowered racemes; e) Leaves lax or dense; f) Leaves cordate; g) Leaf length (mm); h) Leaf width (mm); i) Filaments united; j) Flower bract length (mm); k) Bract width (mm); l) Imbricate scales at base; m) Sepal length (mm); n) Sepal width (mm); o) Petal length (mm); p) Petal width (mm); q) Petal colour; r) Notes.
